# The proteomic landscape of centromeric chromatin reveals an essential role for the Ctf19^CCAN^ complex in meiotic kinetochore assembly

**DOI:** 10.1101/2020.06.23.167395

**Authors:** Weronika E. Borek, Nadine Vincenten, Eris Duro, Vasso Makrantoni, Christos Spanos, Krishna K. Sarangapani, Flavia de Lima Alves, David A. Kelly, Charles L. Asbury, Juri Rappsilber, Adele L. Marston

## Abstract

Kinetochores direct chromosome segregation in mitosis and meiosis. Faithful gamete formation through meiosis requires that kinetochores take on new functions that impact homolog pairing, recombination and the orientation of kinetochore attachment to microtubules in meiosis I. Using an unbiased proteomics pipeline, we determined the composition of centromeric chromatin and kinetochores at distinct cell-cycle stages, revealing extensive reorganisation of kinetochores during meiosis. The data uncover a network of meiotic chromosome axis and recombination proteins that replace the microtubule-binding outer kinetochore sub-complexes during meiotic prophase. We show that this kinetochore remodelling in meiosis requires the Ctf19c^CCAN^ inner kinetochore complex. Through functional analyses, we identify a Ctf19c^CCAN^-dependent kinetochore assembly pathway that is dispensable for mitotic growth, but becomes critical upon meiotic entry. Therefore, extensive kinetochore remodelling and a distinct assembly pathway direct the specialization of meiotic kinetochores for successful gametogenesis.

## Introduction

The kinetochore is a multi-molecular machine that links centromeric nucleosomes to microtubules for chromosome segregation (Hinshaw and Harrison, 2018; Joglekar and Kukreja, 2017). Kinetochores need to be modified for meiosis, the specialised cell division that generates gametes. Meiosis reduces ploidy by half through two consecutive divisions that follow a single round of DNA replication. In meiotic prophase, replicated homologous chromosomes pair, synapse and recombine to generate crossovers that enable their accurate segregation in meiosis I (Gray and Cohen, 2016). While homologous chromosomes segregate to opposite poles, monooriented sister chromatids co-segregate because sister kinetochores are attached to microtubules from the same pole (Duro and Marston, 2015). Loss of sister chromatid cohesion on chromosome arms triggers homolog segregation in meiosis I, while retention of centromeric cohesin allows accurate segregation of sister chromatids in meiosis II (Marston and Amon, 2004). Kinetochores play multiple roles in these meiosis-specific processes throughout the meiotic divisions (Brar and Amon, 2009). Importantly, kinetochore defects have been implicated in age-related oocyte deterioration in humans (Patel et al., 2015; Zielinska et al., 2015, 2019) and proposed to contribute to the high degree of meiotic chromosome segregation errors, causing infertility, birth defects and miscarriages (Gruhn et al., 2019; Hassold and Hunt, 2001).

Core kinetochore factors show a high degree of conservation and henceforth, *Saccharomyces cerevisiae* protein names are used with the human homolog given in superscript. Kinetochore assembly is restricted to centromeres because of specific incorporation of the histone H3 variant CENP-A, which is positioned epigenetically in most organisms (Allshire and Karpen, 2008). However, in *S. cerevisiae*, sequence-specific binding of the Cbf3 complex (Cbf3c) enables formation of a single Cse4^CENP-A^-containing nucleosome to direct kinetochore assembly (Furuyama and Biggins, 2007; Lechner and Carbon, 1991). The Cse4^CENP-A^ nucleosome directly contacts Mif2^CENP-C^ and components of the inner kinetochore 13-subunit Ctf19 complex (Ctf19c, known as CCAN in humans) (Anedchenko et al., 2019; Carroll et al., 2009; Fischböck-Halwachs et al., 2019; Hinshaw and Harrison, 2019; Hornung et al., 2014; Weir et al., 2016; Xiao et al., 2017). Mif2^CENP-C^ and Ctf19c^CCAN^, while directly contacting each other, form separate links to the 4-subunit Mtw1 complex (Mtw1c^MIS12c^, also known as MIND), forming the structural core of the kinetochore (Dimitrova et al., 2016; Hornung et al., 2014; Killinger et al., 2020; Przewloka et al., 2011; Screpanti et al., 2011). The outer Spc105 and Ndc80 complexes (Spc105c^KNL1c^ and Ndc80c^NDC80c^, respectively) assemble onto Mtw1c^MIS12c^ to provide the microtubule binding interface, which is stabilised by the 10-component Dam1 complex (Dam1c) in *S. cerevisiae* or the structurally distinct Ska complex in humans (Van Hooff et al., 2017). A separate link from Ctf19c^CCAN^ to Ndc80c^NDC80c^ is provided by Cnn1^CENP-T^, though this is dispensable for viability in budding yeast (Lang et al., 2018; Pekgöz Altunkaya et al., 2016; Schleiffer et al., 2012).

How kinetochore assembly is regulated remains unclear; however, in budding and fission yeast, *Xenopus* egg extracts, chicken and human cells, Aurora B kinase facilitates kinetochore assembly by phosphorylating two conserved serines on the Dsn1^DSN1^ component of the Mtw1c^MIS12c^ (Akiyoshi et al., 2013a; Bonner et al., 2019; Hara et al., 2018; Kim and Yu, 2015; Rago et al., 2015; Zhou et al., 2017). This displaces an autoinhibitory fragment of Dsn1^DSN1^, exposing a binding site on Mtw1c^MIS12c^ for the inner kinetochore Mif2^CENP-C^ and, in yeast, Ame1^CENP-U^ proteins (Dimitrova et al., 2016; Petrovic et al., 2016). An autoinhibitory mechanism similarly prevents Mif2^CENP-C^ that is not bound to the centromeric nucleosome from binding to Mtw1c (Killinger et al., 2020). In yeast, Ipl1^AURORA B^ may further stabilise the kinetochore through phosphorylation of Cse4^CENP-A^ (Boeckmann et al., 2013). However, given that depletion of Aurora B does not cause a profound kinetochore assembly defect in multiple organisms, with the exception of *Xenopus* egg extracts, Aurora B cannot be critical for kinetochore assembly, suggesting the existence of additional mechanisms (Bonner et al., 2019; Emanuele et al., 2008; Haase et al., 2017).

In addition to their canonical role in making attachments to microtubules, kinetochores have additional functions that are critical for chromosome segregation. Kinetochores promote pericentromeric cohesin enrichment through cohesin loading onto centromeres (Eckert et al., 2007; Fernius and Marston, 2009; Fernius et al., 2013; Ng et al., 2009), shape pericentromere structure (Paldi et al., 2020), and monitor proper attachment of chromosomes to microtubules (Foley and Kapoor, 2013; Marston and Wassmann, 2017). They also adopt additional roles during meiosis, including non-homologous centromere coupling and repression of meiotic recombination in prophase (Kuhl and Vader, 2019; Kurdzo and Dawson, 2015; Vincenten et al., 2015). During meiotic prophase, the outer kinetochore (Ndc80c^NDC80c^ and Dam1c) is shed, which may facilitate kinetochore specialization in preparation for the meiotic divisions (Hayashi et al., 2006; Kim et al., 2013; Miller et al., 2012). Indeed, prior to metaphase I, factors important for sister kinetochore monoorientation, such as monopolin, Spo13^MEIKIN^ and Cdc5^Plk1^, are recruited to kinetochores (Duro and Marston, 2015; Galander et al., 2019; Kim et al., 2015). Kinetochores also direct the localised recruitment of Shugoshin-PP2A, setting up the protection mechanism that ensures the retention of pericentromeric cohesin until meiosis II (Yamagishi et al., 2010).

Many of these roles of the kinetochore require Ctf19c^CCAN^, but only the Ame1^CENP-U^-Okp1^CENP-Q^ heterodimer is essential for growth. Though viable, cells lacking other Ctf19c^CCAN^ subunits show chromosome segregation defects, which may, in part, be attributed to their role in loading cohesin (Eckert et al., 2007; Fernius and Marston, 2009; Ng et al., 2009). Cohesin loading occurs through the phosphorylation-dependent docking of the Scc2^NIPBL^-Scc4^MAU2^ complex onto an N-terminal extension of Ctf19^CENP-P^ (Hinshaw et al., 2015, 2017; Natsume et al., 2013). Ctf19^CENP-P^, through its C-terminal domain, also acts as an inner kinetochore receptor for the Ipl1^AURORA B^ kinase (Fischböck-Halwachs et al., 2019; García-Rodríguez et al., 2019). The Ctf19c^CCAN^ is further implicated in various meiosis-specific processes, including the non-homologous coupling of centromeres in early meiotic prophase, suppression of crossovers near centromeres during meiotic recombination, and the maintenance of pericentromeric cohesion until meiosis II (Marston et al., 2004; Vincenten et al., 2015).

Accumulating evidence supports the notion that kinetochore function, composition, and regulation are specialised during meiosis. Here, we use quantitative proteomics to determine the composition of the budding yeast kinetochore and centromere in meiotic prophase I, metaphase I and mitotically cycling cells, revealing the unique adaptations made for meiosis. Using a minichromosome purification proteomics pipeline and live-cell imaging, we demonstrate a central role for the Ctf19c^CCAN^ in meiotic kinetochore assembly, composition and function. These findings identify distinct requirements for kinetochore assembly and function in gametogenesis.

## RESULTS

### Chromatin, centromere, and kinetochore proteomes

To reveal the changes in centromeric chromatin composition that underlie its specialised functions during meiosis, we analysed the proteome of circular centromeric minichromosomes isolated from budding yeast cells at different cell cycle stages. We immunoprecipitated LacI-FLAG bound to *lacO* arrays on a circular minichromosome carrying the budding yeast centromere 3 (*CEN3*) sequence (Figure 1A; *CEN* chromatin) (Akiyoshi et al., 2009a; Unnikrishnan et al., 2012). To identify chromatin-associated proteins that require a functional centromere, in parallel, we analysed the proteome of a minichromosome that is identical except for two mutations within *CEN3* that abolish recruitment of the centromeric nucleosome and therefore prevent kinetochore assembly (Figure 1A; *CEN\** chromatin) (Akiyoshi et al., 2009a; Hegemann and Fleig, 1993; Meluh et al., 1998). Using label-free quantitative mass spectrometry (LFQMS), we compared the composition of *CEN* and *CEN\** chromatin in three conditions: mitotically cycling cells, cells arrested in meiotic prophase I (by deletion of *NDT80*, encoding a global meiotic transcription factor (Xu et al., 1995)) and cells arrested in meiotic metaphase I (by depletion of the APC/C activator, Cdc20 (Lee and Amon, 2003)).

**Figure 1.**
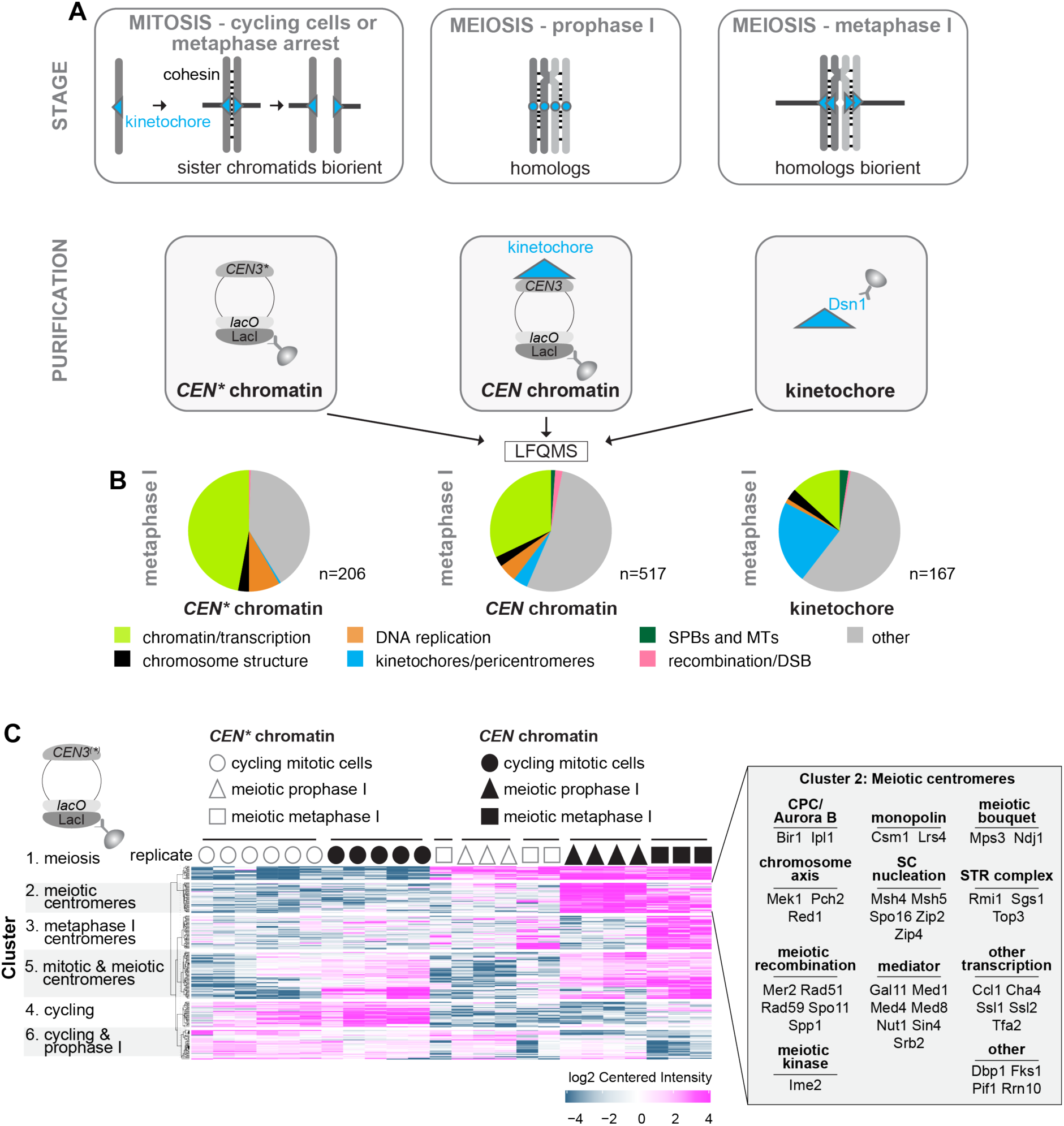
Quantitative label-free mass spectrometry (LFQMS) reveals the complexity of the centromere and kinetochore-associated proteomes. (A) Schematic representation of determined proteomes. *CEN* chromatin, *CEN\** chromatin and kinetochores were isolated from cycling cells, cells arrested in meiotic prophase I and meiotic metaphase I and subjected to LFQMS. (B) *CEN\** chromatin, *CEN* chromatin and kinetochores show respective increases and decreases in the fraction of enriched proteins that are associated with kinetochores or chromatin, respectively. Following immunoprecipitation of LacI-3FLAG (*CEN* chromatin and *CEN\** chromatin) and Dsn1-6His-3FLAG (kinetochores), proteins were quantified using LFQMS, and those enriched over respective negative controls with a cut-off of Log_2_(Fold Change) > 4 and p < 0.01 were categorised in the indicated groups. (C) Stage-specific functional groups of proteins were found to associate with *CEN* chromatin and *CEN\** chromatin. Clustering analysis of *CEN* chromatin and *CEN\** chromatin samples (k-means clustering). A cut-off of Log_2_(Fold Change) > 2 and p < 0.05 was used. Cluster 2 proteins are listed in the inset. See also Supplemental Table S1.

We also generated an orthogonal LFQMS dataset of the kinetochore by direct immunoprecipitation of FLAG-tagged core kinetochore protein, Dsn1^DSN1^ (Akiyoshi et al., 2010). Like the *CEN* and *CEN\** datasets, our kinetochore dataset represents cycling mitotic, meiotic prophase I, and meiotic metaphase I cells (Figure 1A). In addition, we determined the kinetochore proteome of cells arrested in mitotic metaphase by treatment with the microtubule-depolymerizing drug, benomyl.

In all three datasets (*CEN\** chromatin, *CEN* chromatin and kinetochore), chromatin-associated proteins were highly enriched over a no-tag control (Figure 1B). In metaphase I, *CEN* chromatin contained the largest number of specifically enriched proteins (517), while kinetochore proteins formed the largest and smallest fraction of the kinetochore and *CEN\** proteomes, respectively (Figure 1B). Clustering analysis revealed greater similarity between *CEN* and *CEN\** chromatin from mitotically cycling cells than to either of the meiotic samples and showed that the presence of a centromere affects meiotic chromatin composition more than meiotic stage (Figure 1C; Supplemental Table S1). We identified groups of proteins enriched on meiotic chromatin (cluster 1), meiotic centromeres (cluster 2) or on metaphase I centromeres (cluster 3). These clusters corresponded to the expected functional groups: for example, cluster 2, characterised by meiotic and centromere-dependent enrichment (Figure 1C), included proteins involved in centromere coupling, initiation of synapsis, kinetochore monoorientation and the Aurora B kinase-containing chromosome passenger complex (CPC). The kinetochore proteome also showed stage-dependent clustering similar to *CEN* chromatin (Figure S1A; Supplemental Table S2), with clusters of proteins enriched at kinetochores during meiosis (KTcluster 1), and specifically at either meiotic metaphase I (KTcluster 2), or meiotic prophase I (KTcluster 4). Use of a microtubule depolymerizing drug to arrest cells in mitotic metaphase resulted in a kinetochore proteome that was similar to that of cycling cells except for an expected large increase in spindle checkpoint proteins (Mad1^MAD1^, Mad2^MAD2^, Bub1^BUB1^, Bub3^BUB3^), a modest increase in cohesin and Cdc5^Plk1^, and decreased levels of the Mcm2-7 replicative helicase (Figure S1B). Therefore, our purification and analysis workflow can detect cell cycle-dependent changes in chromatin, centromere, and kinetochore composition.

### Changes in chromatin, centromere and kinetochore composition during meiosis

We first mined our dataset to understand how the composition of chromatin changes in meiosis. Comparison of prophase I and metaphase I *CEN\** chromatin to that of cycling cells revealed specific enrichment of the meiosis-specific cohesin subunit, Rec8, and depletion of mitosis-specific Scc1 (Figures S2A and B). Meiotic axis (Hop1, Red1), and synaptonemal complex-nucleating ZMM (**Z**ip1^SYCP1^-Zip2^SHOC1^- Zip3^RNF212^-Zip4^TEX11^, **M**sh4^MSH4^-Msh5^MSH5^, **M**er3^HFM1^) proteins were enriched in prophase I, together with the STR resolvase (**S**gs1^BLM^, **T**op3^TOPIII^*^α^*, **R**mi1^RMI1/RMI2^), which is consistent with their roles in meiotic recombination and synapsis.

Next, we focused on understanding how the protein composition of centromeres and kinetochores changes during meiosis (Figure S3). Clustering of only those proteins that specifically associate with functional centromeres (*CEN* and not *CEN\**), at each stage, identified 3 clusters of particular interest (Figure 2A). *CEN*cluster 6 proteins show increased centromere association during both meiotic prophase and metaphase I and include the ZMM proteins, which, with the exception of Mer3^HFM1^, were strongly associated with centromeres and kinetochores during prophase I (Figures 2B and C; Figure S2). *CEN*cluster 5 proteins are specifically enriched on meiotic metaphase I centromeres and include Cdc5^Plk1^ kinase, which is recruited to kinetochores at prophase I exit to establish monoorientation (Galander et al., 2019) (Figures 2B and C). *CEN*cluster 3 proteins are associated with centromeres of cycling and meiotic metaphase I cells, but depleted at prophase I. Consistently, this cluster included outer kinetochore complexes Ndc80c^NDC80c^ and Dam1c, which are shed at prophase I due to specific degradation of Ndc80^NDC80^ protein (Chen et al., 2020; Meyer et al., 2015a; Miller et al., 2012) (Supplemental Table S3). Direct comparison of the prophase I and metaphase I datasets revealed extensive remodelling of kinetochores upon prophase exit (Figures 2B and C). **Z**ip1^SYCP1^ together with SZZ (**S**po16^SPO16^, **Z**ip2^SHOC1^, **Z**ip4^TEX11^) and Msh4^MSH4^- Msh5^MSH5^ complexes are lost from *CEN* chromatin and kinetochores as cells transition from prophase I to metaphase I. They are replaced by outer kinetochore proteins (Ndc80c^NDC80c^ and Dam1c) together with spindle pole body components and microtubule-associated proteins. Spc105^KNL1^, along with its binding partner Kre28^Zwint^, was also specifically depleted from prophase I *CEN* chromatin and kinetochore purifications, returning in metaphase I (Figures 2B and C). In contrast, both Mtw1c^MIS12c^ and Ctf19c^CCAN^ were found associated with *CEN* chromatin at all stages analysed (Figure 2B). We also observed differences between *CEN* and kinetochore purifications. Chromatin assembly factor I (CAF-I, Cac2-Mri1-Rlf2) was enriched on metaphase I *CEN* chromatin (Figure 2B); Ubr2-Mub1, which is known to regulate Dsn1^DSN1^ stability in mitotic cells (Akiyoshi et al., 2013b), was found only on prophase I kinetochore preparations (Figure 2C).

**Figure 2.**
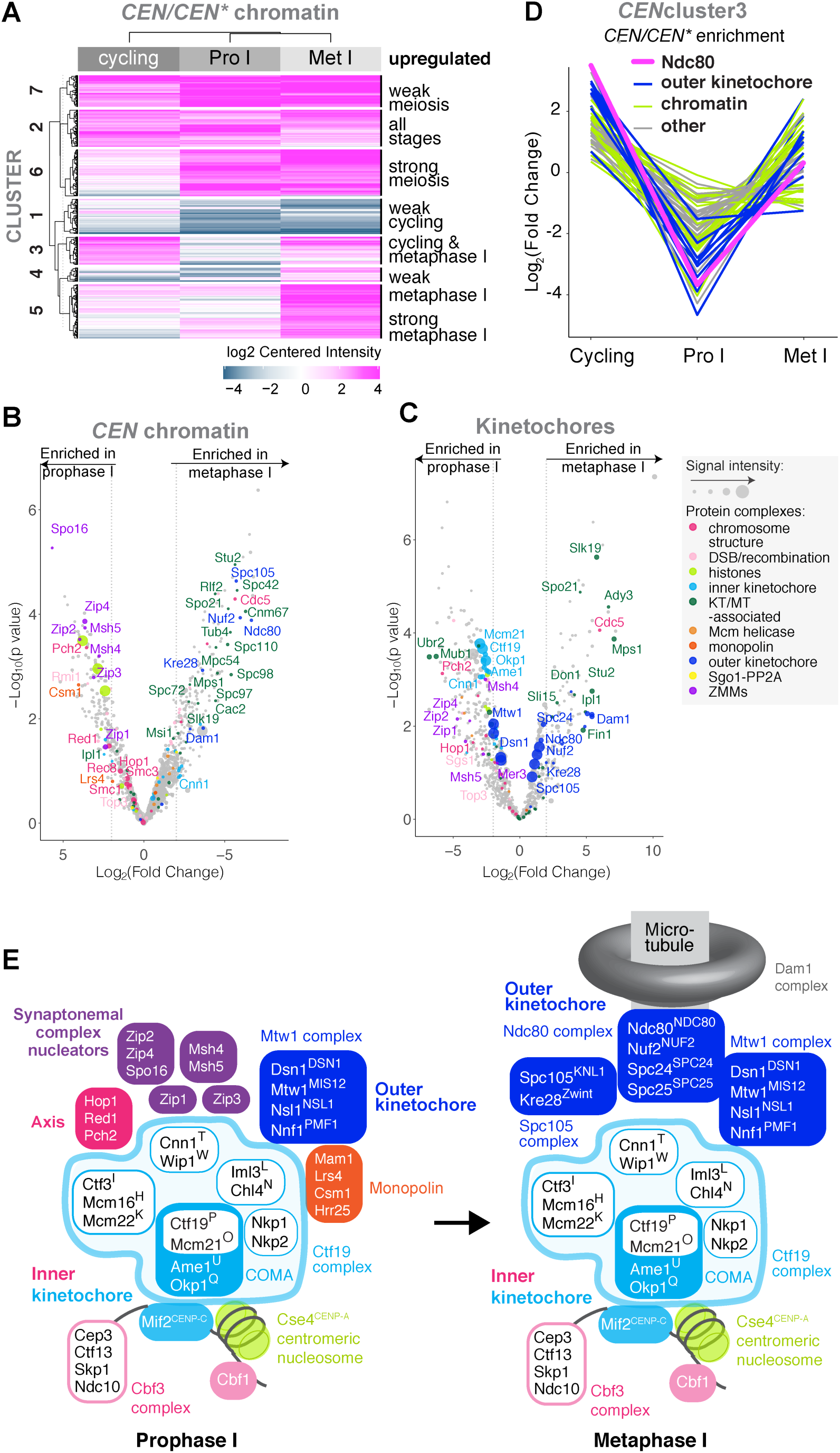
Remodelling of the centromeric and kinetochore proteome throughout the meiotic cell divisions. (A) *CEN* chromatin exhibits distinct composition signatures at different stages. Clustering of *CEN* chromatin/*CEN\** chromatin enrichment value for each of cycling, prophase-arrested and metaphase-arrested conditions were clustered (k-means) to identify groups of proteins with similar behaviour. A cut-off of Log_2_(Fold Change) > 2 and p < 0.05 was used. (B and C) Composition of *CEN* chromatin (B) or kinetochore particles (C) isolated from prophase I and metaphase I is strikingly different. Volcano plot presenting the LFQMS-identified proteins co-purifying with *CEN* plasmids (B) or Dsn1-6His-3FLAG (C) immunopurified from cells arrested in prophase I (*ndt80*Δ (B), inducible*-ndt80* (C)) and metaphase I (*pCLB2-CDC20*). Log_2_(Fold Change) between conditions are shown with their corresponding p values (see methods). Dashed line indicates Log_2_(Fold Change) = |2|. (D) Several proteins exhibit Ndc80-like depletion from centromeres specifically during meiotic prophase. Mean-centered Log_2_(Fold Change) from *CEN*cluster3 is plotted to show abundance of individual proteins in the indicated stages. (E) The extent of kinetochore remodelling during meiosis observed by proteomics is depicted as a schematic. See also Supplemental Figures S1, S2 and S3 and Supplemental Table S3.

To visualise proteins that show a similar timing of kinetochore association to Spc105c^KNL1c^, Ndc80c^NDC80c^ and Dam1c, we plotted the intensity of proteins in *CEN*cluster3 at the different stages (Figure 2D; Supplemental Table S3). In addition to the Ndc80c^NDC80c^-associated PP1 phosphatase regulator, Fin1, and the microtubule regulator Stu2^XMAP215^ (Akiyoshi et al., 2009a; Miller et al., 2016), several chromatin regulators were depleted from centromeric chromatin during prophase I, suggesting a need for centromere re-structuring in preparation for chromosome segregation. Therefore, our proteomics has revealed how kinetochores are remodelled between meiotic prophase and metaphase I, shown schematically in Figure 2E.

### The Ctf19 complex is essential for meiotic chromosome segregation

The remodelling of centromeric chromatin and kinetochores during meiosis suggests the existence of specialised assembly mechanisms. Because the Ctf19c^CCAN^ persists on centromeres through all stages of vegetative growth and gametogenesis, we hypothesised that it plays a critical role in this re-structuring. Ctf19c^CCAN^ proteins that are dispensable for vegetative growth (Measday et al., 2002; Meluh and Koshland, 1997; Ortiz et al., 1999; Pot et al., 2003; De Wulf et al., 2003) become essential for chromosome segregation during meiosis (Fernius and Marston, 2009; Marston et al., 2004) and Ctf19^CENP-P^ is implicated in meiotic kinetochore assembly (Mehta et al., 2014). Consistently, cells lacking the non-essential Ctf19^CENP-P^, Mcm21^CENP-O^, Chl4^CENP-N^ or Iml3^CENP-L^ show ∼50% wild-type viability after mitosis (Figure 3A), while only around ∼1-10% of tetrads produced four viable spores after meiosis (Figure 3B). Furthermore, although histone H2B (Htb1-mCherry)-labelled nuclei partitioned uniformly in live mitotic *mcm21Δ* cells, meiosis resulted in profoundly unequal distribution into gametes (Figure 3C-E). Therefore, Ctf19c^CCAN^ proteins that are dispensable for mitotic growth, are essential for meiosis.

**Figure 3.**
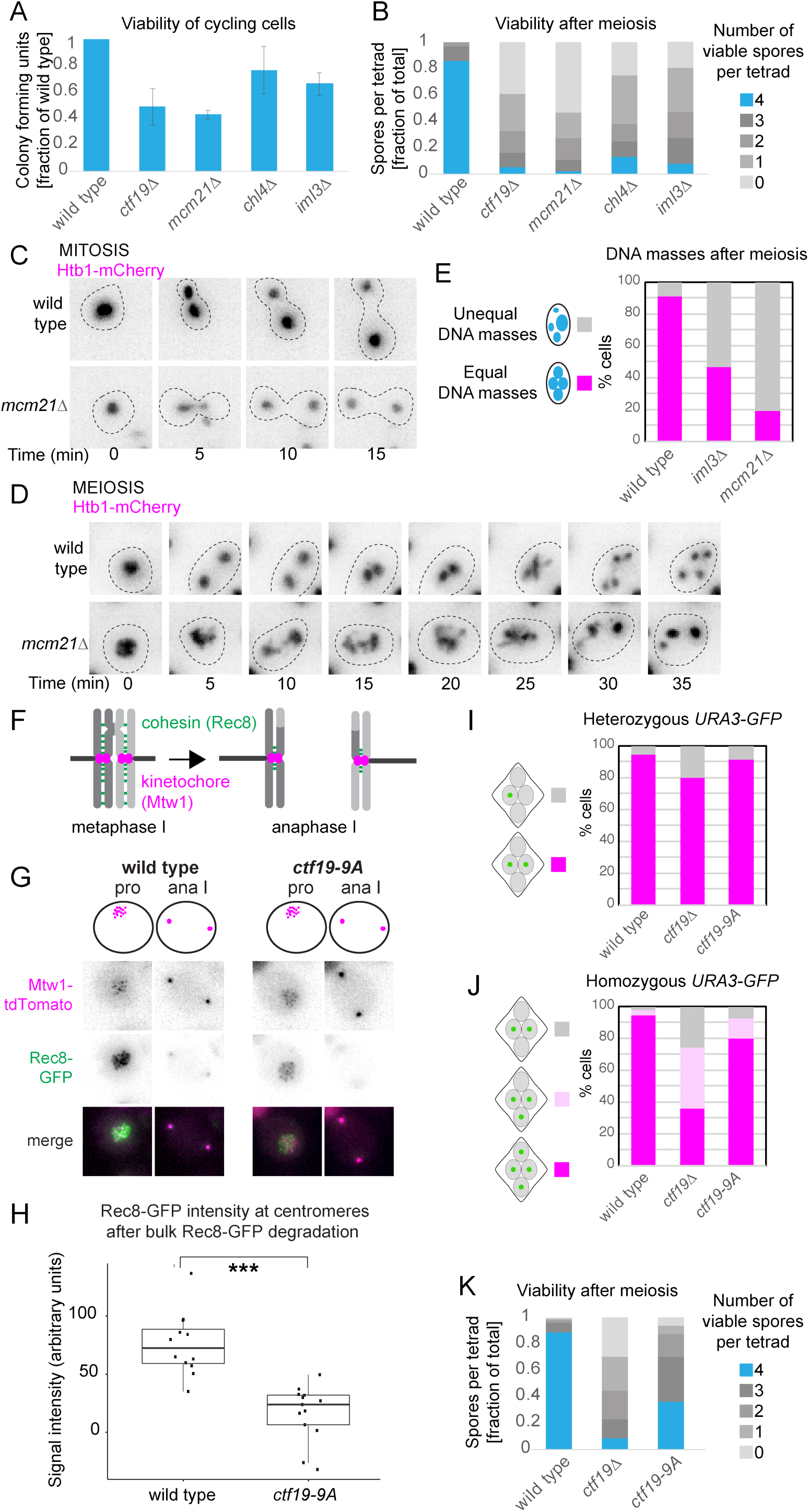
Essential roles of Ctf19c^CCAN^ in meiosis include, but are not limited to, centromeric cohesion establishment. (A) Deletion of non-essential Ctf19c^CCAN^ components mildly impairs mitotic viability. Viability of cycling cells of the indicated genotypes is shown after plating as proportion of wild-type. n = 3 – 4 biological replicates, minimum of 200 cells plated for each genotype. (B) Ctf19c^CCAN^ components that are non-essential for mitotic growth are critical for viability after meiosis. The number of viable progeny was scored following dissection of 36 tetrads for each genotype. (C) Deletion of *MCM21* does not cause gross chromosome missegregation in mitotic cells. Mitotically cycling wild-type and *mcm21*Δ cells expressing Htb1-mCherry were imaged. t = 0 min time-point is defined as the last time-point before DNA masses attempt to separate. (D and E) Deletion of *MCM21* causes gross chromosome missegregation in meiotic cells. (D) Wild-type and *mcm21*Δ cells expressing Htb1-mCherry were released from prophase I arrest and imaged through meiosis. t = 0 min time-point is defined as the last time-point before DNA masses attempt to separate. (E) Quantification of D. 58-92 cells were scored. (F-H) Pericentromeric cohesin is absent in *ctf19-9A* anaphase I cells. Wild-type and *ctf19-9A* cells expressing Rec8-GFP and Mtw1-tdTomato were imaged throughout meiosis. (F) Schematic showing Rec8^REC8^ loss from chromosome arms, but not pericentromeres in anaphase I (G) Representative images are shown. (H) Quantification of Rec8-GFP signal in the vicinity of Mtw1-tdTomato foci immediately following bulk Rec8-GFP degradation. *** p < 10^-5^; Mann-Whitney test. n > 11 cells. (I and J) *ctf19-9A* cells show meiosis II chromosome segregation defects that are less severe than *ctf19Δ* cells. The percentage of tetra-nucleate cells with the indicated patterns of GFP dot segregation was determined in wild-type and *ctf19-9A* cells with either one copy (heterozygous, I) or both copies (homozygous, J) of chromosome V marked with GFP at *URA3* locus. n = 2 biological replicates, 100 tetrads each; mean values are shown. (K) Spore viability of *ctf19-9A* cells is impaired, but less severe than *ctf19Δ* cells. The number of viable progeny was scored following tetrad dissection. n = 3 biological replicates, > 70 tetrads each; mean values are shown. Error bars represent standard error.

### Centromeric cohesin enrichment is not the only critical meiotic function of Ctf19c^CCAN^

The Ctf19c^CCAN^ directs cohesin loading at centromeres, establishing robust pericentromeric cohesion, which ensures accurate segregation of sister chromatids during meiosis II (Eckert et al., 2007; Fernius et al., 2013; Marston et al., 2004; Ng et al., 2009). Interestingly, Ctf19c^CCAN^ mutants show chromosome segregation defects also during meiosis I (Fernius and Marston, 2009; Mehta et al., 2014), suggesting meiotic roles important for homolog segregation. We analysed the *ctf19-9A* mutation, which fails to localise cohesin to centromeres in mitosis, while leaving other functions of the kinetochore intact (Hinshaw et al., 2017). In contrast to wild-type cells, in which Rec8-containing cohesin was specifically retained at pericentromeres following anaphase I, pericentromeric cohesin was absent in *ctf19-9A* cells (Figure 3F-H). Consistently, while *ctf19-9A* impaired segregation of GFP-labelled sister chromatids during meiosis II (Figure 3I and J), it did not impact chromosome segregation and spore viability as severely as *ctf19Δ* (Figure 3I-K). Therefore, while Ctf19-directed cohesin loading is crucial for accurate for meiosis II chromosome segregation, additional Ctf19c^CCAN^ function(s) that are essential for meiosis must exist.

### Ctf19c^CCAN^ is central for kinetochore integrity and remodelling during meiosis

To understand the role of Ctf19c^CCAN^ in kinetochore remodelling during meiosis, we exploited our *CEN* chromatin proteomics pipeline. We selected the Ctf19c^CCAN^ proteins Mcm21^CENP-O^ and Iml3^CENP-L^ for further analysis (Figure 2E). Cycling, prophase I, and metaphase I *mcm21Δ* and *iml3Δ CEN* chromatin datasets revealed significant deviations in composition from wild type, affecting multiple different complexes of both the core kinetochore and those with kinetochore-associated functions (Figure 4A, Figure S4A and B). In cycling *mcm21Δ* and *iml3Δ* cells, in addition to expected reductions in Ctf19c^CCAN^ components, the abundance of the central Mtw1c^MIS12c^ and outer kinetochore complexes was modestly decreased. In cycling *mcm21Δ*, but not *iml3Δ* cells, levels of Cse4^CENP-A^ and Mif2^CENP-C^ were reduced (Figure 4A, Figure S4A and B, Cycling cells). Recovery of cohesin was also less efficient in *mcm21Δ* and *iml3Δ* cycling cells, consistent with the key role of the Ctf19c^CCAN^ in kinetochore-targeted cohesin loading (Hinshaw et al., 2017) (Figure 3F-H). At prophase I, the outer kinetochore components Ndc80c^NDC80c^, Dam1c, and Spc105c^KNL1c^ were largely depleted from wild-type, *mcm21Δ,* and *iml3Δ CEN* chromatin, as expected (Figure 4A, Prophase I). Interestingly, we found that Mcm21^CENP-O^ and Iml3^CENP-L^ are required for association of Msh4^MSH4^-Msh5^MSH5^, SZZ and STR complexes with *CEN* chromatin in prophase I. Furthermore, Mtw1c^MIS12c^ was lost from *mcm21Δ* and *iml3Δ* prophase I and metaphase I *CEN* chromatin, and Ndc80c^NDC80c^, Dam1c, and Spc105c^KNL1c^ did not reappear in metaphase I (Figure 4A, Figure S4A and B, Metaphase I). Consistently, microtubule-associated and spindle pole body proteins were recovered with wild-type, but not *iml3Δ* or *mcm21Δ,* metaphase I *CEN* chromatin (Figure 4A, Figure S4A and B, Metaphase I). Surprisingly, we also observed effects on the inner kinetochore. Although the DNA-binding Cbf3c was recruited at normal levels in all stages in *iml3Δ* or *mcm21Δ* cells, Mcm21^CENP-O^ and Iml3^CENP-L^ were required for proper centromeric association of Cse4^CENP-A^and Mif2^CENP-C^ at metaphase I (Figure 4A). We conclude that the Ctf19c^CCAN^ plays a major role in ensuring the integrity of the kinetochore in meiotic prophase, its reassembly upon prophase I exit, and in orchestrating kinetochore remodelling in meiosis.

**Figure 4.**
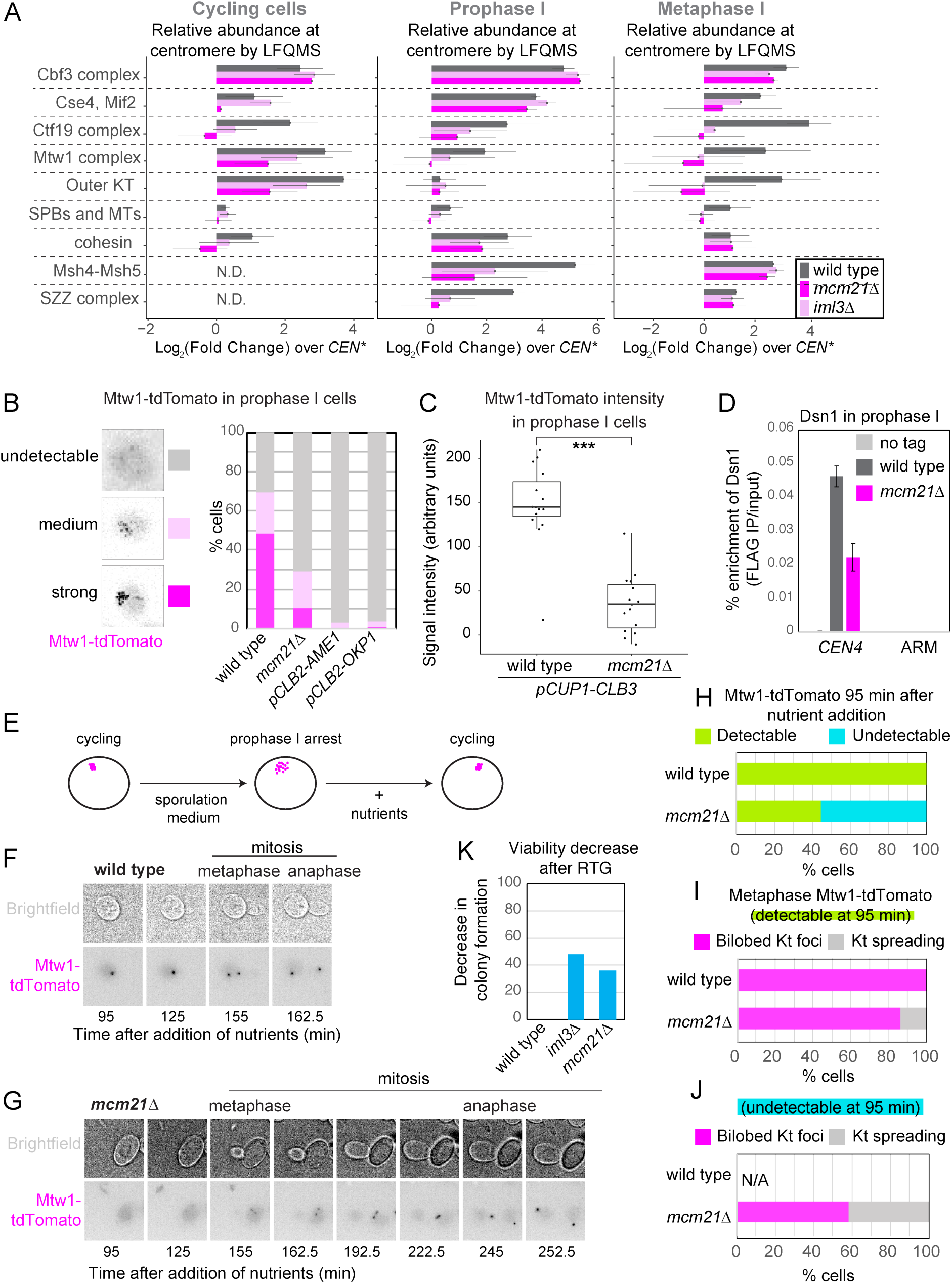
Proteomics identifies a critical role of Ctf19c^CCAN^ in meiotic kinetochore assembly. (A) Global *CEN/CEN\** proteomics reveals that kinetochore composition is radically altered in *mcm21Δ* and *iml3Δ* meiotic prophase and metaphase I cells. Sum of LFQMS abundance of protein complexes on *CEN* chromatin in wild-type, *iml3*Δ and *mcm21*Δ cells are shown as enrichment over *CEN\** chromatin isolated from wild-type cells. The abundance of Iml3 ^CENP-L^ and Mcm21^CENP-O^ proteins was not included in the total Ctf19c^CCAN^ count, as these proteins are missing in *iml3*Δ and *mcm21*Δ cells, respectively (see methods). KT – kinetochore, MT – microtubule, SPB – spindle pole body, SZZ – Spo16^SPO16^, Zip2^SHOC1^, Zip4^TEX11^. (B-D) Mtw1c^MIS12c^ components Mtw1^MIS12^ (B and C) and Dsn1^DSN1^ (D) are lost from kinetochores in the absence of essential and non-essential Ctf19c^CCAN^ components. (B) Wild-type, *mcm21Δ, pCLB2-AME1* and *pCLB2-OKP1* cells were imaged immediately after release from prophase I arrest and representative images and scoring of cells with Mtw1-tdTomato signal are shown. n > 58 cells. (C) Quantification of Mtw1-tdTomato signal intensity in prophase I arrested wild-type and *mcm21*Δ cells ectopically expressing Clb3 to ensure kinetochore clustering and allow signal quantification. ***, p < 10^-5^; Mann-Whitney test. 15 cells were scored. (D) Prophase I-arrested wild-type and *mcm21*Δ cells carrying *ndt80Δ* and *DSN1-6His-3FLAG,* together with untagged control were subjected to anti-FLAG ChIP-qPCR. Error bars represent standard error (n=4 biological replicates). p < 0.05, paired t-test. (E-J) Mtw1c^MIS12c^ binding and kinetochore function is not restored in *iml3Δ* and *mcm21Δ* cells in the mitotic division following return to growth. (E) Schematic of return-to-growth experiments. (F and G) Mtw1-tdTomato fails to localise to kinetochores in the absence of *MCM21* following return to growth. Nutrients were added to wild-type (F) and *mcm21*Δ (G) cells arrested in prophase I. Imaging commenced 95 minutes after nutrients addition. (H and I) Loss of kinetochore-localised Mtw1-tdTomato correlates with further kinetochore spreading. (H) Fraction of cells with detectable (green) and undetectable (blue) Mtw1-tdTomato signal 95 min after return to growth is shown. n > 64 cells. (I-J) Following the cells scored in (H), the appearance of Mtw1-tdTomato signal at metaphase in those cells in which foci were detectable (n > 28 cells, I) or undetectable (n > 29 cells, J) upon return to growth is shown. Numbers of cells in H vs. I and J are not identical, as it was not possible to score all the cells later in mitosis. (K) Cells lacking *IML3* or *MCM21* exhibit loss of viability following return to growth. Wild-type, *iml3Δ* and *mcm21Δ* cells were induced to sporulate and plated at t_0h_ (before meiosis) and t_5h_ (prophase I arrest). Viability drop from t_0h_ to t_5h_ was calculated. n > 158 cells. RTG – return to growth. See also Supplemental Figure S4 and S5.

### Non-essential Ctf19c^CCAN^ subunits are critical for retention of Mtw1c^MIS12c^ at prophase I kinetochores

Our *CEN* proteomics shows that Mtw1c^MIS12c^ is lost from prophase I kinetochores in *mcm21Δ* and *iml3Δ* cells. Because Mtw1c^MIS12c^ structurally links the inner and outer kinetochore and is essential for chromosome segregation and cell viability, the lack of Mtw1c^MIS12c^ could explain the severe effects observed in *mcm21Δ* and *iml3Δ* cells. To visualise Mtw1c^MIS12c^ behaviour in meiosis, we performed live-cell imaging of its Mtw1^MIS12^ subunit tagged with tdTomato in wild-type and Ctf19^CCAN^-deficient cells. At meiotic prophase I, kinetochores are de-clustered and appear as up to 16 individual foci, each representing a pair of homologous centromeres (Figure 4B). De-clustered Mtw1-tdTomato foci were detected in ∼70% of wild-type prophase I cells, but only ∼30% of *mcm21Δ* cells (Figure 4B). Meiotic depletion of Ame1^CENP-U^ or Okp1^CENP-Q^ (by placement under control of the mitosis-specific *CLB2* promoter (Lee and Amon, 2003)) also largely prevented detection of Mtw1-tdTomato in prophase I cells (Figure 4B). In mitotic cells, a version of Ame1^CENP-U^ lacking the Mtw1c^MIS12c^ receptor motif leads to a reduction in Mtw1c^MIS12c^ incorporation into the kinetochore and is unable to support growth (Hornung et al., 2014). Taken together, these data suggest that the Ctf19c^CCAN^ is important for stable Mtw1c^MIS12c^ association with centromeres in mitosis, but is crucial for meiosis, where subunits that are non-essential for mitosis, Mcm21^CENP-O^ and Iml3^CENP-L^, are additionally required (Figure 4B).

To ease Mtw1-tdTomato signal quantification at kinetochores, we overexpressed the cyclin, *CLB3*, which precludes kinetochore detachment from spindle pole bodies, thereby preventing de-clustering (Miller et al., 2012). In *CLB3*- overexpressing prophase I cells, Mtw1-tdTomato intensity was decreased in *mcm21Δ* compared to wild-type cells (Figure 4C). Consistently, chromatin immunoprecipitation followed by qPCR (ChIP-qPCR) showed that Dsn1^DSN1^ was significantly reduced at prophase I centromeres of *mcm21Δ* cells (Figure 4D), confirming loss of Mtw1c^MIS12c^ from kinetochores.

To determine whether the loss of Mtw1c^MIS12c^ from kinetochores at prophase I has an irreversible effect on the ensuing chromosome segregation, we performed a return-to-growth experiment. Prophase I-arrested cells that have undergone DNA replication and meiotic recombination are not committed to meiosis; consequently, addition of nutrients causes them to re-enter the mitotic programme (Tsuchiya et al., 2014) (Figure 4E). As expected, upon return-to-growth, wild-type cells re-clustered kinetochores, budded and the single Mtw1-tdTomato focus split into two foci that segregated into the daughter cells (Figure 4F). In contrast, in the majority of *mcm21Δ* cells returning to growth, Mtw1-tdTomato foci were initially undetectable; faint foci appeared in these cells around the time of bud emergence, splitting into daughter cells with a substantial delay (Figure 4G). Examination of the fate of those *mcm21Δ* cells in which Mtw1-tdTomato foci were undetectable 95 min after adding nutrients revealed an increased tendency for Mtw1-tdTomato foci to spread in metaphase, suggestive of failed microtubule attachments (Figure 4H-J). Consistently, *mcm21Δ* cells show decreased viability after return to growth (Figure 4K). Therefore, at the mitosis to meiosis transition, kinetochores undergo a precipitous and irreversible event that relies upon the presence of the Ctf19c^CCAN^ to preserve their capacity to attach to microtubules.

### Preventing outer kinetochore shedding in prophase I does not restore kinetochore function in Ctf19c^CCAN^-deficient metaphase I cells

Re-synthesis of the Ndc80^NDC80^ protein at prophase I exit allows outer kinetochore assembly and re-attachment to microtubules (Chen et al., 2017, 2020; Miller et al., 2012). In mitotic cells, an outer kinetochore Dam1c component, Ask1, can recruit inner kinetochore components, including the centromeric histone, Cse4^CENP-A^, when tethered to a chromosomal arm region (Ho et al., 2014; Kiermaier et al., 2009; Lacefield et al., 2009). We therefore hypothesised that production of the outer kinetochore at prophase I exit might be sufficient to stabilise kinetochores in *mcm21Δ* cells. However, in contrast to wild-type cells (Figure 5A), Ndc80-GFP re-accumulation at kinetochores was delayed in *mcm21Δ* cells, and the observed foci were fainter (Figure 5A and B). Where Mtw1-tdTomato was detected in *mcm21Δ* cells, individual kinetochore foci tended to spread, rather than form bilobed foci typical of metaphase I (Figure 5C, see below). This suggested that outer kinetochore assembly was aberrant in *mcm21Δ* cells at prophase I exit. Consistently, prevention of outer kinetochore shedding by *CLB3* overexpression was unable to restore Mtw1-tdTomato localization in *mcm21Δ* cells (Figure 4C). Furthermore, expression of a non-degradable version of Ndc80^NDC80^, which prevents outer kinetochore shedding in prophase I (Ndc80(Δ2-28); (Chen et al., 2020)), did not restore Dsn1-tdTomato kinetochore localisation in *mcm21Δ* prophase I cells (Figure 5D). Therefore, kinetochore shedding during prophase I is not the cause of kinetochore disintegration in *mcm21Δ* and *iml3Δ* meiotic cells.

**Figure 5.**
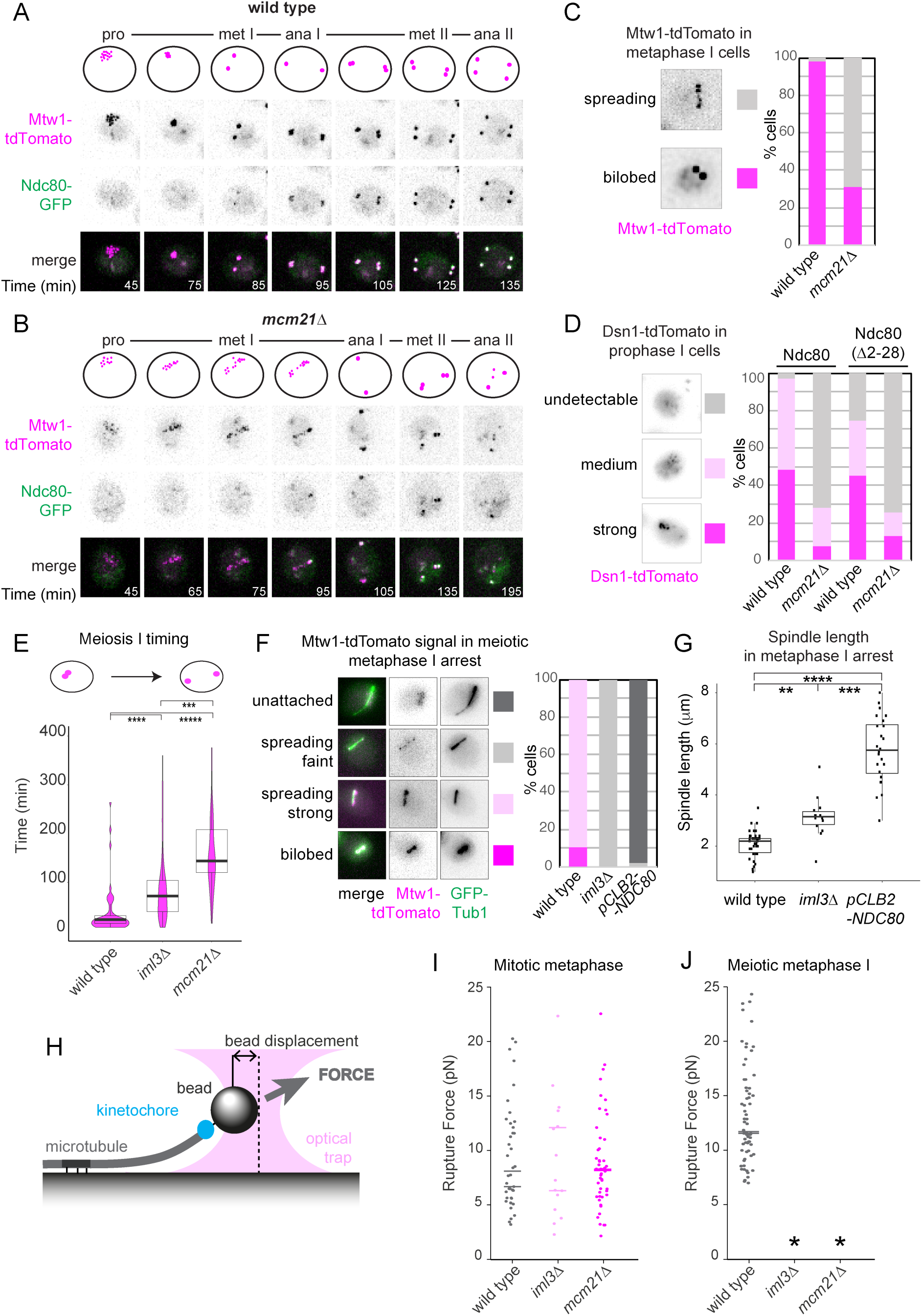
An intact Ctf19c^CCAN^ is required for outer kinetochore assembly and kinetochore microtubule attachments in meiosis I. (A-C) Abnormal kinetochore behaviour in the absence of *MCM21.* Representative images of live wild-type (A) and *mcm21*Δ (B) cells carrying Mtw1-tdTomato and Ndc80-GFP released from prophase I arrest and imaged throughout meiosis. Time after release from prophase I is indicated. (C) Scoring of Mtw1-tdTomato signal in A and B. Cells showing kinetochore spreading in at least one timepoint during the time-lapse were included in the “spreading” category. n > 49 cells (D) Non-degradable *ndc80(Δ2-28)* does not rescue kinetochore function upon the loss of *MCM21.* Dsn1-tdTomato signal was scored in prophase I wild-type and *mcm21*Δ cells expressing either Ndc80-GFP or Ndc80(Δ2-28)-GFP. n > 56 cells. (E) Cells lacking non-essential components of the Ctf19c^CCAN^ are delayed in meiosis I chromosome segregation. Wild-type, *iml3*Δ and *mcm21*Δ cells expressing Mtw1-tdTomato and GFP-Tub1 were released from prophase arrest and imaged. Time between the emergence of a bilobed kinetochore structure and the last timepoint before emergence of two pairs of kinetochore foci was measured. *** p < 10^-4^, **** p < 10^-8^, ***** p< 10^-15^; Mann-Whitney test. n > 37 cells. (F and G). Meiosis I spindles are elongated in *iml3Δ* cells. Wild-type and *iml3*Δ cells carrying *pCLB2-CDC20* and expressing Mtw1-tdTomato and GFP-Tub1 were imaged undergoing meiosis. (F) Representative images (left) and scoring of Mtw1-tdTomato appearance in cells with spindles. Cells showing unattached or ‘spreading’ kinetochores in at least one timepoint during the time-lapse were included in these categories. n > 52 cells. (G) Spindle lengths measured 75 min after spindle emergence (see methods). **** p < 10^-12^,*** p < 10^-8^, ** p < 10^-3^; t-test. n = 15 cells (*iml3Δ*) or 65 cells (wild type). (H-J) Purified kinetochore particles (Dsn1-6His-3FLAG immunoprecipitation) from *iml3Δ* and *mcm21Δ* cells fail to attach to microtubules in a single molecule assay. (H) Schematic of assay showing the optical trap pulling on a bead attached to a coverslip-immobilised microtubule. The bead-microtubule interaction is facilitated by purified kinetochores. (I and J) Kinetochore particles isolated from meiosis I-arrested cells lacking *IML3* and *MCM21* are not able to form kinetochore-microtubule attachments *in vitro.* Rupture force measurements of kinetochore particles isolated from mitotically arrested (I) or meiosis metaphase I-arrested (J) wild-type, *iml3*Δ and *mcm21*Δ cells are shown. Total particles analysed: n = 65 (wild type, meiosis), n = 41 (wild type, mitosis),), n = 15 (*iml3Δ*, mitosis), n = 49 (*mcm21Δ*, mitosis from 2 biological replicates; bar represents median for each replicate. (I) Cycling cells were arrested in mitotic metaphase by the addition of benomyl. (J) Cells were arrested in meiotic metaphase I due to the presence of *pCLB2-CDC20* allele. Asterisks indicate conditions for which no initial kinetochore-microtubule attachment were formed and thus rupture force could not be measured. See also Supplemental Figure S5.

### Ctf19c^CCAN^ is critical for kinetochore-microtubule attachments in meiosis I

We noticed two characteristic features of metaphase *mcm21Δ* cells. First, while bilobed Mtw1-tdTomato foci were observed in wild-type metaphase I cells, in *mcm21Δ* cells multiple faint foci appeared with similar timing and spread along a linear axis (Figure 5B and C). Second, the duration of meiosis I was extended in *mcm21Δ* and, to a lesser extent, in *iml3Δ* cells (Figure 5E). Together, these observations suggest that kinetochores are not attached to microtubules and/or improperly assembled, leading to activation of the spindle checkpoint in Ctf19c^CCAN^ mutants. Consistent with this idea, we observed a delay in the degradation of anaphase inhibitor securin (Pds1^SECURIN^) and cleavage of cohesin in both *mcm21Δ* and *iml3Δ* cells (Figure S5A-C). To further examine the importance of the Ctf19c^CCAN^ for kinetochore-microtubule attachments, we arrested wild-type and *iml3Δ* cells carrying GFP-labelled alpha tubulin to label microtubules, and Mtw1-tdTomato in metaphase I. As a control, we also imaged Ndc80^NDC80^-depleted cells (by placement under control of the mitosis-specific *CLB2* promoter (Vincenten et al., 2015)), where kinetochore-microtubule attachments are absent. Compared to wild-type cells, in which the Mtw1-tdTomato signal was localized along short spindles, in both *pCLB2-NDC80* and *iml3Δ* cells the Mtw1-tdTomato signal was reduced (Figure 5F). While Mtw1-tdTomato foci were displaced from highly elongated spindles in *pCLB2-NDC80* cells, *iml3Δ* cells showed spreading of faint Mtw1-tdTomato along modestly extended spindles (Figure 5F and G, see also Figure S5B). This indicates that kinetochore-microtubule attachments are impaired, but not absent, in meiotic cells lacking Ctf19c^CCAN^. To address this directly, we purified kinetochores from both mitotic and meiotic metaphase I wild-type, *mcm21Δ* and *iml3Δ* cells (Dsn1-6His- 3FLAG immunoprecipitation; Figure 1A) and assayed their ability to resist laser trap forces after binding to microtubules growing from a coverslip-anchored seed (Figure 5H; (Akiyoshi et al., 2010; Sarangapani et al., 2014)). Wild-type kinetochore particles from mitotic metaphase cells bound microtubules with a mean rupture force of ∼9pN, and this was unchanged for kinetochore particles purified from *mcm21Δ* or *iml3Δ* mitotically cycling cells (Figure 5I). Wild-type meiotic metaphase I kinetochore particles showed an increased rupture force (mean ∼12pN), as reported previously (Sarangapani et al., 2014), whereas both *mcm21Δ* and *iml3Δ* kinetochore particles failed to bind to microtubules (Figure 5J). Therefore, meiotic kinetochores in *mcm21Δ* and *iml3Δ* cells fail to reassemble after prophase I, resulting in their inability to make load-resisting kinetochore-microtubule attachments during metaphase I.

### The essential inner kinetochore is lost at meiotic entry in cells lacking Ctf19c^CCAN^ components

The common effects of *MCM21* deletion and Ame1^CENP-U^ or Okp1^CENP-Q^ depletion on the association of Mtw1c^MIS12c^ with the kinetochore (Figure 4B) suggested the possibility that Mcm21^CENP-O^ might become important for Ame1^CENP-U^-Okp1^CENP-Q^ association with kinetochores specifically during meiosis. Indeed, ChIP-qPCR revealed reduced levels of Ame1^CENP-U^ at kinetochores of *mcm21Δ* and *iml3Δ* prophase I cells (Figure 6A). Therefore, in meiosis, Ctf19c^CCAN^ proteins that are dispensable for mitotic growth become crucial for localizing the components of the Ctf19c^CCAN^ that are essential for viability.

**Figure 6.**
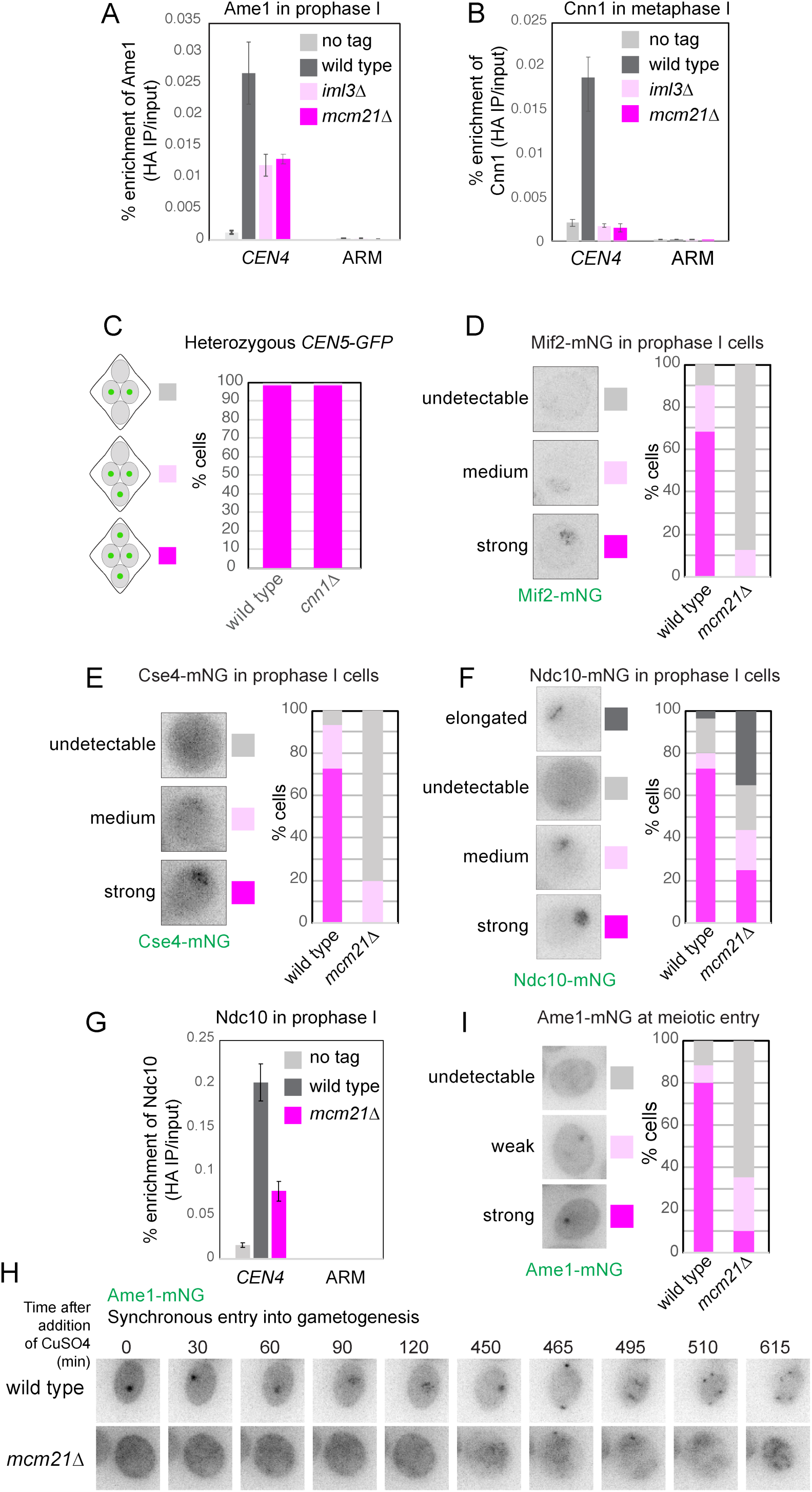
Ctf19c^CCAN^ subunits that are dispensable for mitosis become essential for retention of the inner kinetochore upon entry into gametogenesis. (A) Loss of essential inner kinetochore component Ame1^CENP-U^ in meiotic cells lacking *MCM21*. Prophase I-arrested wild-type, *iml3Δ* and *mcm21*Δ cells carrying *AME1-6HA*, together with untagged control were subjected to anti-HA ChIP-qPCR. Error bars represent standard error (n = 4 biological replicates). p < 0.05, paired t-test. (B) Cnn1^CENP-T^ is lost from kinetochores in the absence of *IML3* and *MCM21*. Metaphase I-arrested wild-type, *iml3Δ* and *mcm21*Δ cells carrying *pCLB2-CDC20* and *CNN1-6HA* together with untagged control were subjected to anti-HA ChIP-qPCR. Error bars represent standard error (n = 3 biological replicates). p < 0.05, paired t-test. (C) *cnn1Δ* cells segregate chromosomes faithfully in meiosis. Wild-type and *mcm21Δ* cells with both copies of chromosome V marked with GFP were sporulated. The percentage of tetra-nucleate cells with the indicated patterns of GFP dot segregation was determined. n = 100 tetrads. (D-G) Loss of non-Ctf19c^CCAN^ inner kinetochore proteins Mif2^CENP-C^ (D), the centromeric nucleosome Cse4^CENP-A^ (E) and Ndc10 (F and G) in meiotic prophase cells lacking *MCM21*. (D-F) Wild-type and *mcm21*Δ cells were imaged immediately after release from prophase I arrest and Mif2-mNeonGreen (D, n > 56 cells), Cse4-mNeonGreen (E, n > 58 cells) or Ndc10-mNeonGreen (F, n > 60 cells) was scored. (G) Prophase I-arrested wild-type and *mcm21*Δ cells carrying *NDC10-6HA* together with untagged control were subjected to anti-HA ChIP-qPCR. Error bars represent standard error (n = 4 biological replicates). p < 0.05, paired t-test. (H and I) Ame1-mNeonGreen is lost from kinetochores immediately upon entry into gametogenesis. Wild-type and *mcm21*Δ cells expressing Ame1-mNeonGreen were allowed to synchronously enter gametogenesis through induction of *IME1* and *IME4*, and followed by live-cell imaging. (H) Representative images are shown. (I) Quantification of Ame1-mNeonGreen appearance at t = 0. n > 226 cells.

As predicted from previous work in mitosis (Hornung et al., 2014), Cnn1^CENP-T^ was also lost from kinetochores in *mcm21Δ* and *iml3Δ* meiotic cells (Figure 6B). However, loss of the Cnn1^CENP-T^-dependent Ctf3^CENP-I^-Ndc80c^NDC80c^ link cannot be responsible for meiotic failure of Ctf19c^CCAN^-compromised cells since *cnn1Δ* cells show no apparent chromosome segregation defects in meiosis (Figure 6C). Remarkably, though below the detection limit of proteomics (Figure 4A), centromeric levels of Mif2^CENP-C^ (Figure 6D), Cse4^CENP-A^ (Figure 6E) and the DNA-binding component, Ndc10, of the Cbf3c (Figure 6F and G), were also decreased in the *mcm21Δ* prophase I cells. Overall, we show that the entire meiotic kinetochore, including its DNA-binding components, is impacted by loss of the non-essential Ctf19c^CCAN^ subunits, while only modest effects on mitotic kinetochores are observed.

To address how early in gametogenesis the inner kinetochore is lost, we imaged wild-type and *mcm21Δ* cells carrying Ame1-mNeonGreen directly after synchronous induction of sporulation (by conditional expression of the meiotic master regulators, Ime1 and Ime4; (Berchowitz et al., 2013; Chia and van Werven, 2016)). Remarkably, Ame1^CENP-U^ was already undetectable at kinetochores upon entry into gametogenesis, and did not appear during the time of imaging (Figure 6H and I). Therefore, components of the Ctf19c^CCAN^ that are non-essential for vegetative growth become essential for kinetochore integrity at the mitosis to meiosis transition.

### Functional targeting modules for the CPC ensure kinetochore integrity in mitosis

Our findings show that, during meiosis, the Ctf19c^CCAN^ is critical for localization of the essential Mtw1c^MIS12c^, which links the inner and outer kinetochore. In mitosis, Ame1^CENP-U^ promotes Mtw1c^MIS12c^ kinetochore localization (Hornung et al., 2014). Our *CEN* chromatin proteome from mitotically cycling *iml3Δ* and *mcm21Δ* cells showed modestly reduced levels of Mtw1c^MIS12c^ and outer kinetochore proteins (Figure 4A, Cycling cells), and these mutants also show a prolonged mitotic metaphase (Figure S6A). These observations prompted us to test whether the non-essential Ctf19c^CCAN^ subunits might also contribute to kinetochore assembly during mitosis.

Mtw1c^MIS12c^ incorporation into kinetochores is enhanced by Ipl1^AURORA B^- dependent phosphorylation of Dsn1^DSN1^, which promotes association of Mtw1c^MIS12c^ with Ame1^CENP-U^-Okp1^CENP-Q^ (Akiyoshi et al., 2009a; Dimitrova et al., 2016). Recently, the Ctf19^CENP-P^ protein has been identified as a kinetochore receptor for Ipl1^AURORA B^ kinase (Fischböck-Halwachs et al., 2019; García-Rodríguez et al., 2019; Knockleby and Vogel, 2009). Centromeric targeting of Ipl1^AURORA B^ occurs by two further mechanisms in mitotic cells, both requiring the CPC subunit Bir1^SURVIVIN^. Simultaneous loss of both Bir1-dependent and Ctf19-dependent recruitment of Ipl1^AURORA B^ to centromeres is lethal (Fischböck-Halwachs et al., 2019; García-Rodríguez et al., 2019). We hypothesised that the inability to recruit Ipl1^AURORA B^ to centromeres results in a failure to incorporate Mtw1c^MIS12c^ into the kinetochore, and lethality. To test this idea, we used the auxin-inducible degron to degrade Bir1-aid upon release of otherwise wild-type and *mcm21Δ* cells from a G1 arrest, and measured Mtw1-tdTomato intensity as cells entered mitosis (Figure 7A-C). Though degradation of Bir1^SURVIVIN^ alone did not decrease Mtw1-tdTomato signal intensity (*BIR1-aid,* NAA; Figure 7A-D), loss of Mcm21^CENP-O^ alone did (*mcm21Δ BIR1-aid,* DMSO; Figure 7A-D), and this was exacerbated upon degradation of BIR1-aid (*mcm21Δ BIR1-aid,* NAA, Figure 7A-D). Furthermore, subsequent cell divisions were largely aborted in the absence of both Ipl1^AURORA B^-targeting modules (*mcm21Δ BIR1-aid,* NAA; Figure 7E), whereas Bir1^SURVIVIN^-depleted cells proliferated similarly to wild-type cells, consistent with the finding that Ipl1^AURORA B^ retains partial activity when Bir1^SURVIVIN^ function is compromised (Makrantoni and Stark, 2009). Therefore, Bir1^SURVIVIN^ and Mcm21^CENP-O^ collaborate not only to direct proper error correction and biorientation (Fischböck-Halwachs et al., 2019; García-Rodríguez et al., 2019), but also, as we show here, in promoting Mtw1c^MIS12c^ binding to the inner kinetochore, presumably through localizing Ipl1^AURORA B^.

**Figure 7.**
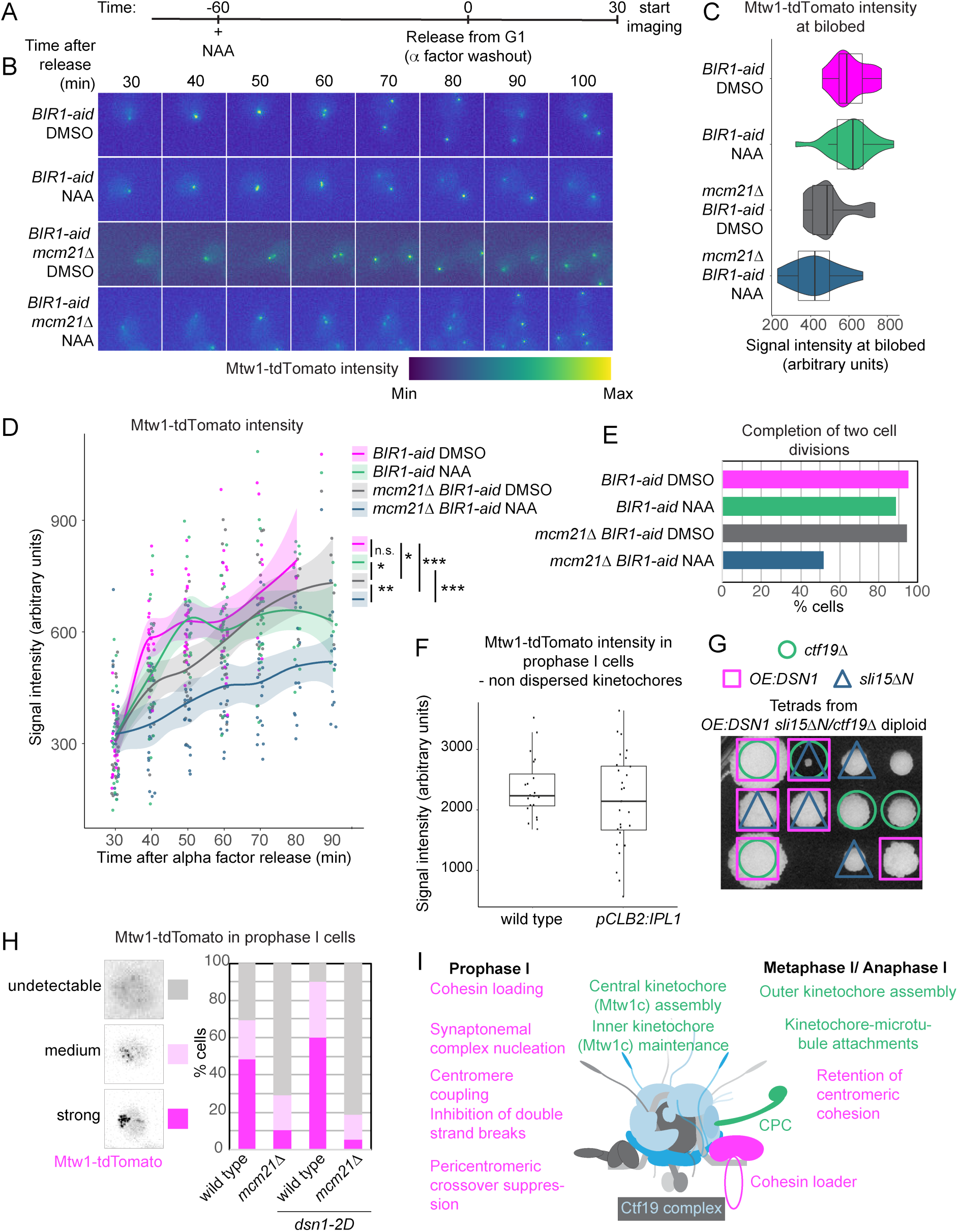
Inner-kinetochore localised Ipl1^AURORA B^ activity is required for kinetochore assembly in mitosis. (A-D) Loss of both of mechanisms of targeting Ipl1 to kinetochores disrupts kinetochore localisation of Mtw1-tdTomato in mitotic cells. (A) Schematic of the experiment shown in B-D. *BIR1-aid* and *mcm21Δ BIR1-aid* cells expressing Mtw1-tdTomato were released from G1 arrest into the mitotic cell cycle, with addition of NAA/DMSO 1 h before release. (B) Representative images of live cells are shown. (C and D) Measurements of Mtw1-tdTomato intensity either (C) at a timepoint where Mtw1-tdTomato signal appeared bilobed or (D) from the beginning of the movie until anaphase (defined as the time point when two separate kinetochore foci reach opposite ends of mother and daughter cells). The last value is an average value of intensities of the two separated Mtw1-tdTomato foci. The line represents a smoothed mean, the shaded area indicates 95% confidence interval. * p < 0.05, ** p < 0.01, *** p < 10^-5^; Mann-Whitney test. n > 19 cells. (E) Mcm21^CENP-O^ and Bir1^SURVIVIN^ play redundant roles in mitotic progression. The proportion of cells that go on to do two or more cell divisions following alpha factor release was determined. n > 33 cells. (F) Meiotic depletion of Ipl1 does not impair kinetochore structure. Intensity of Mtw1-tdTomato foci in wild-type and *pCLB2-IPL1* cells harbouring inducible-*ndt80* was measured in prophase arrest. Intensity was measured only in cells which did not show dispersed kinetochores. n > 19 cells. (G) Overexpression of Dsn1 protein rescues synthetic lethality of *sli15ΔN* and *ctf19Δ*. A diploid which is heterozygous for each of *sli15ΔN*, *ctf19Δ* and multiple extra copies of *DSN1* (*OE:DSN1*) was induced to undergo meiosis and the four resultant spores were separated by dissection and arrayed in rows. The progeny were genotyped for *OE:DSN1* (magenta squares), *sli15ΔN* (blue triangles) and *ctf19Δ* (green circles). Although *sli15ΔN ctf19Δ* cells are inviable, *OE:DSN1 sli15ΔN ctf19Δ* survivors were observed after prolonged growth. (H) Phosphomimetic mutations within Ipl1-dependent Dsn1 phosphorylation sites do not restore kinetochore assembly in cells lacking *MCM21.* Wild-type, *mcm21Δ, dsn1-2D* and *mcm21Δ dsn1-2D* cells were imaged immediately after release from inducible-*ndt80* arrest. Mtw1-tdTomato signal was scored. Data were acquired together with that shown in Figure 4A and wild-type and *mcm21*Δ data are identical. n > 56 cells. *dsn1-2D* – *dsn1-S240D S250D*. (I) Schematic model of cohesin-dependent (magenta) and Ipl1^AURORA B^-dependent (green) roles of the Ctf19 complex in meiotic cell divisions. See also Supplemental Figure S6.

Although several studies have implicated Ipl1^AURORA B^ in stabilizing interactions between the inner and outer layers of the kinetochore (Akiyoshi et al., 2009b; Bonner et al., 2019; Kim and Yu, 2015), paradoxically, its activity does not appear to be essential for outer kinetochore assembly, at least in otherwise wild-type yeast cells (e.g. (Akiyoshi et al., 2013a; Marco et al., 2013; Meyer et al., 2015a; Pinsky et al., 2006). Indeed, we found that Mtw1-tdTomato intensity is not significantly different in wild-type and Ipl1^AURORA B^-depleted cells at clustered kinetochores in meiotic prophase (Ipl1^AURORA B^ was depleted by placement of *IPLl1* gene under control of the mitosis-specific *CLB2* promoter ; Figure 6F). In summary, Ipl1^AURORA B^ targeting to inner kinetochores becomes critical for kinetochore assembly in the absence of functional Ctf19c^CCAN^.

### Increased dosage of *DSN1* rescues the viability of cells lacking CPC targeting modules

If the crucial role of Ipl1^AURORA B^ recruitment to the inner kinetochore by the Bir1^SURVIVIN^- and Ctf19^CENP-P^-dependent pathways is to promote kinetochore integrity, we reasoned that the synthetic lethality upon loss of both of these pathways might be rescued by stabilizing the Ipl1^AURORA B^ target, Dsn1^DSN1^, within the kinetochore. However, phosphomimetic Dsn1-S240D S250D, which carries mutations in Ipl1^AURORA B^ target sites, allowing a more stable interaction between Ame1^CENP-U^ and Mtw1^MIS12^ (Akiyoshi et al., 2013a; Dimitrova et al., 2016; Lang et al., 2018), was not sufficient to restore proper Mtw1-dtTomato levels to *mcm21Δ* meiotic kinetochores (Figure 7H), suggesting that other substrates may be important.

During strain construction, we have serendipitously identified an aneuploid strain carrying two copies of chromosome IX, which was able to bypass the dependence of mitotic Ctf19c^CCAN^-deficient cells on Bir1^SURVIVIN^-bound Ipl1^AURORA B^ (our observations). In that strain, Bir1^SURVIVIN^-dependent Ipl1^AURORA B^ positioning was abolished using a truncation mutant of Sli15^INCENP^, *sli15ΔN*, which lacks the Bir1^SURVIVIN^ binding site (Campbell and Desai, 2013), and which is synthetic lethal with Ctf19c^CCAN^-deficient cells (*mcm21Δ* and *ctf19Δ*) due to abolition of both mechanisms of targeting Ipl1^AURORA B^ to inner kinetochores ((Fischböck-Halwachs et al., 2019; García-Rodríguez et al., 2019)). Because *DSN1* is located on chromosome IX, we constructed a strain with an increased dosage of *DSN1* (by integration of additional *DSN1* copies at a heterologous locus), and asked whether it can restore viability of *sli15ΔN ctf19Δ* cells (Figure 7G). Remarkably, this was the case, indicating that kinetochore stabilization, albeit not solely through Dsn1 S240 and S250 residues, is likely the essential role of Ipl1^AURORA B^ targeting to inner kinetochores when Ctf19c^CCAN^ is compromised.

## DISCUSSION

Meiotic kinetochores perform a myriad of functions that are essential for healthy gamete formation, from homolog pairing and spatial regulation of meiotic recombination in prophase I, to the establishment and monitoring of oriented attachments to microtubules in metaphase I. Our global analysis of centromeric chromatin and kinetochores provides a framework for understanding how their meiosis-specific functions are executed and regulated. The resultant datasets expose the extent of kinetochore remodelling: inner kinetochore sub-complexes serve as a platform for factors that execute prophase I functions of centromeres, while assembly of the outer kinetochore at prophase exit drives attachment of fused sister kinetochores to microtubules later in meiosis I. Our approach to document chromatin composition in meiosis could be adapted to study other genetic loci of interest, such as recombination hotspots and replication origins.

### Remodelling of kinetochores for meiosis-specific functions

Our data have highlighted the re-purposing of kinetochores in meiotic prophase when microtubule-binding elements of the outer kinetochore (Ndc80c^NDC80c^ and Dam1c) are absent (Chen et al., 2020; Hayashi et al., 2006; Miller et al., 2012). Proteomics suggested that, though detectable by live-cell imaging (Meyer et al., 2015a), Spc105c^KNL1c^ is also diminished at prophase I kinetochores. This implies that loss of Ndc80c^NDC80c^ weakens Spc105c^KNL1c^ interaction with the inner kinetochore, reducing its recovery by immunoprecipitation. We find that, in the absence of the Ndc80c^NDC80c^ and Dam1c, the inner kinetochore Ctf19c^CCAN^ and central Mtw1c^MIS12c^ serve as a platform for assembly of prophase I-specific regulators. These include meiotic axis proteins, the STR resolvase and the ZMM pro-crossover and synaptonemal complex (SC) nucleation factors (Figure S4). Potentially, the centromeric localization of these factors plays a role in preventing crossover formation within pericentromeres. This would require that the pro-crossover properties of ZMMs are silenced at this region. Notably, the crossover-promoting factors MutLγ (Mlh1-Mlh3), which associate with ZMMs at presumed crossover sites on chromosome arms (Dubois et al., 2019; Duroc et al., 2017), were not found at meiotic centromeres, suggesting that regulating their recruitment may restrict crossover formation. Our data also reveal that the monopolin complex, which binds directly to the Dsn1^DSN1^ subunit of Mtw1c^MIS12c^ and directs kinetochore monoorientation during meiosis I (Plowman et al., 2019; Sarkar et al., 2013), associates with centromeres already in meiotic prophase I. Therefore, extensive re-organisation during meiotic prophase establishes kinetochore functionality that persists into the meiotic divisions.

### Central role of the Ctf19c^CCAN^ in defining meiotic kinetochores

Through a global proteomics approach and single-cell imaging, we identified a central role for the Ctf19c^CCAN^ in remodelling kinetochores for meiosis. Ctf19c^CCAN^ is critical both for preventing pericentromeric crossovers (Vincenten et al., 2015), and for maintaining cohesive linkages between sister chromatids until meiosis II (Fernius and Marston, 2009; Marston et al., 2004; Mehta et al., 2014). Targeted Ctf19c^CCAN^- dependent cohesin loading at centromeres contributes to both pericentromeric crossover suppression (Kuhl et al., 2020) and meiosis II chromosome segregation (Figure 3I-K). However, we find that defective pericentromeric cohesin cannot fully explain the profound meiotic chromosome segregation defects observed in Ctf19c^CCAN^-deficient meiosis. Indeed, multiple key regulators are lost from meiotic kinetochores lacking Ctf19c^CCAN^ components. This includes the ZMM and STR complexes, loss of which might underlie Ctf19c^CCAN^ functions in SC nucleation, non-homologous centromere coupling and prevention of pericentromeric crossover formation (De Muyt et al., 2018; Tsubouchi et al., 2008; Vincenten et al., 2015). However, future work will be required to determine whether these proteins are recruited directly by Ctf19c^CCAN^ or indirectly, for example by recognising a specific cohesin-orchestrated pericentromere structure (Paldi et al., 2020).

We also identified a distinct Ctf19c^CCAN^-dependent pathway of kinetochore assembly in meiosis. We show that in the absence of Ctf19c^CCAN^ subunits that are dispensable for viability, the Ctf19c^CCAN^, Mtw1c^MIS12c^, Ndc80c^NDC80c^, Spc105c^KNL1c^ and Dam1 complexes are all lost from meiotic centromeres, resulting in a failure of chromosomes to attach to microtubules, catastrophic segregation errors and inviable gametes. The near-complete absence of kinetochore proteins assembled on centromeric DNA in Ctf19c^CCAN^-deficient cells may also explain our previous observation that a cohesin-independent function of Ctf19c^CCAN^ prevents double strand break formation in an ∼6kb zone surrounding centromeres (Vincenten et al., 2015). Consistently, kinetochore attrition in the absence of *MCM21* or *IML3* occurs prior to double strand break formation and surprisingly early in gametogenesis. This suggests the need for continual active replenishment of kinetochores, as is observed in a wide variety of quiescent somatic as well as meiotic cells (Swartz et al., 2019), a plasticity which may facilitate the restructuring of the kinetochore for specific functions.

### A distinct kinetochore assembly pathway that is critical for meiosis

Why are meiotic kinetochores so critically dependent on the Ctf19c^CCAN^, while mitotic cells can survive in the complete absence of most of its subunits? Budding yeast kinetochores remain attached to microtubules throughout the mitotic cell cycle, except for a brief period during S phase (Kitamura et al., 2007). In contrast, meiotic kinetochores remain partially disassembled during a prolonged S phase and prophase I and are subsequently rebuilt later in meiosis. This is similar to the mammalian mitotic cell cycle, in which CCAN directs the sequential assembly of MIS12c and NDC80c as cells progress from interphase into mitosis (Musacchio and Desai, 2017). We propose that kinetochore turnover exposes a vulnerability that leads to more stringent requirements for kinetochore assembly. In the case of budding yeast meiosis, this vulnerability is manifest as a critical dependence on Ctf19c^CCAN^. This is reminiscent of situations where *de novo* kinetochore assembly is required. *Xenopus* egg extracts acutely rely on Aurora B kinase for kinetochore assembly (Bonner et al., 2019), while *de novo* assembly in yeast extracts exposed a critical requirement for Cnn1^CENP-T^, which is dispensable *in vivo* (Lang et al., 2018).

How is kinetochore remodelling and assembly achieved in meiosis? Several lines of evidence suggest a key role for Ipl1^AURORA B^ kinase which is recruited to centromeres/kinetochores in three ways: Two of these are Bir1^SURVIVIN^-dependent, through interactions with Ndc10 (in the DNA-binding Cbf3c) (Yoon and Carbon, 1999) and the pericentromeric adaptor protein Sgo1^SHUGOSHIN^ (Kawashima et al., 2007). The third involves Ctf19^CENP-P^-mediated recruitment of Ipl1^AURORA B^ to inner kinetochores, independently of Bir1^SURVIVIN^ (Fischböck-Halwachs et al., 2019; García-Rodríguez et al., 2019). Ipl1^AURORA B^ corrects improper kinetochore-microtubule attachments in mitosis and meiosis, through the Sgo1^SHUGOSHIN^-dependent recruitment pathway (Cairo et al., 2019; Meyer et al., 2013; Peplowska et al., 2014; Verzijlbergen et al., 2014). In addition, a growing list of additional Ipl1^AURORA B^ meiotic functions includes triggering outer kinetochore shedding and preventing premature spindle assembly in prophase I (Kim et al., 2013; Meyer et al., 2015b; Newnham et al., 2013). Paradoxically, in addition to these kinetochore-destabilizing activities, Ipl1^AURORA B^-dependent phosphorylation of Dsn1^DSN1^ facilitates stable kinetochore assembly (Akiyoshi et al., 2013a; Dimitrova et al., 2016). Our findings indicate that centromere-localised Ipl1^AURORA B^ becomes essential for Mtw1c^MIS12c^ recruitment and, consequently, kinetochore stability, when the Ctf19c^CCAN^ is compromised. Mitotic cells lacking non-essential Ctf19c^CCAN^ proteins are viable because Bir1^SURVIVIN^-dependent Ipl1^AURORA B^ targeting to inner centromeres compensates. In contrast, during meiosis, Bir1^SURVIVIN^-bound pool of Ipl1^AURORA B^ cannot compensate for the lack of Ctf19c^CCAN^. Whether this is due to the absence of Bir1-dependent targeting mechanisms at the critical time in gametogenesis or due to a specialised role of Ctf19^CENP-P^-positioned Ipl1^AURORA B^ in building a remodelled kinetochore specifically for meiosis, is a question for the future. Potentially, distinct kinetochore assembly mechanisms are needed to allow establishment of kinetochore monoorientation, cohesin protection or regulation of meiotic recombination.

Our data reveal that specialised assembly and preservation mechanisms underlie the unique functions of kinetochores during meiosis. We propose that such kinetochore plasticity is necessary to perform distinct functions at different stages of meiosis. Loss of kinetochore integrity may contribute to age-related decline in female fertility, where gametogenesis is protracted (Patel et al., 2015; Zielinska et al., 2015, 2019). Our comprehensive analysis of meiotic kinetochore composition provides an extensive resource for the discovery of mechanisms underlying the specialised functions of kinetochores throughout meiosis.

## Acknowledgements

We thank Angelika Amon, Sue Biggins, Christopher Campbell, Tom Ellis, Franz Herzog, Toshio Tsukiyama, Elçin Unal, Jackie Vogel and Wolfgang Zachariae for yeast strains and plasmids, and Matthias Trost for reagents. This work was performed using the Wellcome Centre for Cell Biology Bioinformatics and Proteomics core facilities, and the Centre Optical Imaging Laboratory. We are grateful to Stefan Galander, Stephen Hinshaw, Lori Koch and Gerben Vader for comments on the manuscript. This study was supported by a Wellcome Senior Research Fellowships to A.L.M [107827] and J.R. [103139], a Sir Henry Wellcome Fellowship to E.D. [096078], a Wellcome studentship to N.V. [096994], an instrument grant [108504], core funding for the Wellcome Centre for Cell Biology [092076, 203149] and NIH funding [R35GM134842] to C.L.A.

## Author contributions

Conceptualization, W.B., N.V., E.D. and A.L.M.; Methodology, W.B. and E.D.; Investigation, W.B., N.V., E.D., V.M., K.K.S. and A.L.M. Microscopy support, D.A.K.; Formal analysis, C.S. and F. L. A.; Writing, W.B. and A.L.M.; Writing – Review and Editing, all authors; Visualization, W.B. and A.L.M.; Supervision, J.R., C.L.A. and A. L. M.; Funding acquisition, J. R. and A. L. M.

## STAR methods

### RESOURCE AVAILABILITY

#### Lead Contact

Further information and request for resources and reagents should be directed to and will be filled by the Lead Contact, Adele Marston

#### Materials availability

Yeast strains and plasmids generated in this study are available without restriction through the lead contact.

#### Data availability

The mass spectrometry proteomics data have been deposited to the ProteomeXchange Consortium via the PRIDE partner repository with the dataset identifier PXD019754. Interactive volcano plots for comparison of different conditions are available for download as .html files from https://doi.org/10.7488/ds/2850.

### EXPERIMENTAL MODEL AND SUBJECT DETAILS

#### Yeast strains and plasmids

All yeast strains are either SK1 derivatives or w303 derivatives and are listed in Supplemental Table S4. Plasmids generated in this study are listed in Supplemental Table S5. Gene deletions, promoter replacements and gene tags were introduced using standard PCR-based methods, with the exception of the *CSE4-mNeonGreen, sli15ΔN* and *ndc80*(*Δ2-28*)*-GFP* alleles that were generated by CRISPR (see below). *DSN1-6His-3FLAG* (Sarangapani et al., 2014), *pCLB2-CDC20* (Lee and Amon, 2003), inducible-*ndt80 (pGAL1-NDT80*, *pGPD1-GAL4.ER*; (Benjamin et al., 2003))*, ndt80Δ* (Vincenten et al., 2015), *CEN5*-GFP dots (Toth et al., 2000), *okp1-5* (Ortiz et al., 1999)*, PDS1-tdTomato* and *HTB1-mCherry* (Matos et al., 2008), *pCUP1-IME1/pCUP1-IME4* (Chia and van Werven, 2016) and *ndc80*(*Δ2-28*) (Chen et al., 2020) were described previously.

#### CSE4-mNeonGreen

Cse4 was internally tagged with mNeonGreen by inserting the fluorescent tag flanked by two long linkers into the long N-terminal tail of Cse4, between L81 and E82. mNeonGreen flanked by linkers was amplified from AMp1604 (pFA6a-mNeonGreen-KlLEU2) using primers each with 100 bp homology to *CSE4* (AMo8738, AMo8660). Primers encoding sgRNA (AMo7441, AMo7442) allowing a Cas9 cut at *CSE4* G79 were cloned into AMp1278 (pWS082; (Shaw et al., 2019)) to produce AMp1295. The sgRNA encoding fragment was amplified from AMp1295 using primers AMo6663, AMo6664 (guide: AGCAGGTAATCTAGAAATCG). A fragment containing Cas9 and a URA marker was amplified from AMp1279 (pWS158; (Shaw et al., 2019)) with primers AMo6723, AMo6724. All three fragments were transformed into yeast and correct integrants confirmed by PCR. Primer sequences are given in Supplemental Table S6.

#### *NDC80*(*Δ2-28*)*-GFP* construction

A fragment of AMp1362 (3xV5-*NDC80Δ2-28*, *LEU2*, kind gift from Elçin Unal (Chen et al., 2020)) was amplified using primers AMo6819, AMo6853. A fragment containing Cas9 and a URA marker was amplified from AMp1279 (pWS158; (Shaw et al., 2019)) with primers AMo6723, AMo6724. Primers encoding sgRNA (AMo6847, AMo6846) allowing a Cas9 cut at *NDC80* M15 were cloned into AMp1278 (pWS082; (Shaw et al., 2019)) to produce AMp1467. The sgRNA encoding fragment was amplified from AMp1467 using primers AMo6663, AMo6664 (guide TCAACATGTGCTACATCACA). All three fragments were transformed into a strain carrying *NDC80-GFP* and correct integrants confirmed by PCR.

#### sli15ΔN construction

A fragment of AMp524 (*pGAL-GST-SLI15*, (Kim et al., 1999)) was amplified using primers AMo8665, AMo8666. A fragment containing Cas9 and a URA marker was amplified from AMp1279 (pWS158; (Shaw et al., 2019)) with primers AMo6723, AMo6724. Primers encoding two sgRNA (pair 1: AMo8661, AMo8662; pair 2: AMo8663, AMo8664) allowing a Cas9 cut at *SLI15* T194 and V129 were cloned into AMp1278 (pWS082; (Shaw et al., 2019)) to produce AMp1702 and AMp1703. The sgRNA encoding fragment were amplified from AMp1702 and AMp1703 using primers AMo6663, AMo6664 (guides: CTTCGATAACCAAACATGGG and CGCAGGAAGGAAGTCACCGA). All four fragments were transformed into a strain carrying *SLI15-3FLAG* and correct integrants confirmed by PCR.

#### Yeast carrying *CEN* and *CEN\** minichromosomes

Plasmids AMp1103 (pSB963; *CEN3*, *8xlacO*, *TRP1*) and AMp1106 (pSB972; *CEN3\**, *8xlacO, TRP1*) (Akiyoshi et al., 2009a) were amplified by PCR, digested with *EcoR*I to remove sequences required for propagation in *E. coli* (Unnikrishnan et al., 2012), re-ligated and the ∼2kb minichromosomes were transformed into haploid SK1 strains carrying integrated Stu1-cut AMp747 (pSB737; LacI-3FLAG, *URA3*; (Akiyoshi et al., 2009a). Diploids of the appropriate genotype were generated by mating.

### METHOD DETAILS

#### Yeast growth conditions

##### Meiotic Induction

To obtain meiotic cultures (apart from those used for *CEN/CEN*\* IPs, described below), yeast strains were grown at 30°C for 16 h on YPG plates (1% yeast extract, 2% bactopeptone, 2.5% glycerol, and 2% agar) and then for 8 – 24 h on YPD4% plates (1% yeast extract, 2% bactopeptone, 4% glucose, and 2% agar). Cells were then cultured in YPDA (1% yeast extract, 2% bactopeptone, 2% glucose, 0.3 mM adenine) overnight, and then inoculated at an OD_600_=0.2 – 0.5 into BYTA (1% yeast extract, 2% bactotryptone, 1% potassium acetate, 50 mM potassium phthalate) and grown overnight to an OD_600_ of ≥ 3. Cells were harvested, washed twice with an equal volume of water, resuspended into SPO medium (0.3% potassium acetate) to an OD_600_ of ≥ 1.9 and incubated at 30°C with vigorous shaking. For metaphase I arrest, a meiotic shut-off allele of *CDC20* was used (*pCLB2-CDC20*) (Lee and Amon, 2003) with harvesting of cells 8 h after resuspension in SPO medium. For prophase I arrest, *ndt80Δ* (Chu and Herskowitz, 1998) or inducible-*ndt80* was used and cells were fixed or harvested 5 – 6 h after resuspension in SPO medium. For synchronous meiosis, inducible-*ndt80* was used to allow prophase I block-release, as described by (Carlile and Amon, 2008). For synchronization at meiotic entry, *pCUP1-IME1, pCUP1-IME4* was used, as described (Berchowitz et al., 2013).

##### Cycling cells and mitotically arrested cultures for kinetochore purifications

To harvest mitotically cycling cells, overnight cultures were diluted to OD_600_=0.1, and in YPDA and harvested by centrifugation at approximately OD_600_=1.0. Cells were washed once with ice-cold dH_2_O, then washed twice with 50 mL ice-cold dH_2_O supplemented with 0.2 mM PMSF. dH_2_O supplemented with 0.2 mM PMSF was added to 15% volume of the pellet, mixed and drop frozen into liquid nitrogen before storage at −80°C.

To harvest mitotic cultures for Dsn1-6His-3FLAG immunoprecipitation, 30 mg/mL benomyl was added to 1.9 L of boiling YEP (1% yeast extract, 2% bactopeptone) and allowed to cool, before adding glucose to 2% and adenine to 0.3 mM. An overnight starter culture in YPDA was diluted into 4 x 500 mL of YPDA to OD_600_=0.3 in 4 L flasks and grown to OD_600_=1.6 – 1.8, before adding 500 mL of cooled benomyl-containing YPDA to each flask.

##### Growing strains for *CEN* and *CEN\** immunoprecipitation

Cryo-stored diploids were grown on YPG for 16 h, then inoculated into 50 mL of liquid YPDA and shaken overnight at 250 rpm at 30°C. After ∼20 – 24 h, the 50 mL of YPDA culture (OD_600_ ≥ 10) was transferred to 200 mL of -TRPA medium (see below) in a 2 L flask and shaken at ≥ 200 rpm overnight. The following day ∼2 pm, 60 mL of -TRPA culture was added to each of four 2 L flasks containing 450 mL of - TRPA medium and shaken at ≥ 200 rpm overnight at 30°C. The following morning, cultures with OD600 ≥ 5 were spun for 5 – 8 min at 4 – 5 krpm in a Beckmann centrifuge rotor 91000, and washed twice with dH_2_O at room temperature (wash 1: 1 L, wash 2: 0.5 L). The pellet was resuspended in 150 mL of SPO medium (0.3% potassium acetate), and 50 mL of this cell suspension added to each of three 4 L flasks containing 450 mL of SPO medium. Cells were then grown for 5 – 6 h, spun, drop-frozen and stored at −80°C until needed.

-TRPA medium was adapted from (Suhandynata et al., 2014) and is made by dissolving 28 g of yeast nitrogen base (Formedium) mixed with 16 g of -TRP dropout powder (Formedium, CSM -Trp +20 Ade) in 900 mL of water. Following autoclaving, 12.5 mL sterile-filtered solution of glucose (40%) is added to 0.5% final concentration, and 25 mL potassium acetate (0.8 g/mL) is added to 2% final concentration. The solution is topped-up with sterile water to 1 L.

##### Bir1-aid mitotic depletion

For depletion of Bir1^SURVIVIN^ in mitosis, the endogenous *BIR1* gene was tagged with an aid degron (**a**uxin-**i**nduced **d**egradation) (Nishimura et al., 2009). Cells were grown at room temperature overnight in liquid synthetic complete medium, then diluted to OD_600_=0.1, grown for a further 90 min, before diluting to OD_600_=0.2 and adding alpha factor (4 μg/mL) to arrest cells in G1. After 90 min, additional alpha factor was added, to 6 μg/mL final concentration. After 30 min, DMSO or NAA (5 μM) were added. Cells were transferred onto a microfluidics dish, and alpha factor was washed out to allow release into the cell cycle. Imaging commenced 30 min later as described below.

##### Induction of Clb3 expression in meiotic cells

25 μM copper sulphate was added after 3 h to meiotic cultures of wild-type and *mcm21*Δ cells harbouring *pCUP1-CLB3* allele. Two hours later, cells were released from the prophase I arrest.

#### Immunoprecipitiation

##### Preparation of anti-FLAG conjugated Dynabeads

Protein G Dynabeads (500 μL; Invitrogen) were washed twice in 1 mL 0.1M Na-phosphate, pH 7.0, before incubating with 50 μL M2 anti-FLAG monoclonal antibody (SIGMA) and 50 μL of 0.1M Na-phosphate with gentle agitation for 30 minutes at room temperature. Beads were washed twice in 1 mL of 0.1 M Na-phosphate pH 7.0 with 0.01% Tween 20, then washed twice with 1 mL of 0.2 M triethanolamine, pH 8.2. Antibody-conjugated Dynabeads were resuspended in 1 mL of 20 mM DMP (Dimethyl Pimelimidate, D8388, Sigma) in 0.2 M triethanolamine, pH 8.2 (prepared immediately before use) and incubated with rotational mixing for 30 min at room temperature. Beads were concentrated, the supernatant removed and 1 mL of 50 mM Tris-HCl, pH 7.5 added before incubating for 15 minutes with rotational mixing. The supernatant was removed and beads were washed three times with 1 mL 1XPBST+0.1% Tween-20 before resuspending in 300 mL of 1xPBST.

##### Immunoprecipitation of *CEN/CEN\** chromatin

Yeast cells were pulverised mechanically using a Retsch RM100 mortar-grinder. Approximately 20 g of cryogrindate was used per experiment. Cryogrindates were resuspended in H_0.15_ buffer (25 mM Hepes (pH 8.0), 2 mM MgCl_2_, 0.1 mM EDTA (pH 8.0), 0.5 mM EGTA-KOH (pH 8.0), 15% glycerol, 0.1% NP-40, 150 mM KCl) supplemented with phosphatase inhibitors (2 mM β-glycerophosphate, 1 mM Na_4_P_2_O_7_, 5 mM NaF, 0.1 mM Na_3_VO_4_), protease inhibitors (2 mM final AEBSF, 0.2 μM microcystin and 10 μg/mL each of ‘CLAAPE’ protease inhibitors (chymostatin, leupeptin, antipain, pepstatin, E64)) and 1 mM N-Ethylmaleimide (NEM) at 1 g of grindate to 1.5 mL of complete buffer ratio. Debris was removed by centrifugation (1×5 min at 5 krpm, 1×15 min at 5 krpm) and lysates incubated at 4°C for 3 h with Protein G Dynabeads previously conjugated to mouse anti-Flag (M2, Sigma) with DMP (Dimethyl Pimelimidate, D8388, Sigma). 12.5 μL of bead suspension and 5.8 μL of antibody were used per 1 g of grindate. Beads were washed three times in H_0.15_ buffer before sequential elution at 37°C for 2×30 min in 1% Rapigest (Waters).

##### Immunoprecipitation of Dsn1-6His-3FLAG

Yeast cell pulverization and immunoprecipitation was performed as above except that: (1) H_0.15_ buffer was additionally supplemented with 1 mM benzamidine, and one tablet of EDTA-free protease inhibitor tablet (Roche) per every 25 mL. (2) phosphatase inhibitors concentrations were doubled. (3) for elution, 0.5 mg/mL FLAG peptide in lysis buffer was added to beads, gently mixed and incubated at room temperature for 20 min. Beads were concentrated on a magnet, the supernatant removed and snap-frozen in liquid nitrogen before preparation for mass spectrometry as below.

##### Mass spectrometry

Protein samples were briefly run into an SDS-PAGE gel (NuPAGE Novex 4-12% Bis-Tris gel, ThermoFisher, UK), in NuPAGE buffer (MES) and visualised using Instant*Blue*^TM^ stain (Sigma Aldrich, UK). The stained gel areas were excised and de-stained with 50 mM ammonium bicarbonate (Sigma Aldrich, UK) and 100% v/v acetonitrile (Sigma Aldrich, UK) and proteins were digested with trypsin (Shevchenko *et al,* 1996). In brief, proteins were reduced in 10 mM dithiothreitol (Sigma Aldrich, UK) for 30 min at 37°C and alkylated in 55 mM iodoacetamide (Sigma Aldrich, UK) for 20 min at ambient temperature in the dark. They were then digested overnight at 37°C with 12.5 ng/μL trypsin (Pierce, UK). Following digestion, samples were diluted with equal volume of 0.1% TFA and spun onto StageTips (Rappsilber et al., 2003). Peptides were eluted in 40 μL of 80% acetonitrile in 0.1% TFA and concentrated down to 1 μL by vacuum centrifugation (Concentrator 5301, Eppendorf, UK). Samples were then prepared for LC-MS/MS analysis by diluting them to 5 μL with 0.1% TFA.

For *CEN-* and *CEN\**-chromatin samples, as well as for the no tag sample in Dsn1-6His-3FLAG immunoprecipitation, LC-MS-analyses were performed on an Orbitrap Fusion™ Lumos™ Tribrid™ Mass Spectrometer (Thermo Fisher Scientific, UK) coupled on-line, to an Ultimate 3000 RSLCnano Systems (Dionex, Thermo Fisher Scientific, UK). Peptides were separated on a 50 cm EASY-Spray column (Thermo Fisher Scientific, UK) assembled in an EASY-Spray source (Thermo Fisher Scientific, UK) and operated at a constant temperature of 50°C. Mobile phase A consisted of 0.1% formic acid in water while mobile phase B consisted of 80% acetonitrile and 0.1% formic acid. Peptides were loaded onto the column at a flow rate of 0.3 μL/min and eluted at a flow rate of 0.2 μL/min according to the following gradient: 2 to 40% buffer B in 150 min, then to 95% in 11 min. Survey scans were performed at 120,000 resolution (scan range 350-1500 m/z) with an ion target of 4E5. MS2 was performed in the Ion trap at rapid scan mode with ion target of 2E4 and HCD fragmentation with normalised collision energy of 27 (Olsen et al., 2007). The isolation window in the quadrupole was set at 1.4 Thomson. Only ions with charge between 2 and 7 were selected for MS2.

For the Dsn1-6His-3FLAG immunoprecipitation samples (except the no tag sample which was processed as described above), MS-analyses were performed on a Q Exactive mass spectrometer (Thermo Fisher Scientific, UK), coupled on-line to Ultimate 3000 RSLCnano Systems (Dionex, Thermo Fisher Scientific). The analytical column with a self-assembled particle frit (Ishihama et al., 2002) and C18 material (ReproSil-Pur C18-AQ 3 μm; Dr. Maisch, GmbH) was packed into a spray emitter (75-μm ID, 8-μm opening, 300-mm length; New Objective, UK) using an air-pressure pump (Proxeon Biosystems, UK). Mobile phase A consisted of water and 0.1% formic acid; mobile phase B consisted of 80% acetonitrile and 0.1% formic acid. Peptides were loaded onto the column at a flow rate of 0.5 μL/min and eluted at a flow rate of 0.2 μL/min according to the following gradient: 2 to 40% buffer B in 180 min, then to 95% in 16 min.

The resolution for the MS1 scans was set to 70,000 and the top 10 most abundant peaks with charge ≥ 2 and isolation window of 2.0 Thomson were selected and fragmented by higher-energy collisional dissociation (Olsen et al., 2007) with normalised collision energy of 30. The maximum ion injection time for the MS and MS2 scans was set to 20 and 60 ms respectively and the AGC target was set to 1E6 for the MS scan and to 5E4 for the MS2 scan. Dynamic exclusion was set to 60 s.

The MaxQuant software platform version 1.6.1.0 (released in April 2018) was used to process raw files and searches were conducted against the *Saccharomyces cerevisiae* (strain SK1) complete/reference proteome set of the *Saccharomyces* Genome Database (released in May, 2019), using the Andromeda search engine (Cox *et al*, 2011). The first search peptide tolerance was set to 20 ppm, while the main search peptide tolerance was set to 4.5 pm. Isotope mass tolerance was set to 2 ppm and maximum charge to 7. Maximum of two missed cleavages were allowed. Carbamidomethylation of cysteine was set as fixed modification. Oxidation of methionine and acetylation of the N-terminal as well as phosphorylation of serine, threonine and tyrosine were set as variable modifications. LFQMS analysis was performed by employing the MaxLFQ algorithm) (Cox et al., 2014). For peptide and protein identifications FDR was set to 1%.

##### Quantitative analysis of mass spectrometry data

LFQMS data was processed using Bioconductor ***DEP*** R package (Zhang et al., 2018). Briefly, proteins with indicators Reverse “+” and Potential.contaminant “+” were removed from the dataset. Data were filtered to only keep proteins detected in all replicates of at least one condition, LFQ intensities were log_2_ transformed and normalised using variance-stabilised normalisation. Then, imputation was performed using “MinProb” function, with q=0.001. Log_2_(Fold Change) and p values were calculated using linear models and empirical Bayes method.

Pie charts in Figure 1 b were generated by identifying proteins enriched over no tag control with p value < 0.01 and Log_2_(Fold Change) > 4. A single no tag sample was used for the kinetochore sample. Three previously published metaphase I samples were used as no tag control for *CEN* and *CEN*\* chromatin samples (PRIDE identifier: PXD012627, samples Sgo1_no_tag_1-3, (Galander et al., 2019))).

Heatmaps in Figures 1, 2A and S1 were generated using ***DEP*** package plot_heatmap() function with a modified colour scheme. To generate the heatmap shown in Figure 2, the *CEN/CEN*\* ratio was determined for each protein in each condition. Data were filtered to reject proteins that failed to show *CEN/CEN*\* ratio > |2| and p value < 0.05, which we defined as significant enrichment, in at least one condition.

Cumulative plots shown in Figure 4A were generated using DEP-processed data. Following filtering, normalisation and imputation, the log_2_ transformed data were exponentiated to obtain LFQ intensities. These were then summed for individual complexes in each condition, log_2_ transformed and ratios between conditions were determined and plotted. Complexes were defined as follows: **<u>Cbf3 complex</u>**: Skp1, Cbf2, Cep3, Ctf13; **<u>Cse4, Mif2</u>**: Cse4, Mif2; **<u>Ctf19 complex</u>**: Ame1, Okp1, Chl4, Nkp1, Mcm22, Mcm16, Nkp2, Ctf3, Ctf19, Wip1, Cnn1 ; **<u>Mtw1 complex</u>**: Mtw1, Nnf1, Nsl1, Dsn1; **<u>Outer KT</u>**: Spc105, Kre28, Dad3, Dad1, Dad4, Spc19, Duo1, Dam1, Ask1, Hsk3, Spc34, Dad2, Ndc80, Nuf2, Spc24, Spc25; **<u>SPBs and MTs</u>**: Spc72, Spc110, Spc42, Cnm67, Spo21, Ady3, Nud1, Mpc54, Don1, Mps2, Spc97, Spc98, Tub2, Stu1, Stu2, Tub3, Tub1, Bik1, Cin8, Ase1, Tub4; **<u>cohesin</u>**: Smc1, Smc3, Irr1, Rad61, Pds5, Rec8; **<u>Msh4/Msh5</u>**: Msh4, Msh5; **<u>SZZ</u>**: Spo16, Spo22, Zip2.

The LFQMS data used in Figures 1, 2, S1, S2 utilises both haploid and diploid cycling cells samples, and, following initial analyses, some diploid samples were rejected from the original dataset. For LFQMS data shown in all other figures, only haploid cycling cells were rejected, as no haploid samples were obtained for *iml3Δ* and *mcm21Δ* cells (See Supplemental Table S4).

##### Chromatin immunoprecipitation

Cells in 50 mL of SPO culture at OD_600_≥1.9 were fixed by addition of formaldehyde to 1%. Following 2 h crosslinking, cultures were spun, supernatant was removed and the pellet washed twice in 10 mL of ice-cold TBS (20 mM Tris/HCl at pH 7.5, 150 mM NaCl) and once in 10 mL of ice-cold FA lysis buffer (50 mM HEPES-KOH at pH 7.5, 150 mM NaCl, 1 mM EDTA, 1% v/v Triton X-100, 0.1% w/v Sodium Deoxycholate) with 0.1% w/v SDS and the snap-frozen pellet was kept at −80°C. Next, the pellet was resuspended in 0.4 mL of ice-cold FA buffer supplemented with EDTA-free protease inhibitors (Roche), 1 mM PMSF (=1XcFA) and 0.5% w/v SDS. Cells were lysed using silica beads (Biospec Products) in a Fastprep Bio-pulverizer FP120, with two 30 s rounds of bead-beating at maximum power, with intervening 10 min incubation on ice. The lysate was collected and spun for 15 min at 14 krpm, supernatant was rejected, and the pellet was washed with 1 mL of 1XcFA supplemented with 0.1% w/v SDS. Following another spin for 15 min at 14 krpm, the pellet was resuspended in 0.5 mL of 1XcFA supplemented with 0.1% w/v SDS, and sonicated using BioRuptor Twin sonicating device (Diagenode) at HIGH setting, 30 x 30 s at 4°C. Extract was spun for 15 min at 14 krpm, supernatant recovered, an additional 0.5 mL of 1XcFA supplemented with 0.1% w/v SDS was added, and the mixture was spun again for 15 min at 14 krpm. 1 mL of supernatant was then added to a fresh tube containing 0.3 mL of 1XcFA supplemented with 0.1% w/v SDS. From this solution, 1 mL was used for the IP and 100 μL was stored at −20°C as input sample. Protein G Dynabeads (Invitrogen) were washed four times in 1 mL of ice-cold 1XcFA lysis buffer with 0.1% w/v SDS. For the IP, 15 μL of pre-washed Dynabeads as well as the appropriate amount of antibody (mouse anti-Ha (12CA5, Roche, 7.5 μL), mouse anti-Flag (M2, Sigma, 5 μL)) were added to 1 mL of lysate and incubated overnight at 4°C. Next, the supernatant was removed and the Dynabeads were incubated in 1 mL of ChIP wash buffer 1 (0.1% w/v SDS, 275 mM NaCl, 1XFA) with rotational mixing for 5 min at room temperature. This washing was repeated with ChIP wash buffer 2 (0.1% SDS, 500 mM NaCl, 1XFA), ChIP wash buffer 3 (10 mM Tris/HCl at pH 8, 0.25 M LiCl, 1 mM EDTA, 0.5% v/v NP-40, 0.5% w/v Sodium Deoxycholate), and TE (10 mM Tris/HCl at pH 8, 1 mM EDTA). 200 μL of 10% w/v Chelex (Biorad) suspension in DEPC-treated sterile water (VWR) was added to the Dynabeads as well as to 10 μL of the thawed input sample. This was incubated for 10 minutes at 100°C, cooled and 2.5 μL of 10 mg/mL Proteinase K (Promega) was added. Incubation at 55°C for 30 minutes was followed by incubation for 10 minutes at 100°C and cooling samples on ice. Samples were spun and 120 μL of supernatant of both IP and input samples was collected. qPCR was performed as described in.(Verzijlbergen et al., 2014). Mean values are shown from a minimum of 3 biological repeats, with error bars representing standard error. Primers for qPCR analysis are listed in Supplemental Table S6.

##### Western blotting

For western immunoblotting, samples were fixed in trichloroacetic acid for 10 min, acetone-washed and whole cell extracts prepared by bead-beating in TE-containing protease inhibitors before transferring to nitrocellulose membrane. Antibodies used were rabbit anti-Pgk1 (lab stock,1:50000), mouse anti-Myc (Covance/Biolegend 9E10, 1:1000), mouse anti-Ha (Mono HA.11, Covance, 1:1000), sheep anti-mouse-HRP (GE Healthcare, 1:5000), donkey anti-rabbit-HRP (GE Healthcare, 1:10000).

##### Viability assays

Viability of mitotically cycling cells was determined by growing the cells to OD_600_=1, and diluting them 1000 times, then plating 400 μL of cell suspension onto YPDA plates. Viability of meiotic cells was determined by dissecting 36 or more tetrads of a homozygous diploid carrying the mutation of interest. Viability drop following return to growth was determined by growing cells as described in “Meiotic induction” up until cells were moved into SPO medium. Then, for each SPO culture, 300 cells were plated at t = 0 h and t = 5 h. Cells were counted two days after, ratio 5h/0h was determined, and viability drop was calculated: viability drop = (1 – (5h/0h)) * 100%.

##### Chromosome segregation assay

Diploid strains with either one copy (heterozygous) or both copies (homozygous) of chromosome V marked with GFP were induced to sporulate at 30°C. To score GFP dots, cells were fixed as previously described (Fernius and Marston, 2009) and for each biological repeat 100 tetranucleate cells were counted at 8 or 10 hours after transfer to sporulation medium.

##### Live-cell imaging

###### Equipment

Live-cell imaging quantified in Figure 7F was performed at ambient temperature on a DeltaVision Elite system (Applied Precision) connected to an inverted Olympus IX-71 microscope with a 100x UPlanSApo NA 1.4 oil lens. Images were taken using a Photometrics Cascade II EMCCD camera. The Deltavision system was controlled using SoftWoRx software. Live-cell imaging shown in Figures 3C-D, 3G, 4B-C, 4F-G, 5D-G, 6E-F, 6H-I, 7B-E, 7H, S6A and S6C was performed at 30°C on a Zeiss Axio Observer Z1 (Zeiss UK, Cambridge) equipped with a Hamamatsu Flash 4 sCMOS camera, Prior motorised stage and Zen 2.3 acquisition software. Live-cell imaging shown in Figures 6D and 5A-C (all at 25°C) was done using spinning-disk confocal microscopy employing a Nikon TE2000 inverted microscope with a Nikon X100/1.45 NA PlanApo objective, attached to a modified Yokogawa CSU-10 unit (Visitech) and an iXonþ Du888 EMCCD camera (Andor), controlled by Metamorph software (Molecular Devices).

###### Imaging chamber preparation

For the experiment shown in Figure 7B-E and S6C, cells were imaged at 30°C on an ONIX microfluidic perfusion platform by CellASIC. Cells were transferred to a microfluidics plate 30 min before alpha factor release. For imaging shown in Figures 7F and 5D cells were placed on glass slides before taking still images. For all other experiments, cells were imaged at the indicated temperature in a 4-well or 8-well Ibidi glass-bottom dishes.

###### Growth conditions

Meiotic cultures were pre-grown in culture flasks for ∼3 h (*pCLB2-CDC20* and asynchronous) or about 4.5 h (inducible-*ndt80*) before transfer to concanavalin A-coated dishes, where they were left to attach for 20 – 30 min. Imaging began about 30 min after attachment was completed, with images being acquired every 7.5 – 15 min for 10 – 12 h.

###### Image acquisition and analysis

Eight to eleven z sections were acquired with 0.6 – 0.8 µm spacing. For prophase block-release experiments, beta-estradiol was added immediately before the first image was acquired. Images were analysed using ImageJ (National Institutes of Health). Final image assembly was carried out using Adobe Photoshop and Adobe Illustrator. Where signal intensity was measured (Figures 4C, 7C-D, 3H and S6C), a circular region was drawn that encompassed the region of interest (ROI), and mean ROI intensity was measured. The same size region was then drawn in an area in the vicinity, and the mean intensity of this area was measured and defined as background intensity (bROI). The signal presented in figures is the mean ROI signal minus mean bROI signal. If two foci were measured per cell (Figures 7C and 3H), an average of mean intensity of both ROIs minus an average of both bROIs was calculated. Spindle length in Figure 5G was measured 75 min after the emergence of meiotic spindle or at the last time point before spindle collapse, if it happened before 75 min.

##### Quantification and statistical analysis

R software was used for statistical analysis. Statistical details can be found in figure legends, apart from details about analysis of LFQMS data presented in Figures 1C, 2A-C, 4A, S1, S2, S3, S4, which are described in “Quantitative analysis of mass spectrometry data” part of the Methods section. Details about imaging quantification can be found in “Image acquisition and analysis” part of the Methods section. Following quantification of imaging data, their normality was tested, which guided selection of appropriate statistical test.

## Supplementary Information

Supplementary information consists of 6 supplementary tables and 7 supplemental figures

**Figure S1, related to Figure 2.**
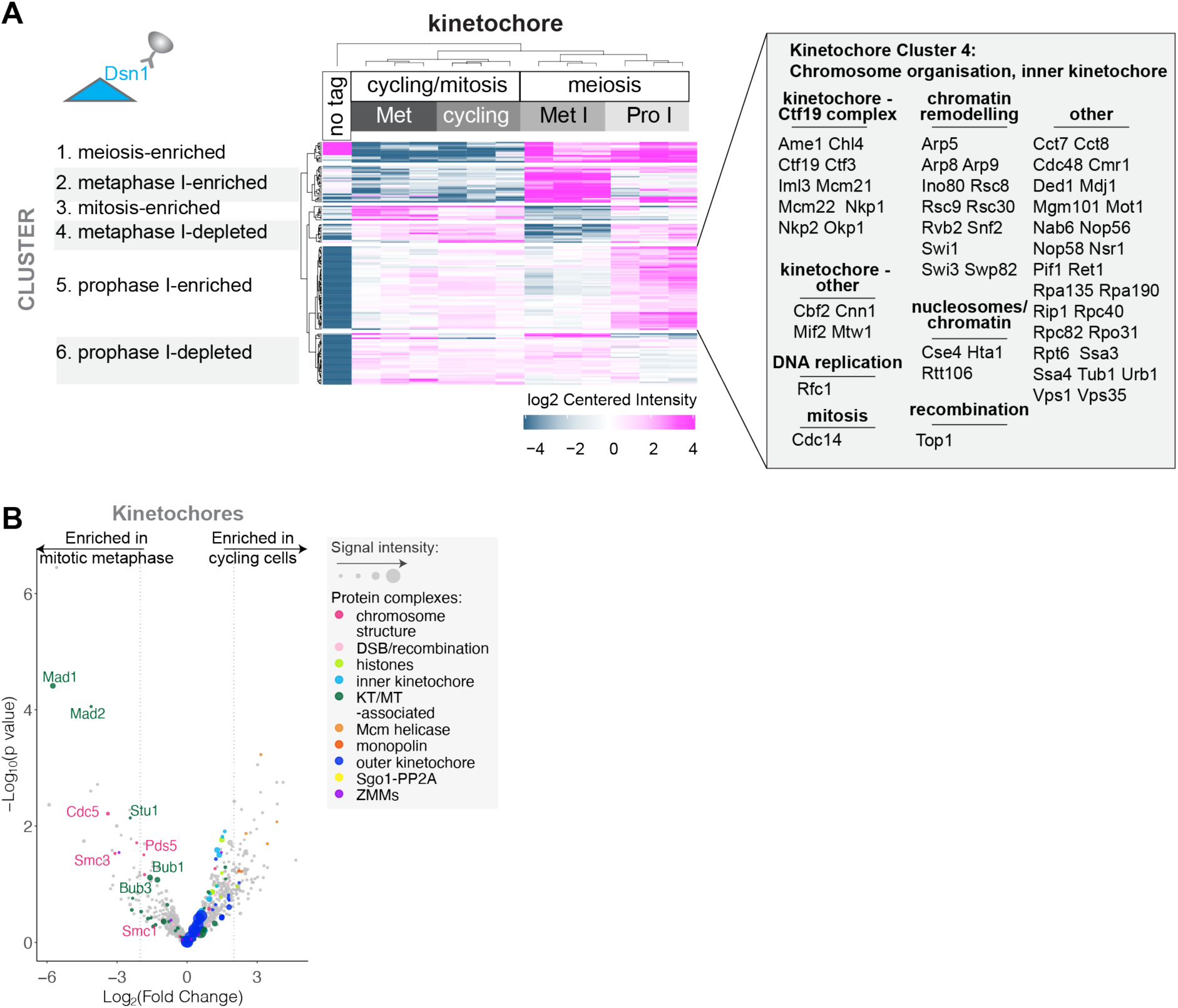
Comparison of mitotic and meiotic kinetochore proteomes. (A) Kinetochore composition varies across cell cycle stages. Clustering analysis of kinetochore samples (k-means). Inset lists proteins present in a cluster containing prophase I-enriched proteins. A cut-off of Log_2_(Fold Change) > 1 and p < 0.05 was used. (B) Kinetochore-associated proteome of cycling cells is similar to that of mitotic metaphase-arrested cells. Volcano plot showing LFQMS-identified proteins co-purifying with Dsn1-6His-3FLAG in cycling cells vs. mitotic metaphase-arrested (benomyl) cells. Log_2_(Fold Change) between conditions are shown with their corresponding p values (see methods). Dashed line indicates Log_2_(Fold Change) = |2|. See also Supplemental Table S2.

**Figure S2, related to Figure 2.**
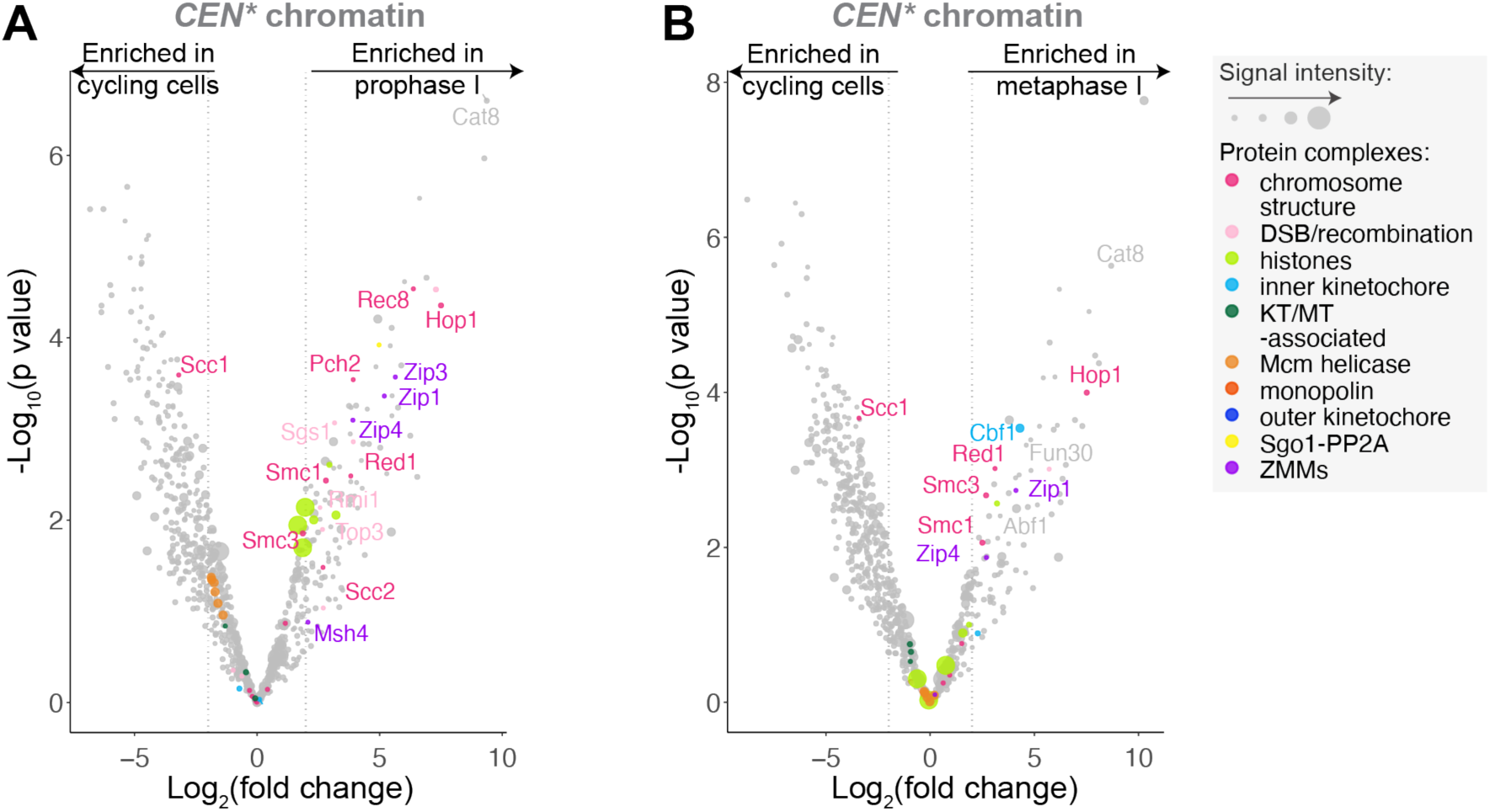
Comparison of *CEN\** chromatin in meiotic prophase and metaphase I compared to cycling cells. (A and B) Comparison of *CEN\** chromatin isolated from cycling cells and two meiotic stages. Volcano plots showing LFQMS-identified proteins co-purifying with *CEN\** plasmids immunopurified from cells arrested in prophase (*ndt80Δ,* A) and metaphase I (*pCLB2-CDC20,* B) as compared to cycling cells. Log_2_(Fold Change) between conditions are presented with their corresponding p values (see methods). Dashed line indicates Log_2_(Fold Change) = |2|.

**Figure S3, related to Figure 2.**
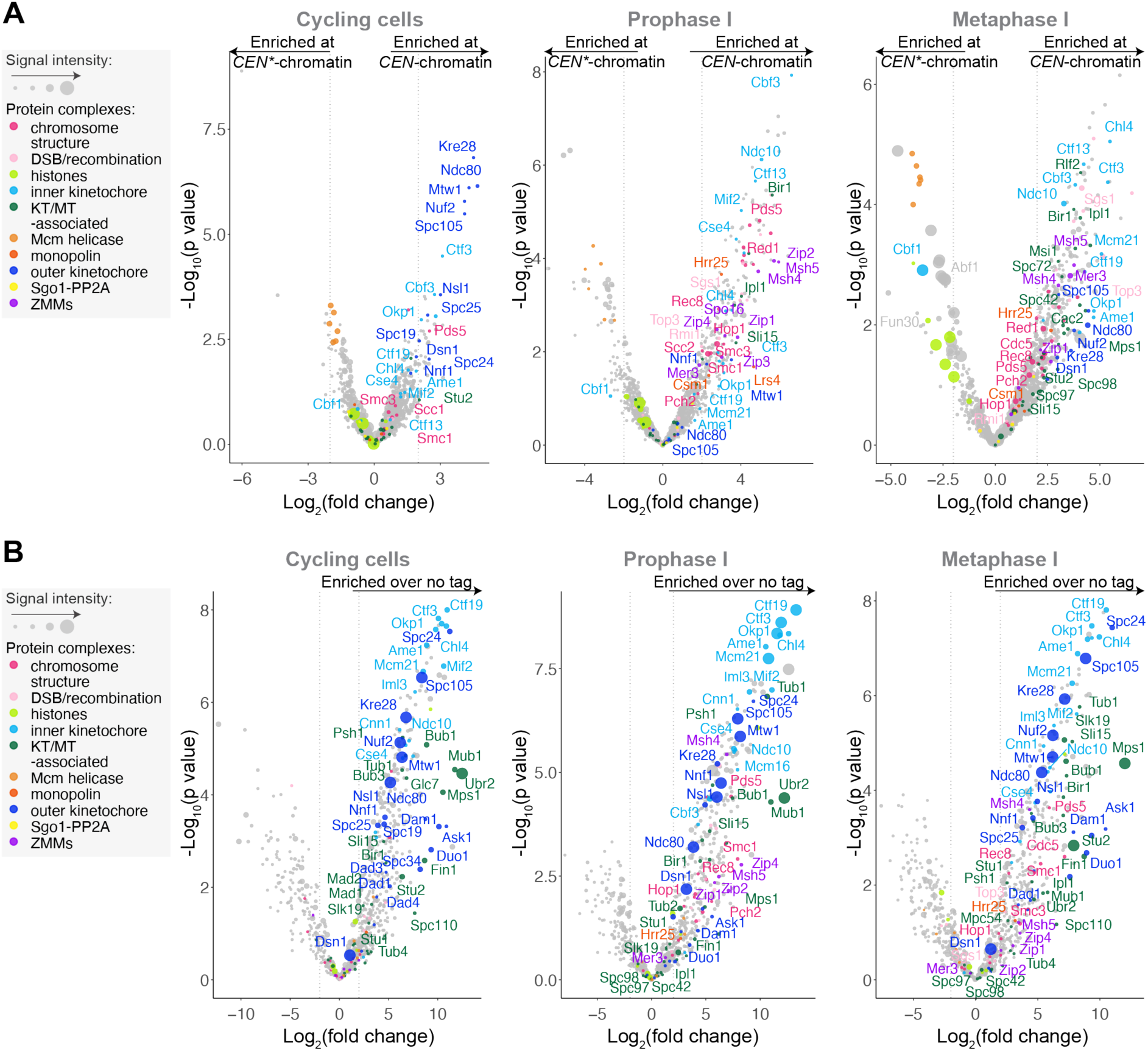
Overview of protein enrichment in *CEN* and kinetochore purifications from cycling, prophase I and metaphase I cells. (A and B) CEN chromatin (A) and kinetochore (B) composition varies depending on cell cycle stage. (A) Volcano plots showing LFQMS-identified proteins co-purifying with *CEN* and *CEN\** plasmids immunopurified from cells that are cycling, arrested in prophase (*ndt80*Δ) and metaphase I (*pCLB2-CDC20*). (B) Volcano plot showing LFQMS-identified proteins co-purifying with Dsn1-6His-3FLAG immunopurified from cells that are cycling, arrested in prophase (inducible-*ndt80)* and metaphase I (*pCLB2-CDC20*).

**Figure S4, related to Figure 4.**
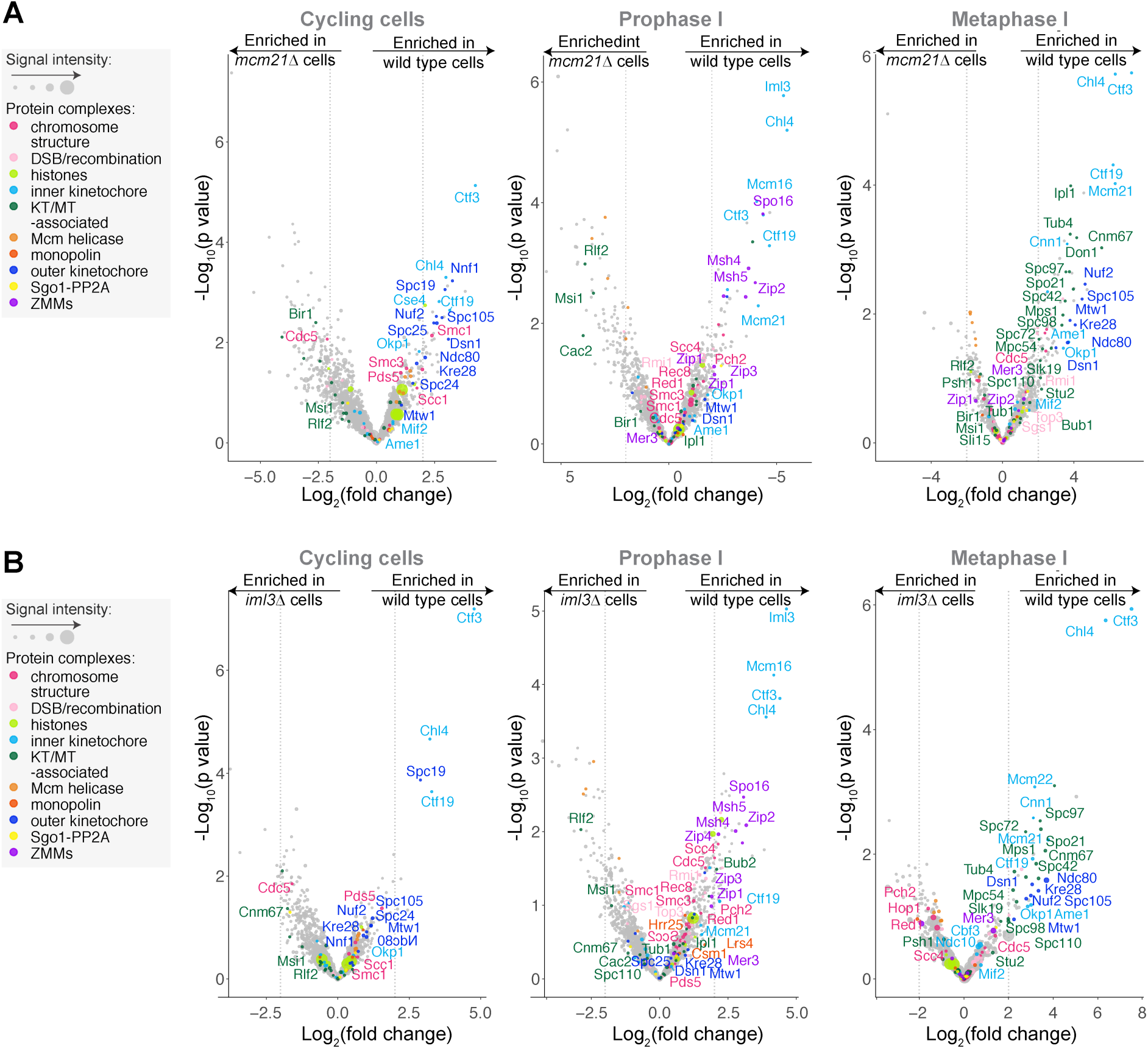
Kinetochores fail to assemble in meiosis in *iml3Δ* and *mcm21Δ* cells. (A and B) The Ctf19c^CCAN^ plays a central role in kinetochore composition during meiosis. (A) Volcano plots showing LFQMS-identified proteins co-purifying with *CEN* chromatin in wild-type vs. *mcm21*Δ cells from mitotically cycling cells, meiotic prophase I and meiotic metaphase I cells. Log_2_(Fold Change) between conditions are presented with their corresponding p values (see methods). Dashed line indicates Log_2_(Fold Change) = |2|. Plots show data for cycling cells, cells arrested in prophase I (*ndt80Δ*) allele and cells arrested in metaphase I (*pCLB2-CDC20*) allele. (B). Volcano plots showing the LFQMS-identified proteins co-purifying with *CEN* chromatin in wild-type vs. *iml3Δ* cells at different cell cycle stages. Log_2_(Fold Change) between conditions are presented with their corresponding p values (see methods). Dashed line indicates Log_2_(Fold Change) = |2|. Plots show data for cycling cells, cells arrested in prophase I (*ndt80Δ*) allele and cells arrested in metaphase I (*pCLB2-CDC20*) allele.

**Figure S5, related to Figure 5:**
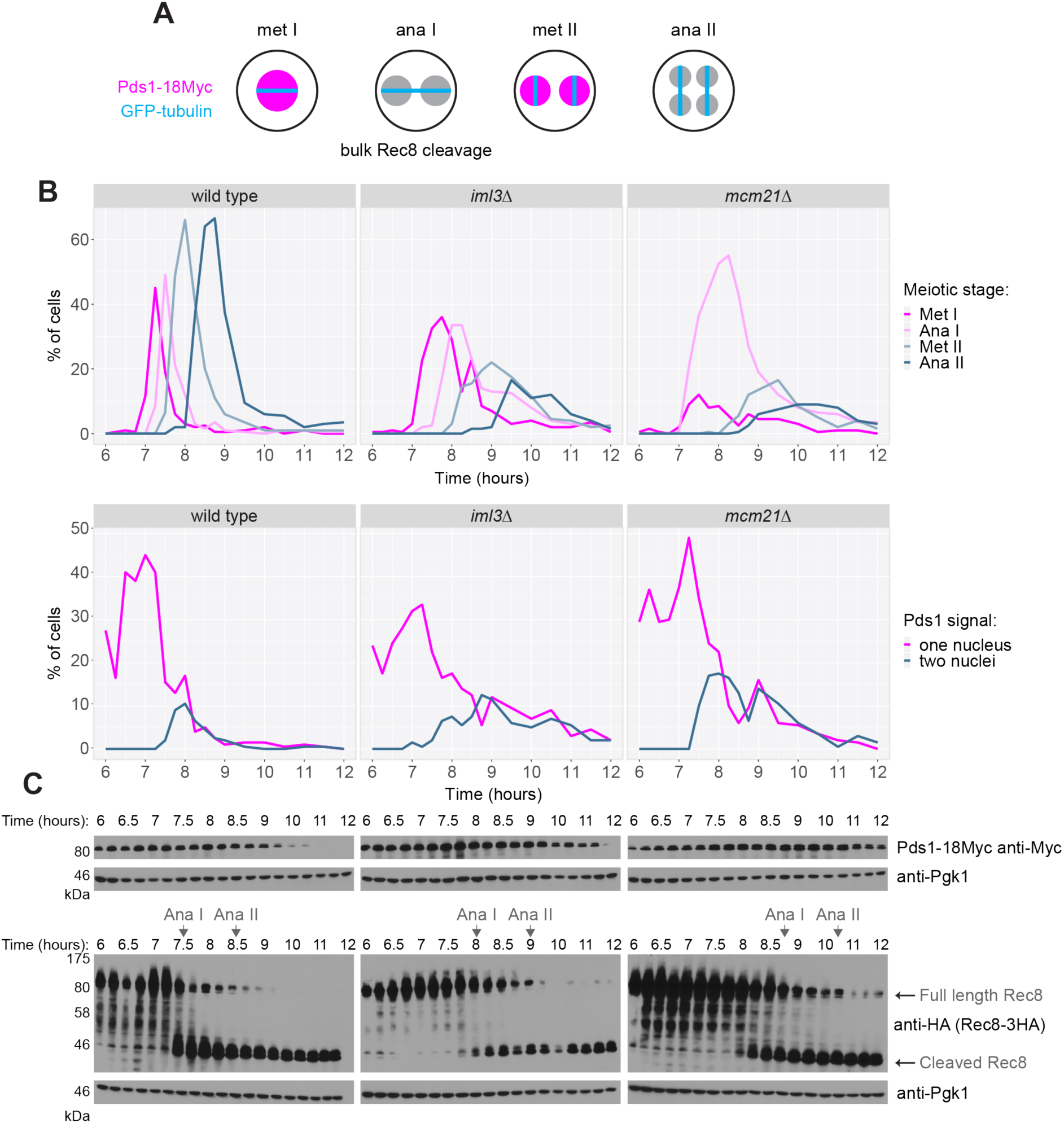
Ctf19 complex deletion mutants show a metaphase I delay. (A-C) Cells lacking *IML3* and *MCM21* are delayed in formation of meiosis II spindles, cleavage of cohesin (Rec8^REC8^) and degradation of securin (Pds1^SECURIN^). (A) Schematic of the experiment shown in B and C. The expected spindle phenotypes, the presence or absence of Pds1^SECURIN^, and the presence or absence of cleaved Rec8^REC8^ are indicated for the different cell cycle stages. (B and C) Wild-type, *iml3Δ* and *mcm21Δ* cells were synchronously released from prophase I and samples were collected at indicated times. (B) Spindle morphology and the presence of Pds1^SECURIN^ were scored by immunofluorescence. (C) Anti-Myc (Pds1-18Myc), anti-HA (Rec8-3HA) and anti-Pgk1 western immunoblots. Arrows (Ana I and Ana II) represent the onset of anaphase I and anaphase II, based on Rec8 cleavage, respectively.

**Figure S6, related to Figure 7.**
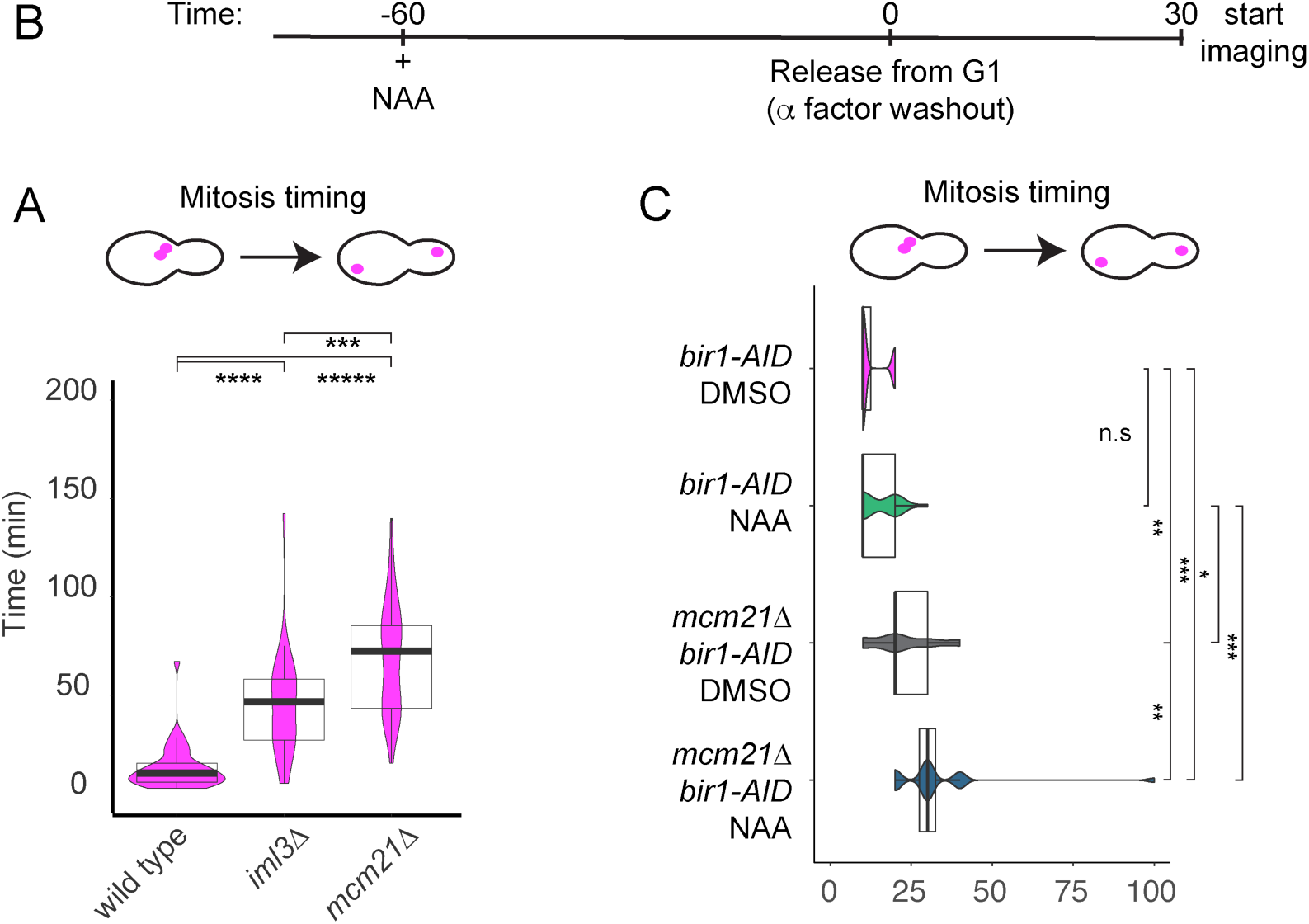
Interplay between the Ctf19 complex and Ipl1 for kinetochore integrity. (A) Cells lacking *IML3* and *MCM21* show delayed cell cycle progression. Asynchronously growing wild-type, *iml3*Δ and *mcm21*Δ cells expressing Mtw1-tdTomato and Ndc80-GFP were imaged. Time between emergence of bilobed kinetochore structure and anaphase (when two Mtw1-tdTomato foci reach opposite ends of mother and daughter cells) was measured. ***p < 10^-4^, ****p < 10^-8^, *****p < 10^-15^; Mann-Whitney test. n > 30 cells. (B) Schematic of the experiment shown in C. (C) Loss of *MCM21* function and Ndc10-localised Ipl1^AURORA B^ at kinetochores has an additive effect extending metaphase duration. Further analysis of the experiment shown in Figure 7A-D. Time between emergence of bilobed kinetochore structure and anaphase (when two Mtw1-tdTomato foci reach opposite ends of mother and daughter cells) was measured. *p < 0.05, *p < 0.01, ***p < 10^-5^; Mann-Whitney test. n > 19 cells.

**Supplemental Table S1**. Protein groups from clustering analysis of *CEN* and *CEN\** chromatin.

**Supplemental Table S2**. Protein groups from clustering analysis of the kinetochore datasets.

**Supplemental Table S3.** Protein groups after clustering analysis of *CEN/CEN\** chromatin

**Supplemental Table S4.** Yeast strains used in this study

**Supplemental Table S5.** Plasmids used in this study

**Supplemental Table S6.** Primers used in this study.

## Notes

https://doi.org/10.7488/ds/2850

### Competing Interest Statement

The authors have declared no competing interest.

