## Supplemental Table S4 for "The proteomic landscape of centromeric chromatin reveals an essential role for the Ctf19^CCAN^ complex in meiotic kinetochore assembly"

Table S4. Yeast strains used in this study

| **Strain number** | **Relevant genotype** | **Yeast** | **In figures:** |
| --- | --- | --- | --- |
| AM1603 | MATa/alpha /promURA3::TetR::GFP::LEU2 tetOx224-URA3 | SK1 | 6c |
| AM1835 | MATa/alpha | SK1 | 3a, 3b |
| AM3560 | MATa/alpha cdc20::pCLB2-3HA-CDC20::KanMX6/cdc20::pCLB2-3HA-CDC20::KanMX6 | SK1 | 6b |
| AM3722 | MATa/alpha ctf19Δ::KanMX6/ctf19Δ::KanMX6 | SK1 | 3a, 3b |
| AM4014 | MATa/alpha ndt80∆::LEU2/ndt80∆::LEU2 | SK1 | 6g |
| AM6142 | MATa/alpha ndt80::pGAL-NDT80::TRP1/ndt80::pGAL-NDT80::TRP1 ura3::pGPD1-GAL4(848).ER::URA3/ura3::pGPD1-GAL4(848).ER::URA3 ubr1Δ::KanMX4/ubr1Δ::KanMX4 REC8-3HA::URA3/ REC8-3HA::URA3 PDS1-18MYC::LEU2/PDS1-18MYC::LEU2 | SK1 | s5b, s5c |
| AM8291 | MATa/alpha cdc20::pCLB2-3HA-CDC20::KanMX6/cdc20::pCLB2-3HA-CDC20::KanMX6 DSN1-6HIS-3FLAG::URA3/DSN1-6HIS-3FLAG::URA3 | SK1 | 1b,2c, s1a, s1b, s3b |
| AM8292 | MATa/alpha cdc20::pCLB2-3HA-CDC20::KanMX6/cdc20::pCLB2-3HA-CDC20::KanMX6 DSN1-6HIS-3FLAG::URA3/DSN1-6HIS-3FLAG::URA3 | SK1 | 5i, 5j |
| AM8382 | MATa/alpha ndt80::pGAL-NDT80::TRP1/ndt80::pGAL-NDT80::TRP1 ura3::pGPD1-GAL4(848).ER::URA3/ura3::pGPD1-GAL4(848).ER::URA3 ubr1Δ::KanMX4/ubr1Δ::KanMX4 REC8-3HA::URA3/ REC8-3HA::URA3 PDS1-18MYC::LEU2/PDS1-18MYC::LEU2 iml3Δ::KanMX6/iml3Δ::KanMX6 | SK1 | s5b, s5c |
| AM8769 | MATa/alpha GAL-NDT80::TRP1/GAL-NDT80::TRP1 ura3::pGPD1-GAL4(848).ER::URA3/ura3::pGPD1-GAL4(848).ER::URA3 NDC10-6HA::HIS3MX6/NDC10-6HA::HIS3MX6 | SK1 | 6g |
| AM8805 | MATa/alpha ndt80::pGAL-NDT80::TRP1/ndt80::pGAL-NDT80::TRP1 ura3::pGPD1-GAL4(848).ER::URA3/ura3::pGPD1-GAL4(848).ER::URA3 ubr1Δ::KanMX4/ubr1Δ::KanMX4 REC8-3HA::URA3/ REC8-3HA::URA3 PDS1-18MYC::LEU2/PDS1-18MYC::LEU2 mcm21Δ::KanMX6/mcm21Δ::KanMX6 | SK1 | s5b, s5c |
| AM8825 | MATa/alpha cdc20::pCLB2-3HA-CDC20::KanMX6/cdc20::pCLB2-3HA-CDC20::KanMX6 his3::HIS3p-GFP-TUB1-HIS3/his3::HIS3p-GFP-TUB1-HIS3 MTW1-tdTomato::NAT/MTW1-tdTomato::NAT | SK1 | 5f, 5g |
| AM8861 | MATa/alpha GAL-NDT80::TRP1/GAL-NDT80::TRP1 ura3::pGPD1-GAL4(848).ER::URA3/ura3::pGPD1-GAL4(848).ER::URA3 NDC10-6HA::HIS3MX6/NDC10-6HA::HIS3MX6 mcm21Δ::KANMX/mcm21Δ::KANMX | SK1 | 6g |
| AM10664 | MATa/alpha ndt80∆::LEU2/ndt80∆::LEU2 mcm21Δ::KANMX/mcm21Δ::KANMX | SK1 | 6g |
| AM10686 | MATa/alpha ndt80∆::LEU2/ndt80∆::LEU2 iml3Δ::KANMX/iml3Δ::KANMX | SK1 | 6g |
| AM11158 | MATa/alpha GAL-NDT80::TRP1/GAL-NDT80::TRP1 ura3::pGPD1-GAL4(848).ER::URA3/ura3::pGPD1-GAL4(848).ER::URA3 DSN1-6HIS-3FLAG::URA3/DSN1-6HIS-3FLAG::URA3 | SK1 | 1b,2c, s1a, s3b |
| AM11189 | MATa/alpha GAL-NDT80::TRP1/GAL-NDT80::TRP1 ura3::pGPD1-GAL4(848).ER::URA3/ura3::pGPD1-GAL4(848).ER::URA3 | SK1 | 6g |
| AM11633 | MATa/alpha ndt80∆::LEU2/ndt80∆::LEU2 | SK1 | 4d |
| AM11720 | MATa/alpha promURA3::TetR::GFP::LEU2/promURA3::TetR::GFP::LEU2 tetOx224-HIS3/tetOx224-HIS3 ctf19Δ::KanMX6/ctf19Δ::KanMX6 | SK1 | 3j |
| AM13406 | MATa/alpha cdc20::pCLB2-3HA-CDC20::KanMX6/cdc20::pCLB2-3HA-CDC20::KanMX6 his3::HIS3p-GFP-TUB1-HIS3/his3::HIS3p-GFP-TUB1-HIS3 MTW1-tdTomato::NAT/MTW1-tdTomato::NAT ndc80::pCLB2-NDC80::KanMX6/ndc80::pCLB2-NDC80::KanMX6 | SK1 | 5f, 5g |
| AM13689 | MATa/alpha cdc20::pCLB2-3HA-CDC20::KanMX6/cdc20::pCLB2-3HA-CDC20::KanMX6 DSN1-6HIS-3FLAG::URA3/DSN1-6HIS-3FLAG::URA3 mcm21∆::KANMX/mcm21∆::KANMX | SK1 | 5i, 5j |
| AM13691 | MATa/alpha GAL-NDT80::TRP1/GAL-NDT80::TRP1 ura3::pGPD1-GAL4(848).ER::URA3/ura3::pGPD1-GAL4(848).ER::URA3 MTW1-tdTomato::NAT/MTW1-tdTomato::NAT Ndc80-yEGFP::KanMX/Ndc80-yEGFP::KanMX | SK1 | s6b |
| AM13693 | MATa/alpha GAL-NDT80::TRP1/GAL-NDT80::TRP1 ura3::pGPD1-GAL4(848).ER::URA3/ura3::pGPD1-GAL4(848).ER::URA3 MTW1-tdTomato::NAT/MTW1-tdTomato::NAT | SK1 | 5a, 4f, 4h-k, s6b |
| AM13699 | MATa/alpha cdc20::pCLB2-3HA-CDC20::KanMX6/cdc20::pCLB2-3HA-CDC20::KanMX6 DSN1-6HIS-3FLAG::URA3/DSN1-6HIS-3FLAG::URA3 iml3∆::KANMX/iml3∆::KANMX | SK1 | 5i, 5j |
| AM13716 | MATa/alpha REC8-GFP-URA3/REC8-GFP-URA3 PDS1-tdTomato-KITRP1/PDS1-tdTomato-KITRP1 MTW1-tdTomato::NAT/MTW1-tdTomato::NAT | SK1 | 4g-h |
| AM13724 | MATa/alpha GAL-NDT80::TRP1/GAL-NDT80::TRP1 ura3::pGPD1-GAL4(848).ER::URA3/ura3::pGPD1-GAL4(848).ER::URA3 MTW1-tdTomato::NAT/MTW1-tdTomato::NAT Ndc80-yEGFP::KanMX/Ndc80-yEGFP::KanMX mcm21Δ::KANMX6/mcm21Δ::KANMX6 | SK1 | 5b, 4g-k, s6b |
| AM14099 | MATa/alpha cdc20::pCLB2-3HA-CDC20::KanMX6/cdc20::pCLB2-3HA-CDC20::KanMX6 his3::HIS3p-GFP-TUB1-HIS3/his3::HIS3p-GFP-TUB1-HIS3 MTW1-tdTomato::NAT/MTW1-tdTomato::NAT iml3∆::KanMX6/iml3∆::KanMX6 | SK1 | 5f, 5g |
| AM14230 | MATa/alpha GAL-NDT80::TRP1/GAL-NDT80::TRP1 ura3::pGPD1-GAL4(848).ER::URA3/ura3::pGPD1-GAL4(848).ER::URA3 MTW1-tdTomato::NAT/MTW1-tdTomato::NAT his3::HIS3p-GFP-TUB1-HIS3/his3::HIS3p-GFP-TUB1-HIS3 | SK1 | 5e |
| AM14746 | MATa/alpha cdc20::pCLB2-3HA-CDC20::KanMX6/cdc20::pCLB2-3HA-CDC20::KanMX6 CNN1-6HA::TRP/CNN1-6HA::TRP | SK1 | 6b |
| AM14752 | MATa/alpha cdc20::pCLB2-3HA-CDC20::KanMX6/cdc20::pCLB2-3HA-CDC20::KanMX6 CNN1-6HA::TRP/CNN1-6HA::TRP mcm21Δ::KANMX6/mcm21Δ::KANMX6 | SK1 | 6b |
| AM14754 | MATa/alpha cdc20::pCLB2-3HA-CDC20::KanMX6/cdc20::pCLB2-3HA-CDC20::KanMX6 CNN1-6HA::TRP/CNN1-6HA::TRP iml3Δ::KANMX6/iml3Δ::KANMX6 | SK1 | 6b |
| AM14986 | MATa/alpha promURA3::TetR::GFP::LEU2/promURA3::TetR::GFP::LEU2 tetOx224-HIS3/tetOx224-HIS3 cnn1Δ::KanMX6/cnn1Δ::KanMX6 | SK1 | 6c |
| AM15740 | MATa/alpha cdc20::pCLB2-3HA-CDC20::KanMX6/cdc20::pCLB2-3HA-CDC20::KanMX6 DSN1-6HIS-6HA:NAT/DSN1-6HIS-6HA:NAT | SK1 | 1b, s1A, s3b |
| AM16170 | MATa/alpha promURA3::TetR::GFP::LEU2 tetOx224-HIS3 | SK1 | 3i, 3k |
| AM16174 | MATa/alpha promURA3::TetR::GFP::LEU2/promURA3::TetR::GFP::LEU2 tetOx224-HIS3/tetOx224-HIS3 | SK1 | 3j |
| AM20054 | MATa/alpha GAL-NDT80::TRP1/GAL-NDT80::TRP1 ura3::pGPD1-GAL4(848).ER::URA3/ura3::pGPD1-GAL4(848).ER::URA3 MTW1-tdTomato::NAT/MTW1-tdTomato::NAT his3::HIS3p-GFP-TUB1-HIS3/his3::HIS3p-GFP-TUB1-HIS3 mcm21Δ::KANMX6/mcm21Δ::KANMX6 | SK1 | 5e |
| AM20078 | MATa/alpha ndt80∆::LEU2/ndt80∆::LEU2 DSN1-6HIS-3FLAG::URA3/DSN1-6HIS-3FLAG::URA3 | SK1 | 4d |
| AM20080 | MATa/alpha ndt80∆::LEU2/ndt80∆::LEU2 DSN1-6HIS-3FLAG::URA3/DSN1-6HIS-3FLAG::URA3 mcm21Δ::KANMX6/mcm21Δ::KANMX6 | SK1 | 4d |
| AM20107 | MATa/alpha GAL-NDT80::TRP1/GAL-NDT80::TRP1 ura3::pGPD1-GAL4(848).ER::URA3/ura3::pGPD1-GAL4(848).ER::URA3 MTW1-tdTomato::NAT/MTW1-tdTomato::NAT his3::HIS3p-GFP-TUB1-HIS3/his3::HIS3p-GFP-TUB1-HIS3 iml3Δ::KANMX6/iml3Δ::KANMX6 | SK1 | 5e |
| AM20705 | MATalpha ura3::LacI-3FLAG::URA3 ndt80D::LEU2 <CEN3-TALO8>:TRP | SK1 | 1c, 2a, 2d, s3a |
| AM20707 | MATalpha ura3::LacI-3FLAG::URA3 ndt80D::LEU2 <CEN3*-TALO8>:TRP | SK1 | 1c, 2a, 2d, s3a |
| AM20732 | MATa/alpha promURA3::TetR::GFP::LEU2/promURA3::TetR::GFP::LEU2 tetOx224-HIS3/tetOx224-HIS3 HTB1-mCherry-HIS3MX6/HTB1-mCherry-HIS3MX6 mcm21Δ::KANMX6/mcm21Δ::KANMX6 | SK1 | 3c, 3d, 3e |
| AM20734 | MATa/alpha promURA3::TetR::GFP::LEU2/promURA3::TetR::GFP::LEU2 tetOx224-HIS3/tetOx224-HIS3 HTB1-mCherry-HIS3MX6/HTB1-mCherry-HIS3MX6 | SK1 | 3c, 3d, 3e |
| AM20751 | MATa/alpha ura3::LacI-3FLAG::URA3 ndt80D::LEU2/ndt80D::LEU2 iml3Δ::KANMX6/iml3Δ::KANMX6 <CEN3-TALO8>:TRP | SK1 | 4a |
| AM20755 | MATa/alpha ura3::LacI-3FLAG::URA3 ndt80D::LEU2/ndt80D::LEU2 mcm21Δ::KANMX6/mcm21Δ::KANMX6 <CEN3-TALO8>:TRP | SK1 | 4a |
| AM20782 | MATa/alpha ura3::LacI-3FLAG::URA3 ndt80D::LEU2/ndt80D::LEU2 <CEN3*-TALO8>:TRP | SK1 | 1c, 2a, 2b, 2d, 4a, s2a, , s3a |
| AM20785 | MATa/alpha ura3::LacI-3FLAG::URA3 ndt80D::LEU2/ndt80D::LEU2 <CEN3-TALO8>:TRP | SK1 | 1c, 2a, 2b, 2d, 4a, s3a |
| AM22604 | MATa/alpha GAL-NDT80::TRP1/GAL-NDT80::TRP1 ura3::pGPD1-GAL4(848).ER::URA3/ura3::pGPD1-GAL4(848).ER::URA3 cdc20::pCLB2-3HA-CDC20::KanMX6/cdc20::pCLB2-3HA-CDC20::KanMX6 | SK1 | 1b |
| AM23739 | MATa/alpha GAL-NDT80::TRP1/GAL-NDT80::TRP1 ura3::pGPD1-GAL4(848).ER::URA3/ura3::pGPD1-GAL4(848).ER::URA3 MTW1-tdTomato::NAT/MTW1-tdTomato::NAT Ame1-6HA:TRP1/Ame1-6HA:TRP1 mcm21Δ::KANMX6/mcm21Δ::KANMX6 | SK1 | 6a |
| AM23740 | MATa/alpha GAL-NDT80::TRP1/GAL-NDT80::TRP1 ura3::pGPD1-GAL4(848).ER::URA3/ura3::pGPD1-GAL4(848).ER::URA3 MTW1-tdTomato::NAT/MTW1-tdTomato::NAT Ame1-6HA:TRP1/Ame1-6HA:TRP1 iml3Δ::KANMX6/iml3Δ::KANMX6 | SK1 | 6a |
| AM23741 | MATa/alpha GAL-NDT80::TRP1/GAL-NDT80::TRP1 ura3::pGPD1-GAL4(848).ER::URA3/ura3::pGPD1-GAL4(848).ER::URA3 MTW1-tdTomato::NAT/MTW1-tdTomato::NAT Ame1-6HA:TRP1/Ame1-6HA:TRP1 mcm21Δ::KANMX6/mcm21Δ::KANMX6 | SK1 | 6a |
| AM23799 | MATa/alpha GAL-NDT80::TRP1/GAL-NDT80::TRP1 ura3::pGPD1-GAL4(848).ER::URA3/ura3::pGPD1-GAL4(848).ER::URA3 MTW1-tdTomato::NAT/MTW1-tdTomato::NAT | SK1 | 6a, 7f |
| AM24060 | MATa/alpha Ndc80-yEGFP::KanMX/Ndc80-yEGFP::KanMX DSN1-tdTomato::NAT/DSN1-tdTomato::NAT mcm21∆::hphMX/mcm21∆::hphMX | SK1 | 5d |
| AM24061 | MATa/alpha Ndc80(∆2-28)-yEGFP::KanMX/Ndc80(∆2-28)-yEGFP::KanMX DSN1-tdTomato::NAT/DSN1-tdTomato::NAT mcm21∆::hphMX/mcm21∆::hphMX | SK1 | 5d |
| AM24132 | MATa/alpha Ndc80(∆2-28)-yEGFP::KanMX/Ndc80(∆2-28)-yEGFP::KanMX DSN1-tdTomato::NAT/DSN1-tdTomato::NAT | SK1 | 5d |
| AM24133 | MATa/alpha Ndc80-yEGFP::KanMX/Ndc80-yEGFP::KanMX DSN1-tdTomato::NAT/DSN1-tdTomato::NAT | SK1 | 5d |
| AM24317 | MATa/alpha ura3::LacI-3FLAG::URA3 cdc20::pCLB2-3HA-CDC20::KanMX6/cdc20::pCLB2-3HA-CDC20::KanMX6 <CEN3-TALO8>:TRP | SK1 | 1c, 2a, 2b, 2d, 4a, , s3a |
| AM24317 | MATa/alpha cdc20::pCLB2-3HA-CDC20::KanMX6/cdc20::pCLB2-3HA-CDC20::KanMX6 <CEN3-TALO8>:TRP | SK1 | 2b |
| AM24318 | MATa/alpha ura3::LacI-3FLAG::URA3 cdc20::pCLB2-3HA-CDC20::KanMX6/cdc20::pCLB2-3HA-CDC20::KanMX6 <CEN3*-TALO8>:TRP | SK1 | 1c, 2a, 2b, 2d, 4a, , s3a |
| AM24319 | MATa/alpha ura3::LacI-3FLAG::URA3 cdc20::pCLB2-3HA-CDC20::KanMX6/cdc20::pCLB2-3HA-CDC20::KanMX6 iml3Δ::KANMX6/iml3Δ::KANMX6 <CEN3-TALO8>:TRP | SK1 | 4a |
| AM24320 | MATa/alpha ura3::LacI-3FLAG::URA3 cdc20::pCLB2-3HA-CDC20::KanMX6/cdc20::pCLB2-3HA-CDC20::KanMX6 mcm21Δ::KANMX6/mcm21Δ::KANMX6 <CEN3-TALO8>:TRP | SK1 | 4a |
| AM25605 | MATa/alpha mcm21Δ::KANMX6/mcm21Δ::KANMX6 | SK1 | 3a, 3b |
| AM25606 | MATa/alpha chl4Δ::KANMX6/chl4Δ::KANMX6 | SK1 | 3a, 3b |
| AM25607 | MATa/alpha iml3Δ::KANMX6/iml3Δ::KANMX6 | SK1 | 3a, 3b |
| AM26610 | MATa/alpha promURA3::TetR::GFP::LEU2/promURA3::TetR::GFP::LEU2 tetOx224-HIS3/tetOx224-HIS3 ctf19-9A::LEU2/ctf19-9A::LEU2 | SK1 | 3j |
| AM26641 | MATa/alpha promURA3::TetR::GFP::LEU2/ tetOx224-HIS3/ ctf19Δ::KANMX6/ctf19Δ::KANMX6 | SK1 | 3i, 3k |
| AM26642 | MATa/alpha promURA3::TetR::GFP::LEU2/ tetOx224-HIS3/ ctf19-9A::LEU2/ctf19-9A::LEU2 | SK1 | 3i, 3k |
| AM26937 | MATa/alpha GAL-NDT80::TRP1/GAL-NDT80::TRP1 ura3::pGPD1-GAL4(848).ER::URA3/ura3::pGPD1-GAL4(848).ER::URA3 MIF2-mNeonGreen:LEU/MIF2-mNeonGreen:LEU mcm21∆::hphMX/mcm21∆::hphMX | SK1 | 6d |
| AM26938 | MATa/alpha GAL-NDT80::TRP1/GAL-NDT80::TRP1 ura3::pGPD1-GAL4(848).ER::URA3/ura3::pGPD1-GAL4(848).ER::URA3 MIF2-mNeonGreen:LEU/MIF2-mNeonGreen:LEU | SK1 | 6d |
| AM26939 | MATa/alpha GAL-NDT80::TRP1/GAL-NDT80::TRP1 ura3::pGPD1-GAL4(848).ER::URA3/ura3::pGPD1-GAL4(848).ER::URA3 MTW1-tdTomato::NAT/MTW1-tdTomato::NAT NDC10-mNeonGreen:LEU2/NDC10-mNeonGreen:LEU2 | SK1 | 4b, 6f, 7h |
| AM26940 | MATa/alpha GAL-NDT80::TRP1/GAL-NDT80::TRP1 ura3::pGPD1-GAL4(848).ER::URA3/ura3::pGPD1-GAL4(848).ER::URA3 MTW1-tdTomato::NAT/MTW1-tdTomato::NAT NDC10-mNeonGreen:LEU2/NDC10-mNeonGreen:LEU2 mcm21∆::hphMX/mcm21∆::hphMX | SK1 | 4b, 6f, 7h |
| AM26943 | MATa/alpha GAL-NDT80::TRP1/GAL-NDT80::TRP1 ura3::pGPD1-GAL4(848).ER::URA3/ura3::pGPD1-GAL4(848).ER::URA3 MTW1-tdTomato::NAT/MTW1-tdTomato::NAT NDC10-mNeonGreen:LEU2/NDC10-mNeonGreen:LEU2 dsn1::dsn1(S240D/S250D)-6HIS-3FLAG::URA3/dsn1::dsn1(S240D/S250D)-6HIS-3FLAG::URA3 mcm21∆::hphMX/mcm21∆::hphMX | SK1 | 7h |
| AM26970 | MATa/alpha REC8-GFP-URA3/REC8-GFP-URA3 PDS1-tdTomato-KITRP1/PDS1-tdTomato-KITRP1 MTW1-tdTomato::NAT/MTW1-tdTomato::NAT ctf19-9A::LEU2/ctf19-9A::LEU2 | SK1 | 4g-h |
| AM27017 | MATa/alpha GAL-NDT80::TRP1/GAL-NDT80::TRP1 ura3::pGPD1-GAL4(848).ER::URA3/ura3::pGPD1-GAL4(848).ER::URA3 MTW1-tdTomato::NAT/MTW1-tdTomato::NAT NDC10-mNeonGreen:LEU2/NDC10-mNeonGreen:LEU2 ame1::pCLB2-3HA-AME1::KanMX6/ame1::pCLB2-3HA-AME1::KanMX6 | SK1 | 4b |
| AM27333 | MATa/alpha GAL-NDT80::TRP1/GAL-NDT80::TRP1 ura3::pGPD1-GAL4(848).ER::URA3/ura3::pGPD1-GAL4(848).ER::URA3 MTW1-tdTomato::NAT/MTW1-tdTomato::NAT NDC10-mNeonGreen:LEU2/NDC10-mNeonGreen:LEU2 clb3::KanMX6::pCUP1-CLB3/clb3::KanMX6::pCUP1-CLB3 mcm21Δ::KANMX6/mcm21Δ::KANMX6 | SK1 | 4c |
| AM27334 | MATa/alpha GAL-NDT80::TRP1/GAL-NDT80::TRP1 ura3::pGPD1-GAL4(848).ER::URA3/ura3::pGPD1-GAL4(848).ER::URA3 MTW1-tdTomato::NAT/MTW1-tdTomato::NAT NDC10-mNeonGreen:LEU2/NDC10-mNeonGreen:LEU2 clb3::KanMX6::pCUP1-CLB3/clb3::KanMX6::pCUP1-CLB3 | SK1 | 4c |
| AM27370 | MATa/alpha GAL-NDT80::TRP1/GAL-NDT80::TRP1 ura3::pGPD1-GAL4(848).ER::URA3/ura3::pGPD1-GAL4(848).ER::URA3 MTW1-tdTomato::NAT/MTW1-tdTomato::NAT NDC10-mNeonGreen:LEU2/NDC10-mNeonGreen:LEU2 dsn1::dsn1(S240D/S250D)-6HIS-3FLAG::URA3/dsn1::dsn1(S240D/S250D)-6HIS-3FLAG::URA3 | SK1 | 7h |
| AM27371 | MATa/alpha GAL-NDT80::TRP1/GAL-NDT80::TRP1 ura3::pGPD1-GAL4(848).ER::URA3/ura3::pGPD1-GAL4(848).ER::URA3 MTW1-tdTomato::NAT/MTW1-tdTomato::NAT NDC10-mNeonGreen:LEU2/NDC10-mNeonGreen:LEU2 okp1::pCLB2-3HA-OKP1::KanMX6/okp1::pCLB2-3HA-OKP1::KanMX6 | SK1 | 4b |
| AM27723 | MATa/alpha GAL-NDT80::TRP1/GAL-NDT80::TRP1 ura3::pGPD1-GAL4(848).ER::URA3/ura3::pGPD1-GAL4(848).ER::URA3 CSE4-mNeonGreen/CSE4-mNeonGreen | SK1 | 6e |
| AM27724 | MATa/alpha GAL-NDT80::TRP1/GAL-NDT80::TRP1 ura3::pGPD1-GAL4(848).ER::URA3/ura3::pGPD1-GAL4(848).ER::URA3 CSE4-mNeonGreen/CSE4-mNeonGreen mcm21∆::hphM/mcm21∆::hphM | SK1 | 6e |
| AM28330 | MATa/alpha irt1::NAT::pCUP-IME1/irt1::NAT::pCUP-IME1 pIME4::NAT::pCUP-IME4/pIME4::NAT::pCUP-IME4 AME1-mNeonGreen:KILEU2/AME1-mNeonGreen:KILEU2 mcm21∆::hphM/mcm21∆::hphM | SK1 | 6h-i |
| AM28331 | MATa/alpha irt1::NAT::pCUP-IME1/irt1::NAT::pCUP-IME1 pIME4::NAT::pCUP-IME4/pIME4::NAT::pCUP-IME4 AME1-mNeonGreen:KILEU2/AME1-mNeonGreen:KILEU2 | SK1 | 6h-i |
| AM28767 | MATa ura3::pADH1-OsTIR1-9MYC::URA3 MTW1-tdTomato::NAT Bir1-AID::kanMX mcm21∆::KanMX6 | w303 | 7b-e, s6c |
| AM28769 | MATa ura3::pADH1-OsTIR1-9MYC::URA3 MTW1-tdTomato::NAT Bir1-AID::kanMX | w303 | 7b-e, s6c |
| AM29121 | MATa/alpha GAL-NDT80::TRP1/GAL-NDT80::TRP1 ura3::pGPD1-GAL4(848).ER::URA3/ura3::pGPD1-GAL4(848).ER::URA3 MTW1-tdTomato::NAT/MTW1-tdTomato::NAT ipl1::KanMX6::pCLB2-3HA-IPL1/ipl1::KanMX6::pCLB2-3HA-IPL1 | SK1 | 7f |
| AM1603 | MATa/alpha /promURA3::TetR::GFP::LEU2 tetOx224-URA3 | SK1 | 6c |
| AM1835 | MATa/alpha | SK1 | 3a, 3b |
| AM3560 | MATa/alpha cdc20::pCLB2-3HA-CDC20::KanMX6/cdc20::pCLB2-3HA-CDC20::KanMX6 | SK1 | 6b |
| AM3722 | MATa/alpha ctf19Δ::KanMX6/ctf19Δ::KanMX6 | SK1 | 3a, 3b |
| AM4014 | MATa/alpha ndt80∆::LEU2/ndt80∆::LEU2 | SK1 | 6g |
| AM6142 | MATa/alpha ndt80::pGAL-NDT80::TRP1/ndt80::pGAL-NDT80::TRP1 ura3::pGPD1-GAL4(848).ER::URA3/ura3::pGPD1-GAL4(848).ER::URA3 ubr1Δ::KanMX4/ubr1Δ::KanMX4 REC8-3HA::URA3/ REC8-3HA::URA3 PDS1-18MYC::LEU2/PDS1-18MYC::LEU2 | SK1 | s5b, s5c |
| AM8291 | MATa/alpha cdc20::pCLB2-3HA-CDC20::KanMX6/cdc20::pCLB2-3HA-CDC20::KanMX6 DSN1-6HIS-3FLAG::URA3/DSN1-6HIS-3FLAG::URA3 | SK1 | 1b,2c, s1a, s1b, s3b |
| AM8292 | MATa/alpha cdc20::pCLB2-3HA-CDC20::KanMX6/cdc20::pCLB2-3HA-CDC20::KanMX6 DSN1-6HIS-3FLAG::URA3/DSN1-6HIS-3FLAG::URA3 | SK1 | 5i, 5j |
| AM8382 | MATa/alpha ndt80::pGAL-NDT80::TRP1/ndt80::pGAL-NDT80::TRP1 ura3::pGPD1-GAL4(848).ER::URA3/ura3::pGPD1-GAL4(848).ER::URA3 ubr1Δ::KanMX4/ubr1Δ::KanMX4 REC8-3HA::URA3/ REC8-3HA::URA3 PDS1-18MYC::LEU2/PDS1-18MYC::LEU2 iml3Δ::KanMX6/iml3Δ::KanMX6 | SK1 | s5b, s5c |
| AM8769 | MATa/alpha GAL-NDT80::TRP1/GAL-NDT80::TRP1 ura3::pGPD1-GAL4(848).ER::URA3/ura3::pGPD1-GAL4(848).ER::URA3 NDC10-6HA::HIS3MX6/NDC10-6HA::HIS3MX6 | SK1 | 6g |
| AM8805 | MATa/alpha ndt80::pGAL-NDT80::TRP1/ndt80::pGAL-NDT80::TRP1 ura3::pGPD1-GAL4(848).ER::URA3/ura3::pGPD1-GAL4(848).ER::URA3 ubr1Δ::KanMX4/ubr1Δ::KanMX4 REC8-3HA::URA3/ REC8-3HA::URA3 PDS1-18MYC::LEU2/PDS1-18MYC::LEU2 mcm21Δ::KanMX6/mcm21Δ::KanMX6 | SK1 | s5b, s5c |
| AM8825 | MATa/alpha cdc20::pCLB2-3HA-CDC20::KanMX6/cdc20::pCLB2-3HA-CDC20::KanMX6 his3::HIS3p-GFP-TUB1-HIS3/his3::HIS3p-GFP-TUB1-HIS3 MTW1-tdTomato::NAT/MTW1-tdTomato::NAT | SK1 | 5f, 5g |
| AM8861 | MATa/alpha GAL-NDT80::TRP1/GAL-NDT80::TRP1 ura3::pGPD1-GAL4(848).ER::URA3/ura3::pGPD1-GAL4(848).ER::URA3 NDC10-6HA::HIS3MX6/NDC10-6HA::HIS3MX6 mcm21Δ::KANMX/mcm21Δ::KANMX | SK1 | 6g |
| AM10664 | MATa/alpha ndt80∆::LEU2/ndt80∆::LEU2 mcm21Δ::KANMX/mcm21Δ::KANMX | SK1 | 6g |
| AM10686 | MATa/alpha ndt80∆::LEU2/ndt80∆::LEU2 iml3Δ::KANMX/iml3Δ::KANMX | SK1 | 6g |
| AM11158 | MATa/alpha GAL-NDT80::TRP1/GAL-NDT80::TRP1 ura3::pGPD1-GAL4(848).ER::URA3/ura3::pGPD1-GAL4(848).ER::URA3 DSN1-6HIS-3FLAG::URA3/DSN1-6HIS-3FLAG::URA3 | SK1 | 1b,2c, s1a, s3b |
| AM11189 | MATa/alpha GAL-NDT80::TRP1/GAL-NDT80::TRP1 ura3::pGPD1-GAL4(848).ER::URA3/ura3::pGPD1-GAL4(848).ER::URA3 | SK1 | 6g |
| AM11633 | MATa/alpha ndt80∆::LEU2/ndt80∆::LEU2 | SK1 | 4d |
| AM11720 | MATa/alpha promURA3::TetR::GFP::LEU2/promURA3::TetR::GFP::LEU2 tetOx224-HIS3/tetOx224-HIS3 ctf19Δ::KanMX6/ctf19Δ::KanMX6 | SK1 | 3j |
| AM13406 | MATa/alpha cdc20::pCLB2-3HA-CDC20::KanMX6/cdc20::pCLB2-3HA-CDC20::KanMX6 his3::HIS3p-GFP-TUB1-HIS3/his3::HIS3p-GFP-TUB1-HIS3 MTW1-tdTomato::NAT/MTW1-tdTomato::NAT ndc80::pCLB2-NDC80::KanMX6/ndc80::pCLB2-NDC80::KanMX6 | SK1 | 5f, 5g |
| AM13689 | MATa/alpha cdc20::pCLB2-3HA-CDC20::KanMX6/cdc20::pCLB2-3HA-CDC20::KanMX6 DSN1-6HIS-3FLAG::URA3/DSN1-6HIS-3FLAG::URA3 mcm21∆::KANMX/mcm21∆::KANMX | SK1 | 5i, 5j |
| AM13691 | MATa/alpha GAL-NDT80::TRP1/GAL-NDT80::TRP1 ura3::pGPD1-GAL4(848).ER::URA3/ura3::pGPD1-GAL4(848).ER::URA3 MTW1-tdTomato::NAT/MTW1-tdTomato::NAT Ndc80-yEGFP::KanMX/Ndc80-yEGFP::KanMX | SK1 | s6b |
| AM13693 | MATa/alpha GAL-NDT80::TRP1/GAL-NDT80::TRP1 ura3::pGPD1-GAL4(848).ER::URA3/ura3::pGPD1-GAL4(848).ER::URA3 MTW1-tdTomato::NAT/MTW1-tdTomato::NAT | SK1 | 5a, 4f, 4h-k, s6b |
| AM13699 | MATa/alpha cdc20::pCLB2-3HA-CDC20::KanMX6/cdc20::pCLB2-3HA-CDC20::KanMX6 DSN1-6HIS-3FLAG::URA3/DSN1-6HIS-3FLAG::URA3 iml3∆::KANMX/iml3∆::KANMX | SK1 | 5i, 5j |
| AM13716 | MATa/alpha REC8-GFP-URA3/REC8-GFP-URA3 PDS1-tdTomato-KITRP1/PDS1-tdTomato-KITRP1 MTW1-tdTomato::NAT/MTW1-tdTomato::NAT | SK1 | 4g-h |
| AM13724 | MATa/alpha GAL-NDT80::TRP1/GAL-NDT80::TRP1 ura3::pGPD1-GAL4(848).ER::URA3/ura3::pGPD1-GAL4(848).ER::URA3 MTW1-tdTomato::NAT/MTW1-tdTomato::NAT Ndc80-yEGFP::KanMX/Ndc80-yEGFP::KanMX mcm21Δ::KANMX6/mcm21Δ::KANMX6 | SK1 | 5b, 4g-k, s6b |
| AM14099 | MATa/alpha cdc20::pCLB2-3HA-CDC20::KanMX6/cdc20::pCLB2-3HA-CDC20::KanMX6 his3::HIS3p-GFP-TUB1-HIS3/his3::HIS3p-GFP-TUB1-HIS3 MTW1-tdTomato::NAT/MTW1-tdTomato::NAT iml3∆::KanMX6/iml3∆::KanMX6 | SK1 | 5f, 5g |
| AM14230 | MATa/alpha GAL-NDT80::TRP1/GAL-NDT80::TRP1 ura3::pGPD1-GAL4(848).ER::URA3/ura3::pGPD1-GAL4(848).ER::URA3 MTW1-tdTomato::NAT/MTW1-tdTomato::NAT his3::HIS3p-GFP-TUB1-HIS3/his3::HIS3p-GFP-TUB1-HIS3 | SK1 | 5e |
| AM14746 | MATa/alpha cdc20::pCLB2-3HA-CDC20::KanMX6/cdc20::pCLB2-3HA-CDC20::KanMX6 CNN1-6HA::TRP/CNN1-6HA::TRP | SK1 | 6b |
| AM14752 | MATa/alpha cdc20::pCLB2-3HA-CDC20::KanMX6/cdc20::pCLB2-3HA-CDC20::KanMX6 CNN1-6HA::TRP/CNN1-6HA::TRP mcm21Δ::KANMX6/mcm21Δ::KANMX6 | SK1 | 6b |
| AM14754 | MATa/alpha cdc20::pCLB2-3HA-CDC20::KanMX6/cdc20::pCLB2-3HA-CDC20::KanMX6 CNN1-6HA::TRP/CNN1-6HA::TRP iml3Δ::KANMX6/iml3Δ::KANMX6 | SK1 | 6b |
| AM14986 | MATa/alpha promURA3::TetR::GFP::LEU2/promURA3::TetR::GFP::LEU2 tetOx224-HIS3/tetOx224-HIS3 cnn1Δ::KanMX6/cnn1Δ::KanMX6 | SK1 | 6c |
| AM15740 | MATa/alpha cdc20::pCLB2-3HA-CDC20::KanMX6/cdc20::pCLB2-3HA-CDC20::KanMX6 DSN1-6HIS-6HA:NAT/DSN1-6HIS-6HA:NAT | SK1 | 1b, s1A, s3b |
| AM16170 | MATa/alpha promURA3::TetR::GFP::LEU2 tetOx224-HIS3 | SK1 | 3i, 3k |
| AM16174 | MATa/alpha promURA3::TetR::GFP::LEU2/promURA3::TetR::GFP::LEU2 tetOx224-HIS3/tetOx224-HIS3 | SK1 | 3j |
| AM20054 | MATa/alpha GAL-NDT80::TRP1/GAL-NDT80::TRP1 ura3::pGPD1-GAL4(848).ER::URA3/ura3::pGPD1-GAL4(848).ER::URA3 MTW1-tdTomato::NAT/MTW1-tdTomato::NAT his3::HIS3p-GFP-TUB1-HIS3/his3::HIS3p-GFP-TUB1-HIS3 mcm21Δ::KANMX6/mcm21Δ::KANMX6 | SK1 | 5e |
| AM20078 | MATa/alpha ndt80∆::LEU2/ndt80∆::LEU2 DSN1-6HIS-3FLAG::URA3/DSN1-6HIS-3FLAG::URA3 | SK1 | 4d |
| AM20080 | MATa/alpha ndt80∆::LEU2/ndt80∆::LEU2 DSN1-6HIS-3FLAG::URA3/DSN1-6HIS-3FLAG::URA3 mcm21Δ::KANMX6/mcm21Δ::KANMX6 | SK1 | 4d |
| AM20107 | MATa/alpha GAL-NDT80::TRP1/GAL-NDT80::TRP1 ura3::pGPD1-GAL4(848).ER::URA3/ura3::pGPD1-GAL4(848).ER::URA3 MTW1-tdTomato::NAT/MTW1-tdTomato::NAT his3::HIS3p-GFP-TUB1-HIS3/his3::HIS3p-GFP-TUB1-HIS3 iml3Δ::KANMX6/iml3Δ::KANMX6 | SK1 | 5e |
| AM20705 | MATalpha ura3::LacI-3FLAG::URA3 ndt80D::LEU2 <CEN3-TALO8>:TRP | SK1 | 1c, 2a, 2d, s3a |
| AM20707 | MATalpha ura3::LacI-3FLAG::URA3 ndt80D::LEU2 <CEN3*-TALO8>:TRP | SK1 | 1c, 2a, 2d, s3a |
| AM20732 | MATa/alpha promURA3::TetR::GFP::LEU2/promURA3::TetR::GFP::LEU2 tetOx224-HIS3/tetOx224-HIS3 HTB1-mCherry-HIS3MX6/HTB1-mCherry-HIS3MX6 mcm21Δ::KANMX6/mcm21Δ::KANMX6 | SK1 | 3c, 3d, 3e |
| AM20734 | MATa/alpha promURA3::TetR::GFP::LEU2/promURA3::TetR::GFP::LEU2 tetOx224-HIS3/tetOx224-HIS3 HTB1-mCherry-HIS3MX6/HTB1-mCherry-HIS3MX6 | SK1 | 3c, 3d, 3e |
| AM20751 | MATa/alpha ura3::LacI-3FLAG::URA3 ndt80D::LEU2/ndt80D::LEU2 iml3Δ::KANMX6/iml3Δ::KANMX6 <CEN3-TALO8>:TRP | SK1 | 4a |
| AM20755 | MATa/alpha ura3::LacI-3FLAG::URA3 ndt80D::LEU2/ndt80D::LEU2 mcm21Δ::KANMX6/mcm21Δ::KANMX6 <CEN3-TALO8>:TRP | SK1 | 4a |
| AM20782 | MATa/alpha ura3::LacI-3FLAG::URA3 ndt80D::LEU2/ndt80D::LEU2 <CEN3*-TALO8>:TRP | SK1 | 1c, 2a, 2b, 2d, 4a, s2a, , s3a |
| AM20785 | MATa/alpha ura3::LacI-3FLAG::URA3 ndt80D::LEU2/ndt80D::LEU2 <CEN3-TALO8>:TRP | SK1 | 1c, 2a, 2b, 2d, 4a, s3a |
| AM22604 | MATa/alpha GAL-NDT80::TRP1/GAL-NDT80::TRP1 ura3::pGPD1-GAL4(848).ER::URA3/ura3::pGPD1-GAL4(848).ER::URA3 cdc20::pCLB2-3HA-CDC20::KanMX6/cdc20::pCLB2-3HA-CDC20::KanMX6 | SK1 | 1b |
| AM23739 | MATa/alpha GAL-NDT80::TRP1/GAL-NDT80::TRP1 ura3::pGPD1-GAL4(848).ER::URA3/ura3::pGPD1-GAL4(848).ER::URA3 MTW1-tdTomato::NAT/MTW1-tdTomato::NAT Ame1-6HA:TRP1/Ame1-6HA:TRP1 mcm21Δ::KANMX6/mcm21Δ::KANMX6 | SK1 | 6a |
| AM23740 | MATa/alpha GAL-NDT80::TRP1/GAL-NDT80::TRP1 ura3::pGPD1-GAL4(848).ER::URA3/ura3::pGPD1-GAL4(848).ER::URA3 MTW1-tdTomato::NAT/MTW1-tdTomato::NAT Ame1-6HA:TRP1/Ame1-6HA:TRP1 iml3Δ::KANMX6/iml3Δ::KANMX6 | SK1 | 6a |
| AM23741 | MATa/alpha GAL-NDT80::TRP1/GAL-NDT80::TRP1 ura3::pGPD1-GAL4(848).ER::URA3/ura3::pGPD1-GAL4(848).ER::URA3 MTW1-tdTomato::NAT/MTW1-tdTomato::NAT Ame1-6HA:TRP1/Ame1-6HA:TRP1 mcm21Δ::KANMX6/mcm21Δ::KANMX6 | SK1 | 6a |
| AM23799 | MATa/alpha GAL-NDT80::TRP1/GAL-NDT80::TRP1 ura3::pGPD1-GAL4(848).ER::URA3/ura3::pGPD1-GAL4(848).ER::URA3 MTW1-tdTomato::NAT/MTW1-tdTomato::NAT | SK1 | 6a, 7f |
| AM24060 | MATa/alpha Ndc80-yEGFP::KanMX/Ndc80-yEGFP::KanMX DSN1-tdTomato::NAT/DSN1-tdTomato::NAT mcm21∆::hphMX/mcm21∆::hphMX | SK1 | 5d |
| AM24061 | MATa/alpha Ndc80(∆2-28)-yEGFP::KanMX/Ndc80(∆2-28)-yEGFP::KanMX DSN1-tdTomato::NAT/DSN1-tdTomato::NAT mcm21∆::hphMX/mcm21∆::hphMX | SK1 | 5d |
| AM24132 | MATa/alpha Ndc80(∆2-28)-yEGFP::KanMX/Ndc80(∆2-28)-yEGFP::KanMX DSN1-tdTomato::NAT/DSN1-tdTomato::NAT | SK1 | 5d |
| AM24133 | MATa/alpha Ndc80-yEGFP::KanMX/Ndc80-yEGFP::KanMX DSN1-tdTomato::NAT/DSN1-tdTomato::NAT | SK1 | 5d |
| AM24317 | MATa/alpha ura3::LacI-3FLAG::URA3 cdc20::pCLB2-3HA-CDC20::KanMX6/cdc20::pCLB2-3HA-CDC20::KanMX6 <CEN3-TALO8>:TRP | SK1 | 1c, 2a, 2b, 2d, 4a, , s3a |
| AM24317 | MATa/alpha cdc20::pCLB2-3HA-CDC20::KanMX6/cdc20::pCLB2-3HA-CDC20::KanMX6 <CEN3-TALO8>:TRP | SK1 | 2b |
| AM24318 | MATa/alpha ura3::LacI-3FLAG::URA3 cdc20::pCLB2-3HA-CDC20::KanMX6/cdc20::pCLB2-3HA-CDC20::KanMX6 <CEN3*-TALO8>:TRP | SK1 | 1c, 2a, 2b, 2d, 4a, , s3a |
| AM24319 | MATa/alpha ura3::LacI-3FLAG::URA3 cdc20::pCLB2-3HA-CDC20::KanMX6/cdc20::pCLB2-3HA-CDC20::KanMX6 iml3Δ::KANMX6/iml3Δ::KANMX6 <CEN3-TALO8>:TRP | SK1 | 4a |
| AM24320 | MATa/alpha ura3::LacI-3FLAG::URA3 cdc20::pCLB2-3HA-CDC20::KanMX6/cdc20::pCLB2-3HA-CDC20::KanMX6 mcm21Δ::KANMX6/mcm21Δ::KANMX6 <CEN3-TALO8>:TRP | SK1 | 4a |
| AM25605 | MATa/alpha mcm21Δ::KANMX6/mcm21Δ::KANMX6 | SK1 | 3a, 3b |
| AM25606 | MATa/alpha chl4Δ::KANMX6/chl4Δ::KANMX6 | SK1 | 3a, 3b |
| AM25607 | MATa/alpha iml3Δ::KANMX6/iml3Δ::KANMX6 | SK1 | 3a, 3b |
| AM26610 | MATa/alpha promURA3::TetR::GFP::LEU2/promURA3::TetR::GFP::LEU2 tetOx224-HIS3/tetOx224-HIS3 ctf19-9A::LEU2/ctf19-9A::LEU2 | SK1 | 3j |
| AM26641 | MATa/alpha promURA3::TetR::GFP::LEU2/ tetOx224-HIS3/ ctf19Δ::KANMX6/ctf19Δ::KANMX6 | SK1 | 3i, 3k |
| AM26642 | MATa/alpha promURA3::TetR::GFP::LEU2/ tetOx224-HIS3/ ctf19-9A::LEU2/ctf19-9A::LEU2 | SK1 | 3i, 3k |
| AM26937 | MATa/alpha GAL-NDT80::TRP1/GAL-NDT80::TRP1 ura3::pGPD1-GAL4(848).ER::URA3/ura3::pGPD1-GAL4(848).ER::URA3 MIF2-mNeonGreen:LEU/MIF2-mNeonGreen:LEU mcm21∆::hphMX/mcm21∆::hphMX | SK1 | 6d |
| AM26938 | MATa/alpha GAL-NDT80::TRP1/GAL-NDT80::TRP1 ura3::pGPD1-GAL4(848).ER::URA3/ura3::pGPD1-GAL4(848).ER::URA3 MIF2-mNeonGreen:LEU/MIF2-mNeonGreen:LEU | SK1 | 6d |
| AM26939 | MATa/alpha GAL-NDT80::TRP1/GAL-NDT80::TRP1 ura3::pGPD1-GAL4(848).ER::URA3/ura3::pGPD1-GAL4(848).ER::URA3 MTW1-tdTomato::NAT/MTW1-tdTomato::NAT NDC10-mNeonGreen:LEU2/NDC10-mNeonGreen:LEU2 | SK1 | 4b, 6f, 7h |
| AM26940 | MATa/alpha GAL-NDT80::TRP1/GAL-NDT80::TRP1 ura3::pGPD1-GAL4(848).ER::URA3/ura3::pGPD1-GAL4(848).ER::URA3 MTW1-tdTomato::NAT/MTW1-tdTomato::NAT NDC10-mNeonGreen:LEU2/NDC10-mNeonGreen:LEU2 mcm21∆::hphMX/mcm21∆::hphMX | SK1 | 4b, 6f, 7h |
| AM26943 | MATa/alpha GAL-NDT80::TRP1/GAL-NDT80::TRP1 ura3::pGPD1-GAL4(848).ER::URA3/ura3::pGPD1-GAL4(848).ER::URA3 MTW1-tdTomato::NAT/MTW1-tdTomato::NAT NDC10-mNeonGreen:LEU2/NDC10-mNeonGreen:LEU2 dsn1::dsn1(S240D/S250D)-6HIS-3FLAG::URA3/dsn1::dsn1(S240D/S250D)-6HIS-3FLAG::URA3 mcm21∆::hphMX/mcm21∆::hphMX | SK1 | 7h |
| AM26970 | MATa/alpha REC8-GFP-URA3/REC8-GFP-URA3 PDS1-tdTomato-KITRP1/PDS1-tdTomato-KITRP1 MTW1-tdTomato::NAT/MTW1-tdTomato::NAT ctf19-9A::LEU2/ctf19-9A::LEU2 | SK1 | 4g-h |
| AM27017 | MATa/alpha GAL-NDT80::TRP1/GAL-NDT80::TRP1 ura3::pGPD1-GAL4(848).ER::URA3/ura3::pGPD1-GAL4(848).ER::URA3 MTW1-tdTomato::NAT/MTW1-tdTomato::NAT NDC10-mNeonGreen:LEU2/NDC10-mNeonGreen:LEU2 ame1::pCLB2-3HA-AME1::KanMX6/ame1::pCLB2-3HA-AME1::KanMX6 | SK1 | 4b |
| AM27333 | MATa/alpha GAL-NDT80::TRP1/GAL-NDT80::TRP1 ura3::pGPD1-GAL4(848).ER::URA3/ura3::pGPD1-GAL4(848).ER::URA3 MTW1-tdTomato::NAT/MTW1-tdTomato::NAT NDC10-mNeonGreen:LEU2/NDC10-mNeonGreen:LEU2 clb3::KanMX6::pCUP1-CLB3/clb3::KanMX6::pCUP1-CLB3 mcm21Δ::KANMX6/mcm21Δ::KANMX6 | SK1 | 4c |
| AM27334 | MATa/alpha GAL-NDT80::TRP1/GAL-NDT80::TRP1 ura3::pGPD1-GAL4(848).ER::URA3/ura3::pGPD1-GAL4(848).ER::URA3 MTW1-tdTomato::NAT/MTW1-tdTomato::NAT NDC10-mNeonGreen:LEU2/NDC10-mNeonGreen:LEU2 clb3::KanMX6::pCUP1-CLB3/clb3::KanMX6::pCUP1-CLB3 | SK1 | 4c |
| AM27370 | MATa/alpha GAL-NDT80::TRP1/GAL-NDT80::TRP1 ura3::pGPD1-GAL4(848).ER::URA3/ura3::pGPD1-GAL4(848).ER::URA3 MTW1-tdTomato::NAT/MTW1-tdTomato::NAT NDC10-mNeonGreen:LEU2/NDC10-mNeonGreen:LEU2 dsn1::dsn1(S240D/S250D)-6HIS-3FLAG::URA3/dsn1::dsn1(S240D/S250D)-6HIS-3FLAG::URA3 | SK1 | 7h |
| AM27371 | MATa/alpha GAL-NDT80::TRP1/GAL-NDT80::TRP1 ura3::pGPD1-GAL4(848).ER::URA3/ura3::pGPD1-GAL4(848).ER::URA3 MTW1-tdTomato::NAT/MTW1-tdTomato::NAT NDC10-mNeonGreen:LEU2/NDC10-mNeonGreen:LEU2 okp1::pCLB2-3HA-OKP1::KanMX6/okp1::pCLB2-3HA-OKP1::KanMX6 | SK1 | 4b |
| AM27723 | MATa/alpha GAL-NDT80::TRP1/GAL-NDT80::TRP1 ura3::pGPD1-GAL4(848).ER::URA3/ura3::pGPD1-GAL4(848).ER::URA3 CSE4-mNeonGreen/CSE4-mNeonGreen | SK1 | 6e |
| AM27724 | MATa/alpha GAL-NDT80::TRP1/GAL-NDT80::TRP1 ura3::pGPD1-GAL4(848).ER::URA3/ura3::pGPD1-GAL4(848).ER::URA3 CSE4-mNeonGreen/CSE4-mNeonGreen mcm21∆::hphM/mcm21∆::hphM | SK1 | 6e |
| AM28330 | MATa/alpha irt1::NAT::pCUP-IME1/irt1::NAT::pCUP-IME1 pIME4::NAT::pCUP-IME4/pIME4::NAT::pCUP-IME4 AME1-mNeonGreen:KILEU2/AME1-mNeonGreen:KILEU2 mcm21∆::hphM/mcm21∆::hphM | SK1 | 6h-i |
| AM28331 | MATa/alpha irt1::NAT::pCUP-IME1/irt1::NAT::pCUP-IME1 pIME4::NAT::pCUP-IME4/pIME4::NAT::pCUP-IME4 AME1-mNeonGreen:KILEU2/AME1-mNeonGreen:KILEU2 | SK1 | 6h-i |
| AM28767 | MATa ura3::pADH1-OsTIR1-9MYC::URA3 MTW1-tdTomato::NAT Bir1-AID::kanMX mcm21∆::KanMX6 | w303 | 7b-e, s6c |
| AM28769 | MATa ura3::pADH1-OsTIR1-9MYC::URA3 MTW1-tdTomato::NAT Bir1-AID::kanMX | w303 | 7b-e, s6c |
| AM29121 | MATa/alpha GAL-NDT80::TRP1/GAL-NDT80::TRP1 ura3::pGPD1-GAL4(848).ER::URA3/ura3::pGPD1-GAL4(848).ER::URA3 MTW1-tdTomato::NAT/MTW1-tdTomato::NAT ipl1::KanMX6::pCLB2-3HA-IPL1/ipl1::KanMX6::pCLB2-3HA-IPL1 | SK1 | 7f |
