## Supplemental Table S5 for "The proteomic landscape of centromeric chromatin reveals an essential role for the Ctf19^CCAN^ complex in meiotic kinetochore assembly"

Table S5. Plasmids used in this study

| **Plasmid** | **name** | **description** | **source** |
| --- | --- | --- | --- |
| AM524 | *GAL-GST-SLI15* | pCC1061 from Clarence Chan lab | (Kim et al., 1999) |
| AM747 | *LacI-3FLAG:URA3* | pSB737 from Sue Biggins lab, LacI-*3FLAG:URA3* integrates at *URA3* locus following StuI digestion | (Akiyoshi et al., 2009) |
| AM1103 | <*CEN3-TALO8>:TRP* | pSB964 from Sue Biggins lab, *CEN3* minichromosome | (Akiyoshi et al., 2009) |
| AM1106 | <*CEN3*-TALO8>:TRP* | pSB972 from Sue Biggins lab, CCG -> GCT mutation introduced in CDEIII, *CEN3** minichromosome | (Akiyoshi et al., 2009) |
| AM1278 | pWS082 | sgRNA entry vector | (Shaw et al., 2019) |
| AM1279 | pWS158 | STRONG Cas9 - gRNA gap repair expression vector for budding yeast CRISPR, *URA3* | (Shaw et al., 2019) |
| AM1295 | pWS082_*CSE4*gRNA. | sgRNA entry vector with sgRNA guide for *CSE4* | this study |
| AM1362 | pLC605-*NDC80-3v5-del2-88* | p888 in EU lab, 3v5 C-terminal tagged Ndc80 without the first 2-28 residues | (Chen et al., 2020) |
| AM1604 | pFA6a-*mNeonGreen-KlLEU2* | mNeonGreen tagging plasmid with Kluyveromyces lactis LEU marker | this study |
| AM1467 | pWS082_*NDC80*_gRNA | sgRNA entry vector with sgRNA guide for *NDC80* | this study |
| AM1702 | pWS082_*SLI15*_gRNA_1 | sgRNA entry vector with sgRNA guide for *SLI15* | this study |
| AM1703 | pWS082_*SLI15*_gRNA_2 | sgRNA entry vector with sgRNA guide for *SLI15* | this study |
