## Supplemental Table S6 for "The proteomic landscape of centromeric chromatin reveals an essential role for the Ctf19^CCAN^ complex in meiotic kinetochore assembly"

Table S6. PCR primers used in this study

| **primer** | **sequence** | **description** |
| --- | --- | --- |
| AMo782 | AGATGAAACTCAGGCTACCA | qPCR Forward Primer, chromosome IV arm |
| AMo783 | TGCAACATCGTTAGTTCTTG | qPCR Reverse Primer, chromosome IV arm |
| AMo794 | CCGAGGCTTTCATAGCTTA | qPCR Forward Primer, chromosome IV centromere |
| AMo795 | ACCGGAAGGAAGAATAAGAA | qPCR Reverse Primer, chromosome IV centromere |
| AMo6663 | CAACGATGTGCTTCAGTATTAC | Forward primer to amplify sgRNA from PWS082 derivatives |
| AMo6664 | GCTGTAGATATCCTGCACTC | Reverse primer to amplify sgRNA from PWS082 derivatives |
| AMo6723 | CTGCGTTTATACGTCTCAGTTTTAGAGC | Forward primer to amplify Ca9 vector backbone for CRISPR transformations |
| AMo6724 | GTTTCACTTTCCGTCTCAAGTC | Revese primer to amplify Ca9 vector backbone for CRISPR transformations |
| AMo6819 | GAATGCTGGTCGCTATACTGCTATCTTCCGTTGGCGCAAAC | Forward primer ~1kb upstream of Ndc80 ORF start |
| AMo6846 | GACTTTCGATTTCTAGATTACCTGCT | Forward primer to generate sgRNA to internally tag Cse4 |
| AMo6847 | AAACAGCAGGTAATCTAGAAATCG | Reverse primer to generate sgRNA to internally tag Cse4 |
| AMo6853 | CGTTCGTTCTCCTGCTTAGAGAGC | reverse internal primer to amplify Ndc80 ORF |
| AMo7441 | AAACTGTGATGTAGCACATGTTGAAA | Reverse primer to generate sgRNA to truncate Ndc80 |
| AMo7442 | GACTTTTCAACATGTGCTACATCACA | Forward primer to generate sgRNA to truncate Ndc80 |
| AMo8660 | CGACAAAGAACCTGTTTCCAAGAAGAGGGGAAAGAAGACGTTATGAAAGCTCAAAAAGTGACCTAGATATCGAAACAGACTACGAAGACCAAGCAGGTAATCTAAGAACGCGGCCGCCAG | Forward primer to amplify mNeonGreen with 100 bp homology to CSE4 |
| AMo8661 | GACTTTCTTCGATAACCAAACATGGG | Forward primer to generate sgRNA against SLI15 – sgRNA1 |
| AMo8662 | AAACCCCATGTTTGGTTATCGAAGAA | Reverse primer to generate sgRNA against SLI15 – sgRNA1 |
| AMo8663 | GACTTTCGCAGGAAGGAAGTCACCGA | Forward primer to generate sgRNA against SLI15 – sgRNA2 |
| AMo8664 | AAACTCGGTGACTTCCTTCCTGCGAA | Reverse primer to generate sgRNA against SLI15 – sgRNA2 |
| AMo8665 | AACAAACAAAAACTCGTTTCAAGTATTGCCAATGATACAAACAAAACGTTTCAAAGTATCTCCTCAACTGACGGACTTTGAAAGAGCCCGAAGATTTAAACAATGTCTCAAAGAAGGTCG | Forward forward primer with a 100 bp homology to the sequence upstream of SLI15 start site, and an 18 bp homology to its internal sequence, starting from aa 229 |
| AMo8666 | GCTAGCTTTTTGTGTGTCG | Reverse primer internal to SLI15, approximately 500nt downstream of start site |
| AMo8738 | GAATGAGTTCGCACTGGTGCAGGTACTTCAGTTTCCATTTCAGCTTCTTCTTCATTTTCTGTCTCGATTTCTCGCTTATTTAGAAGTGGCGCGCCTTCCTTGTATAATTCGTCCATACCC | Reverse primer to amplify mNeonGreen with 100 bp homology to CSE4 |
